# Assortative Mating Biases Marker-based Heritability Estimators

**DOI:** 10.1101/2021.03.18.436091

**Authors:** Richard Border, Sean O’Rourke, Teresa de Candia, Michael E. Goddard, Peter M. Visscher, Loic Yengo, Mathew Jones, Matthew C. Keller

## Abstract

Many complex traits are subject to assortative mating (AM), with recent molecular genetic findings confirming longstanding theoretical predictions that AM alters genetic architecture by inducing long range dependence across causal variants. However, all marker-based heritability estimators assume mating is random. We provide mathematical and simulation-based evidence demonstrating that both method-of-moments estimators and likelihood-based estimators produce biased estimates in the presence of AM and that common approaches to account for population structure fail to mitigate this bias. Then, examining height and educational attainment in the UK Biobank, we demonstrate that these biases affect real world traits. Finally, we derive corrected heritability estimators for traits under equilibrium AM.

## Introduction

Primary phenotypic assortative mating (hereafter simply “AM”; the phenomenon whereby mate-choice is based on phenotypic similarity) has been observed for a variety of heritable traits in human and non-human animals [1–4]. A century ago, Fisher demonstrated that AM induces long-range positive correlations between trait-increasing allele counts at causal loci across the genome, thereby increasing genetic variance across successive generations until it approaches a stable equilibrium [5]. Since Fisher’s time, it has been established that many human traits are subject to AM, and that estimates of genetic and environmental variance from twin and family designs, which assume random mating, can be biased in the presence of AM [6]. However, to date there has been no study of how AM influences marker-based heritability estimators. Moreover, many traits that have been focal in the scientific discourse regarding the so-called still-missing heritability—for example, height and educational attainment—are precisely those traits for which both phenotypic and genetic data is consistent with primary phenotypic assortment [4, 7–9], further motivating the need to understand how AM influences marker-based heritability estimates.

Here, we address this gap in knowledge by characterizing the impact of AM on two major families of marker-based heritability estimators: method of moments estimators (MoM; typified by univariate Haseman-Elston (HE) regression [10] but also including PCGC regression and LD score regression [LDSC] [11, 12]), and residual maximum likelihood (REML [13]; typified by GCTA and BOLT-REML [14, 15]). We assume Fisher’s classical model of AM, which describes the equilibrium properties of a heritable trait for which mates’ genotypes are conditionally independent given their phenotypes, and which has formed the theoretical foundation for recent investigations of AM using measured genetic data [4, 8]. We provide mathematical and simulation-based arguments demonstrating that AM induces a modest but nevertheless non-negligible bias in both classes of estimators that is not addressed by conventional methods of accounting for population structure. In the process, we extend results in random matrix theory and classical quantitative genetics by characterizing the higher-order moments of causal variants and the limiting spectral distribution of the genomic relatedness matrix (GRM) under AM, thereby providing intuition with respect to the observation that genomic principal components do not capture the effects of AM. Additionally, we provide empirical results using data from the UK Biobank that are congruent with our theoretical predictions regarding the influence of AM on marker-based heritability estimators. Finally, we provide guidelines for using and interpreting the results of marker-based heritability estimators when applied to traits subject to AM.

## Results

Our theoretical results depend on several key parameters: *r* denotes the phenotypic correlation between mates on a phenotype 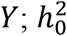 denotes the panmictic heritability, what the heritability of the phenotype would be in the absence of 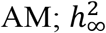 denotes the true equilibrium heritability under AM; and *Z* denotes the *n* × *m* matrix of *n* unrelated individuals’ standardized genotypes at *m* causal loci with effects vector *u*. We initially assume that all causal variants are present in *Z* (i.e., that all the narrow-sense heritability is explained by measured variants); we later relax this condition. The rows of *Z* (individuals’ genotypes) are independent random vectors with *m* × *m* covariance matrix ϒ, which quantifies the correlation between loci. Under random mating, ϒ ≔ ϒ_0_ is approximately block diagonal such that causal variants are largely (aside from linkage disequilibrium between nearby variants) stochastically independent. However, under equilibrium AM, ϒ ≔ ϒ_∞_ is dense due to the presence of positive long-range correlations among trait-increasing allele counts within and across chromosomes. As the elements of ϒ_∞_ agree in sign with the corresponding elements of *uu*^*T*^ (i.e., trait increasing alleles are positively correlated), the equilibrium genetic variance under 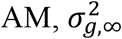, is considerably greater than the panmictic genetic variance 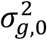. That is, 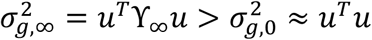 (Figure 1).

**Figure 1.**
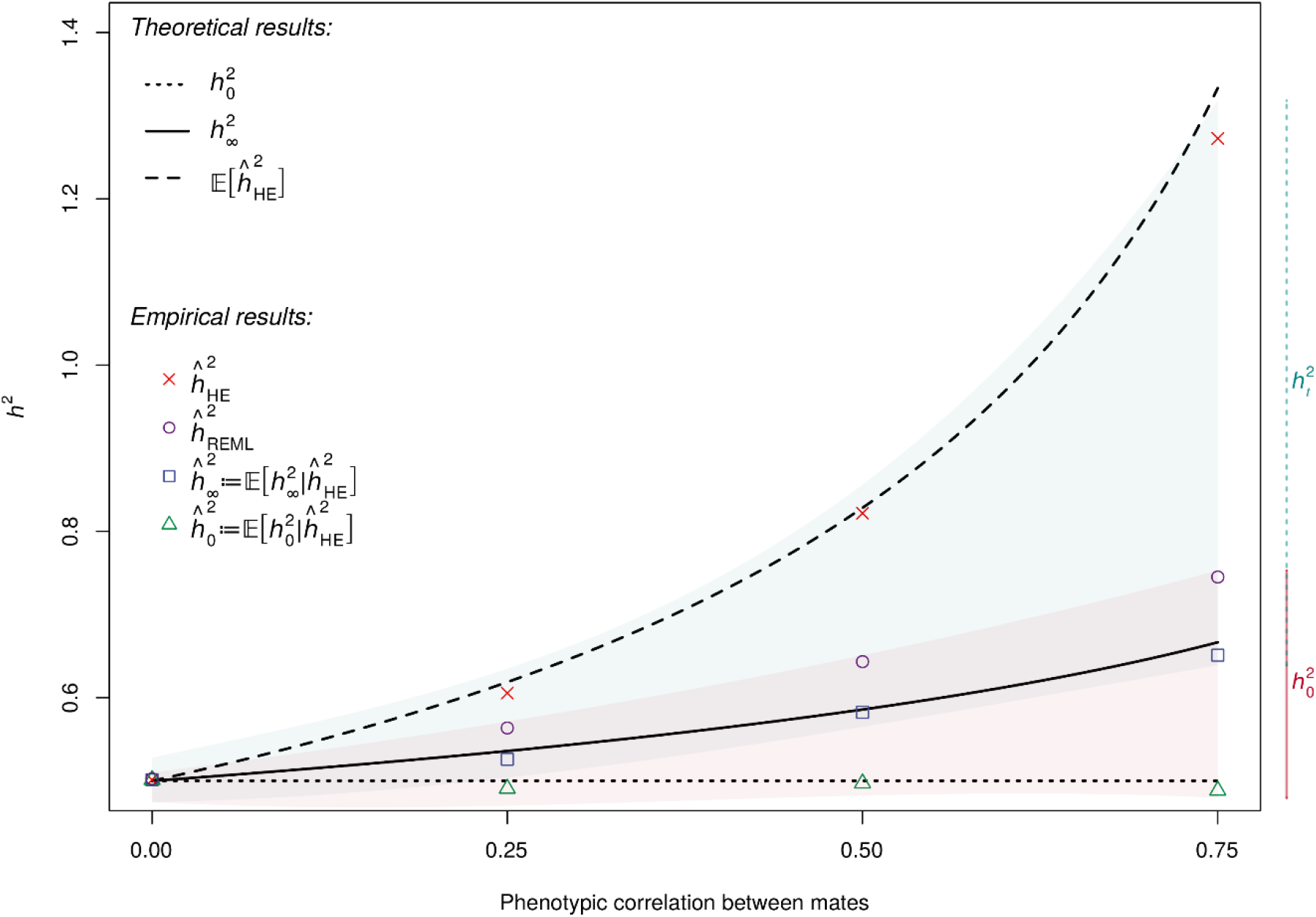
Theoretical and empirical behavior of existing and corrected estimators. Theoretical and empirical behavior of HE regression REML at equilibrium for varying phenotypic correlations among mates (*r*) and fixed panmictic heritability 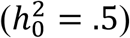 across simulated datasets with sample size fixed at *n* =64,000. HE regression 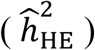 produces upwardly biased estimates consistent with our closed-form approximation under the assumption of exchangeable loci, 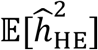. Further, our corrected estimators,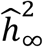 and 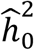, recover the true equilibrium and panmictic heritabilities for a trait at equilibrium. For a trait at disequilibrium, the present day heritability 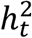 is bounded in expectation between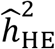 and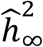 (represented by the teal shaded region), whereas the panmictic heritability 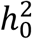 is bounded in expectation between 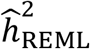 and 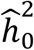 (represented by the red shaded region).

### Haseman-Elston regression estimates under AM

We first derive the influence of AM on heritability estimates from HE regression [10], which is perhaps the simplest MoM marker-based heritability estimator. Let 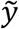 denote the standardized (zero-mean unit-variance) phenotype. The HE regression estimator 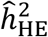 of *h*^*2*^ is the slope of the subdiagonal elements of the phenotypic outer product, 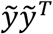, regressed on the subdiagonal elements of the GRM, *m*^*−*1^*ZZ*^*T*^. We demonstrate that 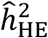 is upwardly biased relative to both 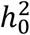 and 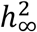 under AM. Intuitively, this is because the phenotypic outer product accurately reflects increases in genetic variance due to positive associations across all pairs trait increasing alleles, whereas the effect of AM on the GRM, the elements of which represent individuals’ average similarity at homologous diploid loci but not across distinct pairs of loci, is negligible. This latter point is itself notable as some studies have erroneously claimed that AM leads to detectable increases in average genomic similarity between mates [17], but the actual increase is trivial (of order *O*(*m*^*−*1^) relative to the increase in genetic variance) and all but undetectable for highly polygenic traits [18]. With respect to the influence of AM on HE regression estimates, and assuming that all the narrow-sense heritability is explained by measured variants, we establish the following general result:

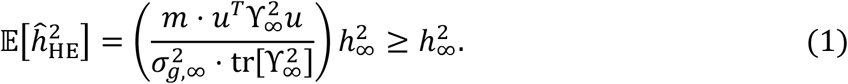

Under AM, the bracketed quantity is greater than one, and thus 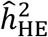 is biased upwards. Under the stricter assumption of exchangeable loci (i.e., each causal variant explains equal variance in the phenotype), we derived the following approximate expression, dependent only on the panmictic heritability and phenotypic correlation between mates:

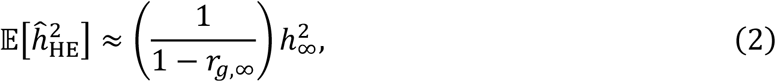

where 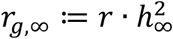 is the equilibrium genetic correlation between mates (Figure 1; see Supp. Materials S3.1 for greater details and proofs). Under exchangeable loci for known *r*, we define estimators of the panmictic and equilibrium heritabilities by applying the following transformations to 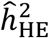 :

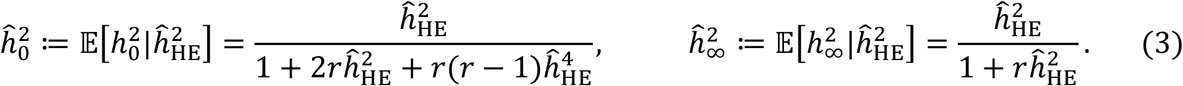

Large scale simulations using realistic genotype data across a variety of scenarios (see Online Methods) demonstrate that the above approximations are accurate even when the exchangeable loci assumption is violated (mean relative error across simulations = -0.009; Figure 1).

In addition, our simulations confirmed that LD score regression [12], which is mathematically equivalent to HE regression when LD scores are exact [19], is analogously biased upwards (Figure 4a). However, the impact of this bias in real world applications depends not only on the extent of AM, but also on the degree to which estimated LD scores reflect the true LD structure in a given population, and therefore no straightforward correction is available.

### Residual maximum likelihood estimates under AM

In contrast to HE regression, for which the upward bias is independent of sample size, we show that the REML estimator, 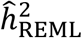, exhibits large upward biases in small samples but converges towards 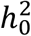 (which is less than the true equilibrium value, 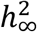) from above in large samples. Formally, under the assumption of exchangeable loci, we prove that

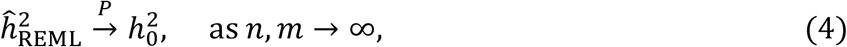

where *n/m* → *c* ∈ (0, ∞); i.e., for polygenic traits, 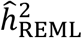 is a consistent estimator of 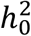. In essence, the parameter values that maximize the residual likelihood function depend only on the eigenvalues of the GRM and the long-range dependence among causal variants induced by AM is “weak” in the sense that the distributions of the eigenvalues (i.e. spectral distributions) of the GRM under random mating and under AM are asymptotically equivalent. However, in *finite* samples, we show via simulation that this convergence can be extremely gradual, requiring samples approaching millions of individuals before estimates reasonably approach 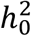(Figure 2a, Figure 2b). Thus, the direction (relative to 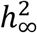) and magnitude of the bias of 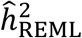 depend on sample size in addition to the panmictic heritability and strength of AM. On the other hand, the number of causal variants, the total number of measured SNPs, and the ratios of these with sample size have no apparent influence on the bias of 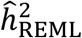 (Supp. Figure S1).

**Figure 2.**
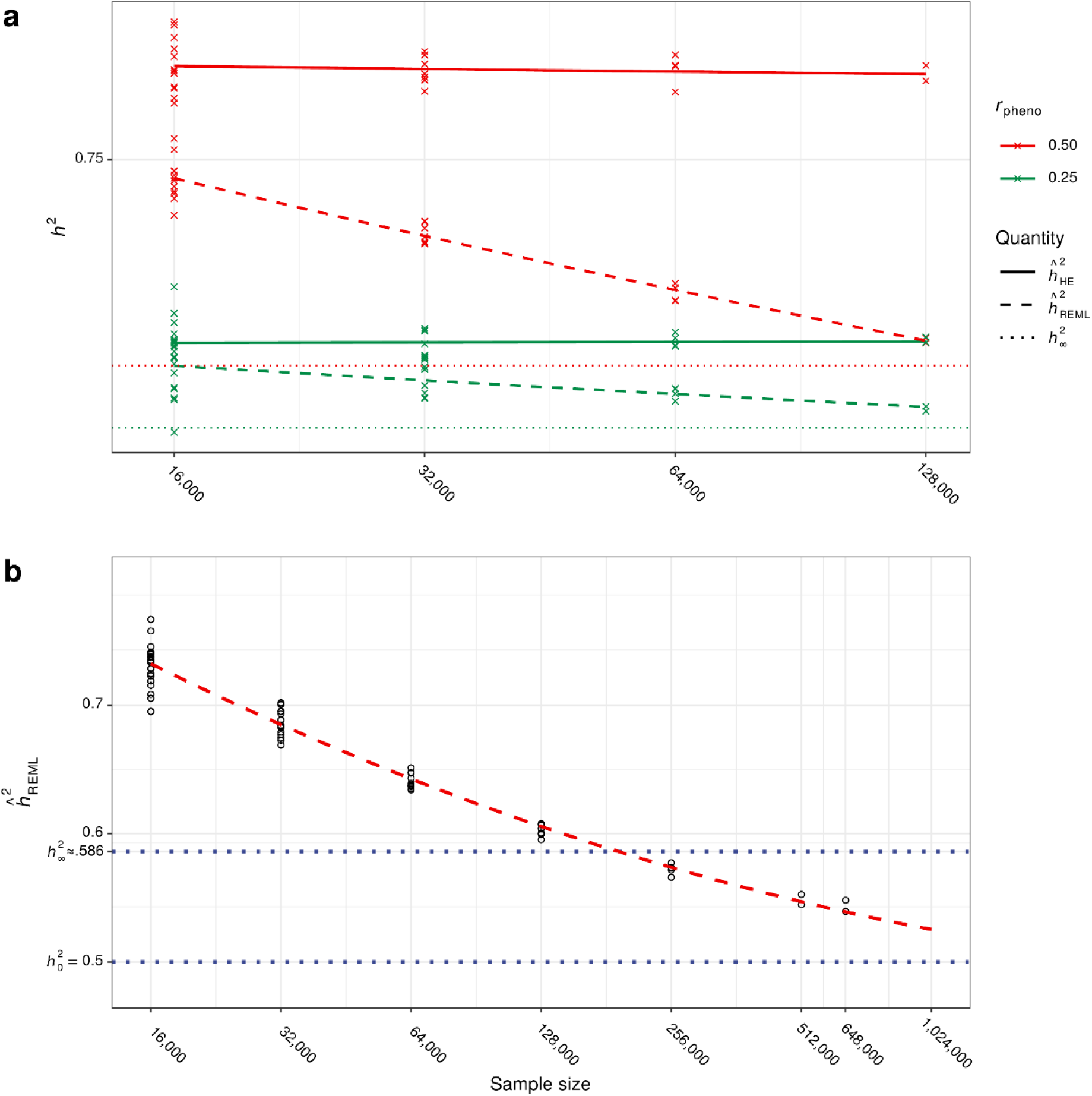
REML and HE estimates across varying sample sizes in simulated data. (a). Comparison of HE regression and REML heritability estimates as functions of sample size for varying phenotypic mating correlation (*r*) and fixed panmictic heritability 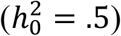 in simulated data. We computed multiple estimates per sample size for each estimator and parameter combination by applying estimators to independent sub-samples. Whereas HE regression estimates are upwardly biased independent of sample size, REML estimates slowly converge to the panmictic heritability as sample sizes increase. (b). Extended simulations demonstrating high-dimensional behavior of the REML estimator as a function of sample size for fixed phenotypic mating correlation (*r* = .5) and panmictic heritability 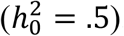. Forward time simulations required a larger population size (*N*_sim_ = 3e6) to obtain samples of up to *n* = 648,000 unrelated individuals. Obtaining REML estimates for samples larger than this was not computationally feasible, but the dashed red line shows predicted values for larger sample sizes extrapolated from a regression model including first and second order log-linear components. Results are consistent with theoretical predictions that the REML estimator converges to the panmictic heritability in very large samples (e.g., >1e6).

### Conventional means of addressing population structure do not mitigate AM-induced bias

Inclusion of ancestral principal components as covariates failed to mitigate the AM-induced bias in both the MoM and the REML estimates (Figure 4a). Indeed, we demonstrate that the AM has a negligible effect on the spectral distribution of the GRM in high-dimensional settings (Supp. Materials S2.4). Similarly, these biases are not mitigated by modeling multiple genetic variance components by partitioning SNPs according to LD score and minor allele frequency (Figure 4a) or by partitioning SNPs by chromosome (results not shown).

### AM-induced bias persists when not all causal variants are measured

In real-world applications, measured genotypes will often include some fraction of the total number of causal variants. To assess the impact of AM when not all variants are measured, we compared HE regression and REML heritability estimates in synthetic data including 100%, 50%, or 25% of both causal and non-causal SNPs by discarding variants at random (Figure 3b). As expected, this resulted in attenuated estimates commensurate with the fraction of missing data. On average, heritability estimates were 8.4% (*se*=0.009%) and 17.6% (*se*=0.009%) lower after randomly discarding 50% and 75% of SNPs, respectively (the degree of attenuation was smaller than the proportion of SNPs dropped due to linkage disequilibrium between retained and discarded SNPs). Nevertheless, while estimates were lower, the pattern of bias due to AM when some of the heritability was missing appeared roughly the same as when no heritability was missing. To test if *ĥ*^2^attenuation varied across methods, we fixed sample size at *n*=128,000 and regressed *ĥ*^2^on the interaction between the fraction of SNPs discarded (0%, 50%, or 75%) and method (HE=1 vs. REML=0). Similarly, to test if *ĥ*^2^attenuation varied with sample size, we regressed *ĥ*^2^on the interaction between the fraction of SNPs discarded and log_10_ sample size. We found that the linear relationship between *ĥ*^2^and the fraction of SNPs discarded did not depend on method 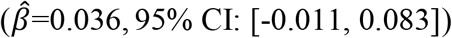 nor on log_10_ sample size 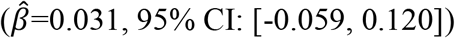.

**Figure 3.**
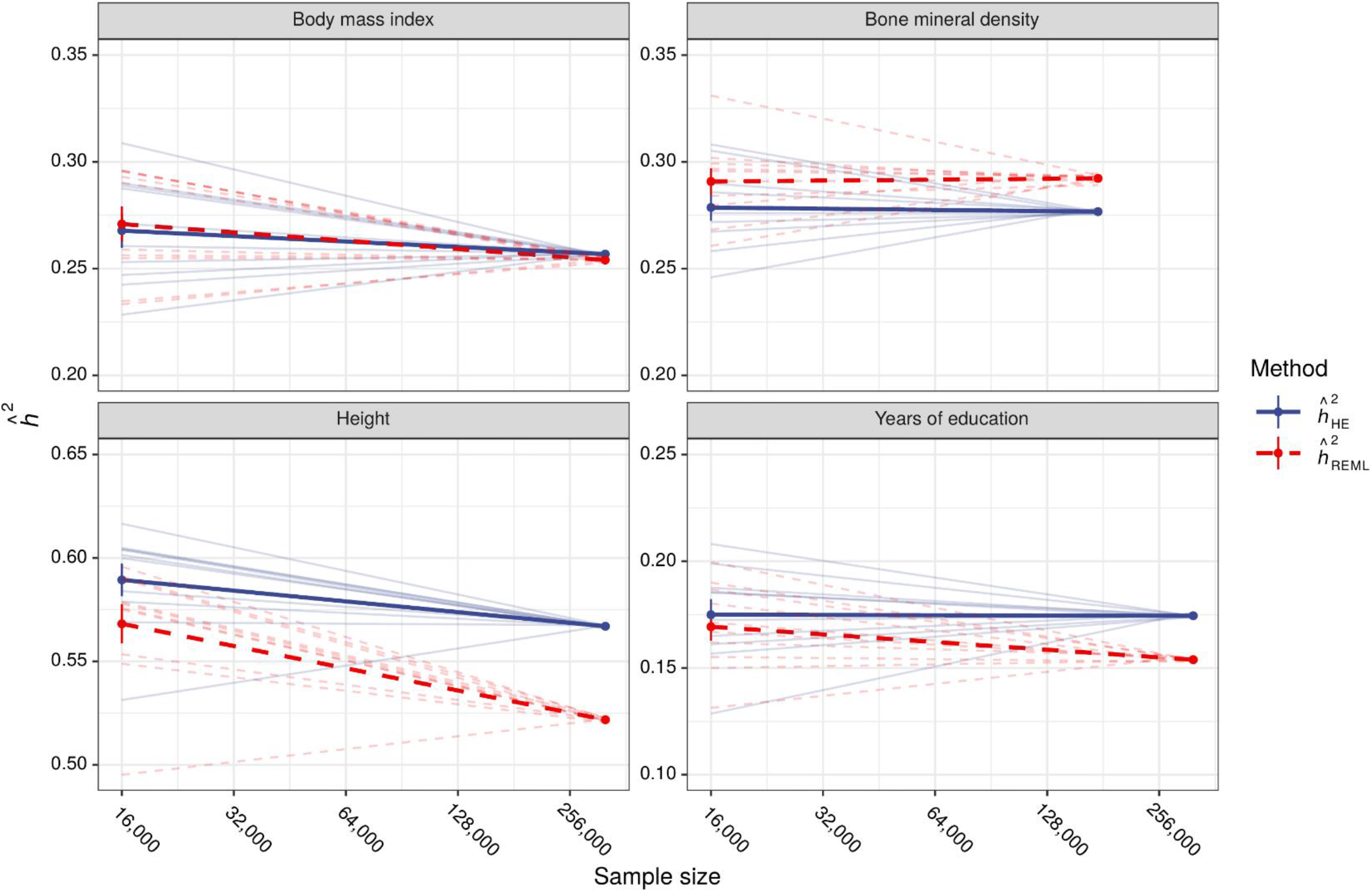
REML and HE estimates across varying sample sizes in UK Biobank data. Comparison of HE regression and REML estimators as a function of sample size for real traits in a sample of unrelated European ancestry UK Biobank participants. Points connected by thin lines represent estimates derived from pairs of complementary disjoint subsamples of size 16,000 and *N −* 16,000, whereas thick lines reflect average log-linear trends. Two negative control traits (body mass index and bone mineral density) and two traits with previous evidence for AM (height and years of education) were selected for analysis *a priori*. Consistent with theoretical predictions, height and years of education demonstrated significant estimator divergence with increasing sample size (*p*=5.24e-4, *p*=3.94e-4, respectively), whereas body mass index and bone mineral density did not (*p*=9.42e-2, *p*=0.302, respectively).

### AM in the UK Biobank

In addition to the experiments using synthetic data described above, we sought to verify our theoretical predictions by examining the relationship between sample size and heritability estimates in a sample 335,551 unrelated European-ancestry individuals in the UK Biobank [20]. We *a priori* selected four phenotypes based on evidence (height, years of education) or lack of evidence (body mass index [BMI], bone mineral density [BMD]) for primary phenotypic AM in a previous study [8]. We then computed HE regression and REML heritability estimates in pairs of small (*n*=16,000) versus large (*n=N*–16,000) non-overlapping subsamples, where *N* depended on the available sample size for each phenotype (Table 1). Congruent with theoretical expectations and simulation results, REML and HE estimates diverged with increasing sample size for height and years of education such that the differences between REML estimates in smaller versus larger subsamples were larger than those of HE estimates (mean difference of differences 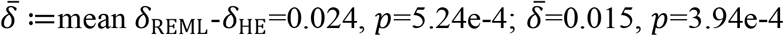; respectively), but not for BMI or BMD 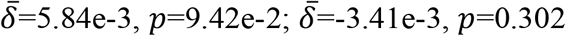; respectively; Figure 3). This was consistent with a previous report that quantified the degree of AM for various traits by correlating polygenic scores between odd-versus even-numbered chromosomes also found that height and educational attainment, but not BMI or BMD, showed signatures of AM [8]. Unlike this previous approach, however, our approach is agnostic to variant direction and effect size. Thus, applying the REML estimator across subsamples of varying sizes provides an alternative and independent way to detect genomic signatures of AM.

**Table 1.** Inflation of heritability estimates for select UK Biobank traits. Spousal correlations (as previously reported in British cohorts [2, 4]) and heritability estimates for height and years of education in the UK Biobank, selected *a priori* on the basis of previous evidence for primary phenotypic AM. Assuming equilibrium, 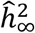 and 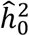 (defined in Equation 3) provide unbiased estimates of the present day and panmictic heritabilites, respectively. Under disequilibrium, they respectively provide probabilistic lower bounds, with 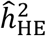 and 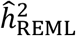 providing complementary upper bounds. The ratio 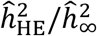 reflects the extent to which HE regression overestimates the true heritability under the assumption of equilibrium.

| Phenotype | $r$ | $n$ | $\hat{h}_{\text{HE}}^2$ [se] | $\hat{h}_{\infty}^2$ [se] | $\hat{h}_{\text{REML}}^2$ [se] | $\hat{h}_0^2$ [se] | $\hat{h}_{\text{HE}}^2/\hat{h}_{\infty}^2$ [se] |
| --- | --- | --- | --- | --- | --- | --- | --- |
| *Height* | 0.240 | 334,798 | 0.567 [4.68e-3] | 0.499 [4.23e-3] | 0.525 [2.22e-3] | 0.467 [3.37e-3] | 1.14 [8.70e-4] |
| *Years of Education* | 0.412 | 332,198 | 0.174 [2.64e-3] | 0.163 [2.46e-3] | 0.155 [2.01e-3] | 0.153 [2.06e-3] | 1.07 [9.46e-4] |

We used previously reported estimates of spousal correlations [2, 4] for height and educational attainment in order to correct HE regression heritability estimates (Table 1) via the estimators in Equation (3). Relative to the corrected equilibrium SNP-heritability estimates, HE regression estimates were inflated by 14% and 7% for height and years of education, respectively.

## Discussion

### Summary of findings

Despite the long-standing understanding that AM alters the genetic architecture of heritable traits, abundant evidence that many phenotypes are subject to AM, and concentrated research activity in marker-based variance component estimation, the effects AM has on these estimators has remained unknown. In the present investigation, we demonstrated that AM biases heritability estimates and that these biases behave differently for MoM estimators versus REML estimators as a function of sample size. In the process, we extended previous results in quantitative genetics and random matrix theory by characterizing the full equilibrium joint distribution of causal variants, demonstrating that the empirical spectral distribution of the resulting relatedness matrix converges to the Marčenko-Pastur law, and thereby proving that REML produces a consistent estimator of the panmictic heritability of polygenic traits in very large samples (see Supp. Materials S2, S3). However, REML estimates of heritability of traits subject to AM behave peculiarly in finite samples, decreasing with larger samples and yielding estimates greater than the true equilibrium heritability in sample sizes typical of those published in the literature (Figure 2b). On the other hand, MoM estimators yield upwardly biased estimates that are higher than REML estimates and remain stable across sample sizes (Figure 2a, Figure 4a). Using UK Biobank data, we observed this differential behavior of estimates for two traits that with previous evidence for AM but not for two negative control traits.

**Figure 4.**
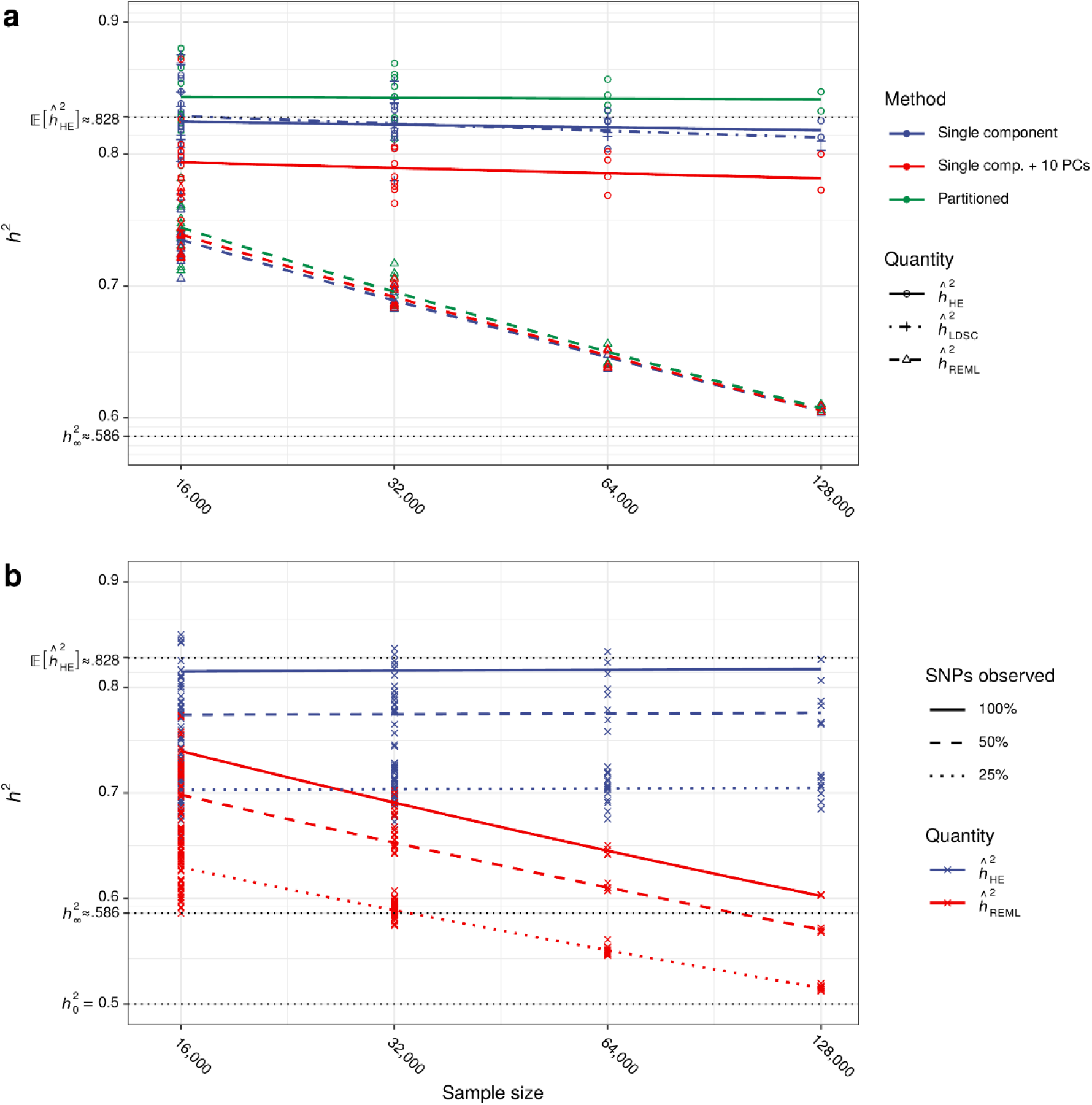
Naïve approaches to addressing AM induced bias and the impact of missing data. (a) Simulations demonstrate that neither partitioned heritability nor principal component adjusted methods mitigate the impact of assortative mating on HE regression and REML estimates. Additionally, simulations confirm that LD score regression (LDSC), which is mathematically equivalent to HE regression, is subject to equivalent biases. “Single component” refers to standard infinitesimal single genomic variance component models, “Single comp. + 10 PCs” included the first ten within-sample principal components as covariates, and “Partitioned” included four annotation-based variance components generated by median splits of within-sample minor allele frequencies and linkage disequilibrium scores. Phenotypic mating correlation and panmictic heritability were fixed (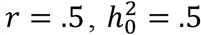, respectively) across simulations. (b) Simulations demonstrate that conclusions regarding estimator bias do not change when only a portion of the genetic variance is explained by measured SNPs. Shown are HE regression and REML estimators when 100%, 50%, or 25% of randomly selected SNPs (both causal and non-causal) were included in the model under the same simulation conditions described in (a). As expected, estimates were attenuated when SNPs were missing but overall patterns remained consistent.

### Implications

Researchers have previously argued that the impact of AM on the heritability of common traits is likely to be “at most modest” [21]. Our results speak to a related but distinct phenomenon: the impact of AM on *heritability estimators* that use molecular genetic data. Notably, this impact is likely to be most salient for “benchmark” traits like height, which has served as a focal point for the discourse surrounding the missing and still-missing heritability phenomena [9, 22]. Our results suggest that the still-missing heritability may be somewhat larger than currently thought for traits subject to AM. For instance, for a trait subject to AM, the REML estimator will produce higher heritability estimates when applied whole-genome sequence data collected from a smaller number of individuals relative to estimates derived from larger samples for whom only sparse array data are available. On the other hand, twin studies are expected to underestimate the total heritability of an additive trait subject to AM [23], implying that the true discrepancy between the total heritability and that which is currently discernible from dense marker data may be greater than previously believed. Our simulations demonstrate that differences in REML heritability estimates derived in samples of 32,000 versus 128,000 individuals are on the order of those induced by randomly omitting 75% of measured SNPs (Figure 2a, Figure 4a). As such, caution is warranted when comparing heritability estimates across methods or across sample sizes for traits subject to AM.

### Interpreting heritability estimates in the presence of AM

As we have demonstrated, existing methods do not produce unbiased heritability estimates for traits subject to primary phenotypic assortment in real world use cases. Although we provide unbiased estimators of both the equilibrium and panmictic heritabilities (by correcting 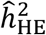 to obtain 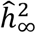 and 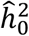 as per Equation [3]), these rely on the strong assumption of equilibrium. Likewise, though REML provides an asymptotically unbiased estimator of the panmictic heritability, the sample sizes required to approach unbiasedness are currently unavailable and computationally impractical. Still, it is possible to derive theoretically-sound bounds on the true values of the present day and panmictic heritabilities using existing methods applied to realistic sample sizes in disequilibrium. Specifically, as a population approaches equilibrium over successive generations of assortative mating, the current heritability at generation 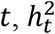, is bounded in expectation from above by 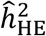, with equality at *t* = 0 (panmixis), and from below by 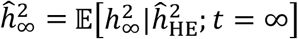, with equality as *t* → ∞ (equilibrium). Likewise, the panmictic heritability 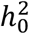 is bounded in expectation from below by 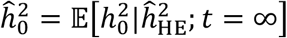, with equality as *t* → ∞, and from above by 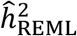(with equality at all generations as *n* → ∞). Thus, as long as the strength of AM isn’t decreasing across generations, 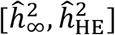 provide probabilistic bounds encompassing the true present day heritability, and 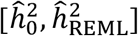 provide probabilistic bounds encompassing the true panmictic heritability under disequilibrium with 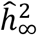 and 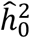 providing conservative point estimates of 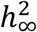 and 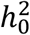, respectively (Figure 1).

### Limitations and future directions

There are several limitations of the current approach. First among these are assumptions inherent to the primary phenotypic AM model. Some of these assumptions, including equilibrium and constancy of the phenotypic mating correlations across generations, provide mathematical tractability and are to some extent inessential to the resulting phenomena. For example, while the problem of characterizing the joint distribution of causal variants becomes substantially more difficult in a population subject to AM that has not reached equilibrium, we observed (results not shown) that estimators behave in a similar, albeit less extreme, fashion relative to their behavior in an equilibrium population. Other assumptions, such as the absence of gene-environment correlation and the conditional independence of mates’ genotypes given their phenotypes (which may be violated in structured populations), are more difficult to evaluate and deserve consideration in future investigations. Additional limitations pertain to our theoretical analysis of the REML estimator, which is rooted in a high-dimensional asymptotics framework. The exact causes of the peculiar behavior of the REML estimator in finite samples, particularly regarding the slow rate of convergence to the panmictic heritability, remain unclear to us. Whether this problem is addressable from an asymptotic perspective or instead requires an alternative, non-asymptotic framework, is an open question and is a target for future theoretical work. Finally, when the phenotypic correlation between mates is known, our results provide a prescription for rectifying AM-induced biases (by adjusting 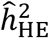 per Equation [3]) under the assumption of equilibrium, which is difficult to verify, and only provide broad bounds on the extent of said biases for populations at disequilibrium. As such, these results should provide motivation and a starting place for the development of new methods that can provide unbiased estimates of genomic variance in the presence of AM.

## Online Methods

### Theoretical framework

#### The primary phenotypic assortment model

Here we introduce the model of AM as proposed by Fisher [5] and further developed by Nagylaki and others [16, 24] (see Supp. Materials S1 for a detailed exposition). Briefly, we consider a phenotype as a random vector composed of independent heritable and non-heritable components:

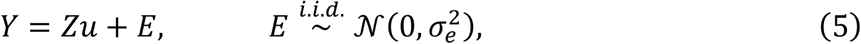

where the rows of *Z*, representing individuals’ standardized genotypes, are independent *m* - dimensional random vectors following a multivariate discrete distribution with finite moments and finite and which we assume are independent under panmixis. The vector of allele substitution effects *u*, which we treat as fixed, is such that 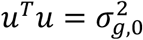. Further, we assume that 1) parent-parent-offspring trios’ phenotypes are jointly Gaussian; 2) the phenotypic correlation between mates, *r* is constant across generations; and 3) there exists *c*_0_ ∈ (0, ∞) such that 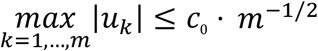; that is, as traits become increasingly polygenic, the maximal variance attributable to individual variants decreases commensurately.

### The equilibrium distribution of causal variants

Over successive generations, the correlation between mates’ phenotypes induces positive correlations across trait increasing allele counts independent of physical position on the genome and thereby increases the total genetic variance of the trait. The genetic variance rapidly approaches a stable equilibrium after several generations (typically within ten generations), at which point the within-individual and cross-mate correlations among causal variants are equal to one another. Using the results of Nagylaki [16], we can express the equilibrium covariance matrix between causal variants as a low rank perturbation of a diagonal matrix of the form: ϒ_∞_ = *D* + *2ϕϕ*^*T*^, where *ϕ* is a known vector-valued function of the substitution effects and mating correlation (Supp. Materials S1.2) with elements *ϕ*_*k*_ = *O*(*m*^*−*1*/2*^) uniformly.

### Higher order moments and the limiting spectral distribution of GRM

Employing tools from the study of thermodynamic equilibria, we extend these classical results to bound moments of higher orders (Supp. Materials S2.1, Supp. Materials S2.4). Using these results, we extend the widely-known Marčenko-Pastur theorem, which describes the limiting distribution of the spectrum of sample covariance matrices corresponding to random matrices with independent sub-Gaussian elements [25], to the case of random matrices with independent rows meeting particular moment conditions (Supp. Materials S2.3). Together, these results establish the limiting spectral distribution of the sample GRM (i.e., the distribution of the eigenvalues of the sample GRM as both sample size and the number of variants examined become large) under AM (Supp. Materials S2.4), providing the necessary theoretical foundation to characterize the asymptotic behavior of the REML estimator. Further, these results explain why controlling for principal components fails to remove AM-induced biases: the impact of AM on the spectrum of the GRM is asymptotically negligible.

### Haseman-Elston regression under AM

The HE regression heritability estimator [26] is obtained by regressing the subdiagonal elements of the standardized phenotypic outer product 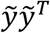 on the subdiagonal elements of the GRM *m*^*−*1^*ZZ*^*T*^. Whereas elements of the outcome (the phenotypic outer product) reflect the dependences among all pairs of causal loci:

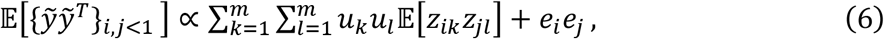

elements of the GRM only capture the dependences among haploid loci that at the same diploid site:

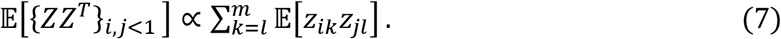

As a result, the variance of the outcome increases whereas the variance of predictor remains largely unaffected, leading to overestimation of the true equilibrium heritability, potentially producing estimates greater than one for strong assortment (Figure 1; see Supp. Materials S3.1 for formal further details and proof). In contrast to the REML estimator, the HE regression estimator is upwardly biased irrespective of sample size (Figure 2a).

### REML and the spectrum of the GRM under AM

The REML estimator [13] models the phenotype as a random vector with marginal distribution,

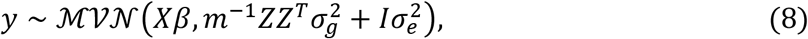

where *X* is an *n* × *c* matrix of covariates with fixed effects *β* and the covariance structure is comprised of a heritable component (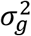 times the GRM) and a non-heritable component (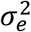 times the identity). The heritability estimator 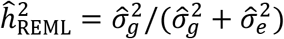 is derived by finding the values of the variance components that satisfy the equation,

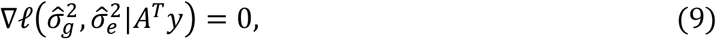

where 𝓁 denotes the marginal log likelihood of the transformed random variable *A*^*T*^*y* for *A*^*T*^: ℝ → (col *X*)^*⊥*^ ⊆ ℝ^*n−c*^, *A*^*T*^*A* = *I*. The conditional expectation of *∇*𝓁 given the genotypes is a function of the eigenvalues of the GRM and, as a result, the asymptotic behavior of 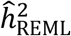 is governed by the asymptotic distribution of the eigenvalues of *m*^*−*1^*ZZ*^*T*^. A foundational result in random matrix theory states that for zero-mean unit-variance sub-Gaussian random matrices *W* ∈ C^*n*×*m*^ with independent elements, the empirical spectral distribution function of *m*^*−*1^*WW*^*T*^ converges almost surely to the Marčenko-Pastur distribution [25]. Employing this result, Jiang and colleagues [27] demonstrated that, in the case of independent causal variants, REML consistently estimates the true heritability in high dimensional settings and is robust to certain forms of model misspecification. In Supp. Materials S2.3, Supp. Materials S2.4, we demonstrate that even though AM induces dependence among causal variants, this dependence is “weak” in the sense that it doesn’t change the limiting spectral distribution of the GRM, thereby allowing us to apply arguments in line with those of Jiang and colleagues’ (Supp. Materials S3.2). Intuitively our result can be summarized as follows: as the sample size and the number of causal variants become large, the eigenvalues of the GRM under AM behave as if the causal variants were independent (as is largely the case under random mating). The behavior of the REML estimator is determined by the behavior of the eigenvalues of the GRM, and thus 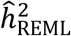 converges to what the heritability would be if the causal variants were independent, i.e., the panmictic heritability.

### Simulation studies

We employed a realistic forward-time simulation framework to generate genotypic and phenotypic data. We then used these data to motivate and verify theoretical results. Below, we describe the general framework and specific simulations we performed.

### Simulation framework

Given a recombination map and *n*_input_ individuals’ phased biallelic genotypes at *p* diploid loci as input, we divided the genome into *k ≪ p* contiguous-within-chromosome, non-overlapping 50kB intervals to obtain a block representation. Recombination events, which occurred with probabilities dictated by the recombination map, were restricted to interval boundaries, thus dramatically reducing the number of haplotypes that had to be tracked while maintaining high genomic resolution. To achieve a target population size *N*_sim_ > *n*_input_, *N*_sim_ pairs of the *n*_input_ individuals were non-monogamously ‘mated’ (i.e., matched and subject to meiosis), resulting in a new generation of *N*_sim_ individuals whose genomes were could be represented in terms of the which of the *n*_input_ haplotype blocks they inherited at each of the *k* intervals. We then repeated this random mating procedure for an additional five generations, resulting in *N*_sim_ chimeric combinations of the original *n*_input_ genotypes while maintaining the linkage disequilibrium structure of the original data. These discretized genotypes comprised the input for the principal AM simulations.

At the beginning of each particular AM simulation with prespecified panmictic heritability 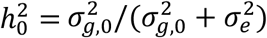, phenotypic correlation between mates *r, p* SNPs, and *m* diploid causal loci *z*_1_, …, *z*_*m*_, *m ≪ p*, the standardized allele substitution effects *u*_1_, …, *u*_*m*_, were independently drawn from a Gaussian distribution with expectation zero and variance 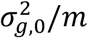. Unless otherwise stated, all simulations used *p* = 10^6^ SNPs. At each generation, phenotypes were constructed via *y* = *Zu* + *e* where *e* was i.i.d. Gaussian with zero expectation and variance 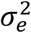. Next, mates were matched according to their respective phenotypes *y*_*i*_, *y*_*j*_ such that corr(*y*_*i*_, *y*_*j*_) ≈ *r*. This was achieved by drawing *N*_sim_ independent doubles 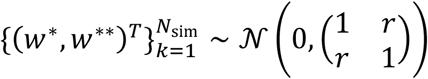 from which *N*_sim_ pairs of indices 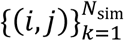 were constructed such that (*i, j*)_*k*_ were the positions of 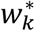 and 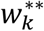 after concatenating and sorting each element of 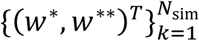. Similarly, *N*_sim_ indices 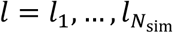 were constructed such that *l*_*k*_ indexed the *k* th largest of the *N*_sim_ simulated phenotypes. Finally, each *k*th mating pair was determined by taking the 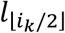 th and 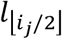 th replicates. Having chosen mates, meiosis occurred as previously detailed to construct the next generations’ genotypes.

### Simulations using UK Biobank data

For each simulation, the input data were derived from phased, imputed genotypes at *p* = 10^6^ randomly selected imputed SNP loci in a sub-sample of *n*_input_ = 435,301 European UK Biobank participants [20]. All SNPs were chosen to meet the following criteria: minor allele frequency greater than 0.01, Hardy-Weinberg *p*-value greater than 10^−6^, INFO score of at least 0.95, and presence on the 1,000 Genomes Phase 3 (1KG3) reference panel [28]. Genotype data were then phased to the 1KG3 reference panel in batches of 40,000 individuals using Eagle v2.4 [29]. This input data was then grown to a population of *N*_sim_ = 10^6^ chimeric genotypes and subjected to an additional five generations of random mating as described in the preceding section.

We conducted AM simulations for varying mating correlations, *r* ∈ {0, .25, .5, .75} and numbers of causal variants, *m* ∈ {10^4^, 10^5^}, with panmictic heritability fixed at 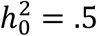. Each simulation consisted of fifteen generations of AM and produced results congruent with classical theory. Prior to heritability estimation, close relatives 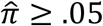, were removed using GCTA v1.93.1 [14], resulting in an average sample size of 141,667 across simulated datasets. Additionally, we ran a limited number of larger, more computationally intensive simulations (*N*_sim_ = 3×10^6^) with mating correlations fixed at *r* = .5 to investigate the large sample behavior of the REML estimator, resulting in at least 648,000 unrelated individuals across simulated datasets. There were no apparent differences across simulations as a function of the number of causal variants or the simulated population size.

### Heritability estimation in simulated data

We split each simulated genotype-phenotype dataset into collections of random subsamples mutually exclusive within collection but not across collections, yielding 16 samples of 16,000 individuals, 8 samples of 16,000, 4 samples of 32,000 individuals, 2 samples of 64,000 individuals, and 1 sample of 128,000 individuals. We then performed HE regression and single-component REML for each subsample (Figure 2a). We used GCTA v1.91.3b [14] to construct genomic related matrices and perform HE regression. We obtained REML heritability estimates using BOLT-LMM v2.3.4 [30] for computational efficiency; though BOLT-LMM uses a randomized algorithm, its numerical accuracy is comparable to that of the exact algorithm implemented GCTA [31].

We also performed a variety of supplementary analyses for a limited set of simulation parameters (*r* = .5, 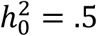, and *m* ∈ {10^4^, 10^5^}, *N* = 10^6^). To demonstrate that including genomic principal components (PCs) as covariates does not mitigate the impact of AM, we included 10 PCs as covariates in the HE regression and REML analyses. For the former, HE regression was conducted in LDAK v5.0 [32], as the HE regression implementation in GCTA cannot accommodate covariates. To demonstrate that the behavior of LD score regression under AM is equivalent to that of HE regression (assuming that the LD scores accurately reflect the LD structure of the sample), we used PLINK v1.9 [33] to obtain GWAS summary statistics and LDSC v1.0.1 [12] to estimate within-sample LD scores using a one centiMorgan sliding window and to perform LD score regression (Figure 4a). To demonstrate that multiple variance component (also known as partitioned approaches [34, 35]) do not mitigate the impact of AM, we fit multicomponent HE regression and REML after partitioning SNPs by minor allele frequency and LD score (Figure 4a).

Finally, to assess the scenario wherein a non-trivial fraction of causal variants aren’t included in the model, we estimated HE regression and REML models after removing 50% or 75% of simulated SNPs at random (Figure 4b).

### Empirical results

#### Sampling procedures

We analyzed 1,211,273 biallelic 1KG3 SNPs with in-sample minor allele frequency greater than 0.01, Hardy-Weinberg *p*-value greater than 10^−6^, and INFO scores of at least 0.95, in a sample of 335,551 unrelated European UK Biobank participants [20]. We selected phenotypes *a priori* on the basis of previous evidence for AM; we chose height (*n* = 335,551) and years of education (*n* = 331,480) as traits with previous evidence of AM, whereas we chose BMI (*n* = 335,551) and BMD (*n* = 191,330) as negative control traits [8]. We measured years of education following the procedures detailed in [36].

#### Analysis/resampling

We tested for evidence of AM by comparing HE and REML heritability estimates in small and large samples. In the presence of AM, our theoretical results imply that HE regression estimates are consistent across all small and large subsamples, whereas REML estimates should decrease with increasing sample size. On the other hand, in the absence of AM, neither HE nor REML estimates should systematically vary with sample size. To this end, we randomly selected ten mutually exclusive subsamples of 16,000 individuals for each trait and compared HE and REML estimates in each subsample to the non-overlapping complementary subsample comprised of the remaining *n −* 16,000 individuals, controlling for sex, age, genotyping batch, testing center, and the first 10 genomic ancestry principal components. To eliminate variance in heritability estimates due to chance differences in covariate effect estimates across subsamples, we adjusted genotypes and phenotypes in the full sample prior to all following analyses. To our knowledge, existing software is incapable of efficient REML analysis using adjusted genotypes (analogous to dosages) in large samples; e.g., BOLT-REML requires hard-calls as input, whereas GCTA and LDAK have cubic complexity in the number of individuals and markers and would require multiple weeks to run on a high thread-count server. We therefore utilized a modified Python implementation of the REML algorithm presented in [31] (code available by request). We used LDAK 5.0 to obtain adjusted HE regression estimates [32]. In order to quantify the divergence of the REML and HE estimators in large versus small samples we performed the following test, analogous to a *t* test of the interaction effect in a 2×2 within-subjects experimental design:

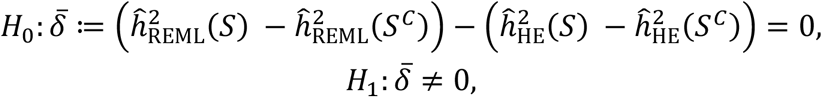

where *S* denotes a given subsample and *S^c^* its complementary subsample. Though this procedure accounts for the dependence among estimates derived in the same subsamples, the individual observations were derived from various partitionings of the same data and do not constitute independent observations. This limitation is not easily avoidable, and the results of this procedure (Figure 3) should be interpreted as descriptive despite our application of inferential procedures.

