## Supplemental Materials for "Assortative Mating Biases Marker-based Heritability Estimators"

March 18, 2021

**A Supplemental figures** **51**

### S1 Assortative mating and the joint distribution of causal variants; previous results

Here we briefly introduce the model of assortative mating (AM) on a quantitative trait first introduced in by Fisher in the early twentieth century (Fisher, 1919/ed) and later developed with greater rigor by a number of quantitative geneticists (Crow *et al.*, 1968; Nagylaki, 1978; Nagylaki, 1982; Karlin *et al.*, 1982; Gimelfarb, 1984). Results, all of which are presented directly in (Nagylaki, 1982) or, in light of a few additional assumptions, are immediate corollaries thereof, are stated without proof.

#### S1.1 Quantitative genetic model and notation

Consider a phenotype $Y$ with zero expectation influenced by $m$ standardized diploid causal SNP loci[1] $\{\mathcal{Z}_k\}_{k=1}^m$,

$$Y = \sum_{k=1}^{m} \mathcal{Z}_k \tilde{u}_k + E,$$

[1]"Standardization" here is with respect to the variance of a diploid biallelic genotype under Hardy-Weinberg equilibrium (HWE). Thus, standardized genotypes at polymorphic loci not in HWE will not have unit variance.

$$E \overset{i.i.d.}{\sim} \mathcal{N}(0, \sigma_e^2).$$

As in (Nagylaki, 1982), we assume that the vector of allele substitution effects $\tilde{u}$ is such that $\sigma_{g,0}^2 = \tilde{u}^T \tilde{u}$, $\sum_k \tilde{u}_k = 0$. We will make the additional assumption that the allelic substitution effects are uniformly bounded in that there exists $0 < c < \infty$ such that, for all $m$, $\max_{k=1,\ldots,m} |\tilde{u}_k| \leq c \cdot m^{-1/2}$. In other words, for increasingly polygenic traits, effect sizes remain commensurately small. At each locus, define the *diploid genic value* as as $Z_k = \mathcal{Z}_k \tilde{u}_k$, $k = 1, \ldots, m$. Each $Z_k$ can be represented as the weighted sum of two *haploid genic values* $Z_k = 2^{-1/2}(H_k^1 + H_k^2) = 2^{-1/2}(\mathcal{H}_k^1 + \mathcal{H}_k^2)\tilde{u}_k$, $k = 1, \ldots, m$. For the sake of cleaner notation, we will sometimes singly index haploid genic values $\{H_k^j\}_{1 \leq k \leq m}^{j \in \{1,2\}}$ as $\{G_k\}_{1 \leq k \leq 2m}$ such that $G_k = H_{\lceil k/2 \rceil}^{1+\bar{k}_2}$ (where $\bar{k}_2$ denotes $k$ mod 2), and proceed analogously for genotypes $\mathcal{G}_k$ such that $\mathcal{G}_k u_k = \mathcal{H}_{\lceil k/2 \rceil}^{1+\bar{k}_2} \tilde{u}_{\lceil k/2 \rceil} := \mathcal{H}_{\lceil k/2 \rceil}^{1+\bar{k}_2} u_k$. Each causal haploid genotype $\mathcal{H}_k$ has been standardized such that $\mathbb{E}[\mathcal{H}_k] = 0$, $Var(\mathcal{H}_k) = 1$. The same is true of each $\mathcal{Z}_k$ when when $\mathcal{H}_k^1$ and $\mathcal{H}_k^2$ are independent as under random mating (i.e., under Hardy-Weinberg equilibrium). Thus, the initial genetic variance of $Y$ is simply

$$Var\left(\sum_{k=1}^{m} \mathcal{Z}_k \tilde{u}_k\right) = \tilde{u}^T \tilde{u} = \sigma_{g,0}^2.$$

We can thus write the phenotype in terms of the genic values or genotypes in multiple ways:

$$\begin{aligned} Y &= \sum_{k=1}^m Z_k + E &&= \sum_{k=1}^m \mathcal{Z}_k \tilde{u}_k + E \\ &= \frac{1}{\sqrt{2}} \sum_{k=1}^{2m} G_k + E &&= \frac{1}{\sqrt{2}} \sum_{k=1}^{2m} \mathcal{G}_k u_k + E \\ &= \frac{1}{\sqrt{2}} \sum_{k=1}^m (H_k^1 + H_k^2) + E &&= \frac{1}{\sqrt{2}} \sum_{k=1}^m (\mathcal{H}_k^1 + \mathcal{H}_k^2)\tilde{u}_k + E. \end{aligned}$$

The haploid genic values follow a multivariate distribution such that, for all $k$,

$$\mathbb{E}[Z_k] = \mathbb{E}[G_k] = \mathbb{E}[H_k^1] = \mathbb{E}[H_k^2] = 0,$$
$$\mathbb{E}[\mathcal{Z}_k] = \mathbb{E}[\mathcal{G}_k] = \mathbb{E}[\mathcal{H}_k^1] = \mathbb{E}[\mathcal{H}_k^2] = 0,$$
$$Var(Z_k)_{t=0} = Var(G_{2k}) = Var(H_k^1) = Var(H_k^2) = \tilde{u}_k^2,$$
$$Var(\mathcal{Z}_k)_{t=0} = Var(\mathcal{G}_k) = Var(\mathcal{H}_k^1) = Var(\mathcal{H}_k^2) = 1,$$

and

$$\mathbb{E}_k[Var(Z_k)_{t=0}] = \mathbb{E}_k[Var(G_k)] = \cdots = \sigma_{g,0}^2/m,$$
$$\mathbb{E}_k[Var(\mathcal{Z}_k)_{t=0}] = \mathbb{E}_k[Var(\mathcal{G}_k)] = \cdots = 1,$$

where $\mathbb{E}_k[\cdot]$ denotes the average over loci (i.e., expectation with respect to the probability measure assigning equal probability to each causal locus). The qualification that the above holds for diploid quantities only at generation zero reflects the fact that the univariate marginal distributions of haploid quantities are time-invariant whereas diploid quantities are subject to departures from Hardy-Weinberg equilibrium.

When all of haploid genic values $\{G_k\}_{k=1}^{2m}$ are independent, as is expected for unlinked loci under random mating, the variance of the phenotype is simply

$$
\begin{aligned}
Var(Y) &= \mathbb{E}\left[\frac{1}{2}\sum_{k=1}^{2m} G_k^2\right] + \sigma_e^2 \\
&= \sigma_{g,0}^2 + \sigma_e^2.
\end{aligned}
$$

On the other hand, if the genic values $\{G_k\}_{k=1}^{2m}$ have non-zero second moments, the total variance is

$$
\begin{aligned}
Var(Y) &= \frac{1}{2}\sum_{k,l=1}^{2m} \mathbb{E}[G_k G_l] + \sigma_e^2 \\
&= \frac{1}{2}\sum_{k,l=1}^{2m} u_k u_l Corr(G_k G_l) + \sigma_e^2.
\end{aligned}
$$

For the special case where the allelic substitution effects $u_i$ and the correlation among haploid genic values are the same for all $i, j, k, l$, $i \neq j \vee k \neq l$ (i.e., for exchangeable loci), $Corr(H_k^i, H_l^j) \equiv \mu$, we have

$$
\begin{aligned}
Var(Y) &= \frac{\mu}{2}\sum_{k,l=1}^{2m} u_k u_l + (1-\mu)\sum_{k=1}^{m} \tilde{u}_k^2 + \sigma_e^2 \\
&= 2\mu\left(\sum_{k=1}^{m} \tilde{u}_m\right)^2 + (1-\mu)\,\sigma_{g,0}^2 + \sigma_e^2 \\
&= \sigma_{g,0}^2\,(1 + (2m-1)\mu) + \sigma_e^2
\end{aligned}
$$

#### S1.2 Causal variant dynamics across generations

Let $Y^*$, $Y^{**}$, and $\tilde{Y}$ and denote the respective phenotypes of parent-parent-offspring trio and similarly denote related quantities. Index generations with $t \in \mathbb{Z}^+$ and define the following parameters:

- $\ell_{kl,t} = Corr(H_{k,t}^1, H_{l,t}^2) = Corr(H_{k,t}^2, H_{l,t}^1)$, $k, l \in \{1, \ldots, m\}$, the correlation between haploid genic values on uniting gametes at generation $t$ (within either parent),
- $\kappa_{kl,t} = Corr(H_{k,t}^1, H_{l,t}^1) = Corr(H_{k,t}^2, H_{l,t}^2)$, $k, l \in \{1, \ldots, m\}$, the correlation between haploid genic values within either parent on the same gamete at generation $t$ (note that for $k = l$, $\kappa_{kl,t} = 1$),
- $\mu_{kl,t} = Corr(H_{k,t}^{i*}, H_{l,t}^{j**}) = Corr(H_{k,t}^{i*}, H_{l,t}^{i**})$ $k, l \in \{1, \ldots, m\}$, $i, j \in \{1, 2\}$, the cross mate correlation between haploid genic values at generation $t$,
- $\Upsilon_t = \{\kappa_{kl,t} + \ell_{kl,t}\}_{kl}$, the $m \times m$ covariance matrix for diploid genotypes $\{\mathcal{Z}_k\}_{k=1}^{m}$ at generation $t$,

- $u_k = \sqrt{Var(G_k)}$, the time-invariant standard deviation of the $k^{th}$ haploid locus; because each $\mathcal{G}_k$ has unit variance, we have $\mathbb{E}_k[u_k^2] = \sigma_{g,0}^2/m$, and $u^T u = 2\sigma_{g,0}^2$; we also denote the $k^{th}$ diploid locus standard deviation by $\tilde{u}_k = u_{\lceil k/2 \rceil}$ with $\tilde{u}^T\tilde{u} = \sigma_{g,0}^2$,
- $\sigma_{g,t}^2 = 2^{-1}Var(\sum_{k=1}^{2m} G_k) = \tilde{u}^T \Upsilon_t \tilde{u}$, the genetic variance at generation $t$ given genotypes,
- $\sigma_e^2 = Var(E)$, the time-invariant residual variance,
- $\sigma_{y,t}^2 = \sigma_{g,t}^2 + \sigma_e^2$, the phenotypic variance at generation $t$,
- $h_t^2 = \sigma_{g,t}^2/\sigma_{y,t}^2$, the heritability at generation $t$,
- $r = Corr(Y^*, Y^{**}) \in (0,1)$, the time-invariant phenotypic correlation between mates,
- $cov_{g,t} = 2^{-1}Cov(\sum_{k=1}^{2m} G_k^*, \sum_{k=1}^{2m} G_k^{**})_t = 2^{-1}\sum_{k,l=1}^{2m} u_k u_l \mu_{\lceil k/2 \rceil \lceil l/2 \rceil, t}$, the genetic covariance between mates at generation $t$,
- $r_{g,t} = cov_{g,t}/\sigma_{g,t}^2$, the genetic correlation between mates at generation $t$. Note that we also have $r_{g,t} = r \cdot h_t^2$, because $corr\left(\sum G_k^*, \sum G_k^{**}\right) = corr\left(\sum G_k^*, Y^*\right) corr\left(Y^*, Y^{**}\right) corr\left(Y^{**}, \sum G_k^{**}\right)$.

At generation zero, we assume that causal variants are unlinked, i.e., that $\kappa_{k \neq l,0} \equiv \ell_{kl,0} \equiv 0$, and at all generations we assume that recombination is equally likely to occur or not to occur between causal loci. Again following (Nagylaki, 1982), in an infinitely large population, a single generation of assortative mating yields the following recurrence for correlations among haploid causal variants:

$$\begin{aligned} \ell_{kl,t} &= \mu_{kl,t-1}, \\ \kappa_{kl,t} &= \frac{1}{2}(\kappa_{kl,t-1} + \ell_{kl,t-1}) \llbracket k \neq l \rrbracket + \llbracket k = l \rrbracket, \end{aligned}$$

such that $\sigma_{g,t}^2 = \sum_{k,l=1}^{m}(\kappa_{kl,t} + \ell_{kl,t})\tilde{u}_k\tilde{u}_l = \tilde{u}^T\Upsilon_t\tilde{u}$. The above system approaches a stable equilibrium such that

$$\begin{aligned} \kappa_{kl,\infty} &= \ell_{kl,\infty} = \mu_{kl,\infty}, \quad \text{for } k \neq l, \\ \ell_{kk,\infty} &= \mu_{kk,\infty}. \end{aligned}$$

At this fixed point, we can write the equilibrium genetic variance in terms of the generation zero genetic variance and the correlations among haploid effects, recalling that $cov_{g,t} = \frac{1}{2}\sum_{kl=1}^{2m} u_k u_l \mu_{\lceil k/2 \rceil \lceil l/2 \rceil, t}$:

$$\sigma_{g,\infty}^2 = \sum_{k,l=1}^{m} (\kappa_{kl,\infty} + \ell_{kl,\infty})\tilde{u}_k\tilde{u}_l$$

$$= \sigma_{g,0}^2 + \tfrac{1}{2} \sum_{k,l=1}^{2m} u_k u_l \mu_{\lceil k/2 \rceil \lceil l/2 \rceil, \infty} - \tfrac{1}{2} \sum_{k=1}^{2m} u_k^2 \mu_{\lceil k/2 \rceil \lceil k/2 \rceil, \infty}$$

$$= \sigma_{g,0}^2 + (1 - m_e^{-1}) cov_{g,\infty}$$

where the effective number of loci is defined as

$$m_e := \left( \sum_{k,l}^{2m} \mu_{\lceil k/2 \rceil \lceil l/2 \rceil, \infty} u_k u_l \right) / \left( \sum_{k=l}^{2m} \mu_{\lceil k/2 \rceil \lceil k/2 \rceil, \infty} u_k^2 \right)$$

Defining $Q = 1 - 1/m_e$ we have that $\sigma_{g,\infty}^2 = \sigma_{g,0}^2 + Q \cdot cov_{g,\infty}$, noting that that $Q \to 1$ as $m_e \to \infty$. We assume that $m_e$ is large and replace $Q$ with the approximation $Q \approx 1$.

Using the relation $\mu_{\lceil k/2 \rceil \lceil l/2 \rceil, \infty} = r \nu_{k,\infty} \nu_{l,\infty}$ derived below, this assumption reduces to the assumption

$$\sum_{k,l}^{2m} \nu_{k,\infty} \nu_{l,\infty} u_k u_l \gg \sum_{k=1}^{2m} \left( \nu_{k,\infty}^2 u_k^2 \right),$$

or, equivalently, that

$$\sum_{k,l}^{2m} \nu_{k,\infty} \nu_{l,\infty} u_k u_l = \nu_\infty^T u u^T \nu_\infty = \left( \sum_{k=1}^{2m} \nu_{k,\infty} u_k \right)^2 \gg \sum_{k=1}^{2m} \left( \nu_{k,\infty}^2 u_k^2 \right).$$

Substituting via $r_{g,t} = cov_{g,t}/\sigma_{g,t}^2$, the equilibrium genetic variance for polygenic traits is approximated as $\sigma_{g,\infty}^2 \approx \sigma_{g,0}^2 (1 - r_{g,\infty})^{-1}$. Substituting $r_{g,\infty} = r h_\infty^2$, the equilibrium heritability is then computed

$$h_\infty^2 \approx \frac{\sigma_{g,0}^2 (1 - r h_\infty^2)^{-1}}{\sigma_{g,0}^2 (1 - r h_\infty^2)^{-1} + \sigma_e^2}.$$

In terms of the time-invariant phenotypic correlations between mates $r = r_{g,\infty}/h_\infty^2$, the equilibrium heritability can be written

$$h_\infty^2 \approx \frac{\sigma_{g,0}^2 (1 - r h_\infty^2)^{-1}}{\sigma_{g,0}^2 (1 - r h_\infty^2)^{-1} + \sigma_e^2}$$

$$\implies h_\infty^2 \approx (2 r \sigma_e^2)^{-1} \left( \sigma_e^2 + \sigma_{g,0}^2 \pm \sqrt{(2 - 4r) \sigma_e^2 \sigma_{g,0}^2 + \sigma_e^4 + \sigma_{g,0}^4} \right)$$

$$= (2 r \sigma_e^2)^{-1} \left( \sigma_e^2 + \sigma_{g,0}^2 \pm \sqrt{\sigma_{y,0}^4 - 4 r \sigma_e^2 \sigma_{g,0}^2} \right).$$

The upper root in this expression can be excluded because it is necessarily greater than or equal to 1, with equality only when $r = 1$. This can be shown from the following observation together with $\sqrt{(1 - h_0^2)^{-2} - 4 r h_0^2 (1 - h_0^2)^{-1}} \geq \left| (1 - h_0^2)^{-1} - 2r \right|$. Observing that $\sigma_{g,0}^2 / \sigma_e^2 = h_0^2 (\sigma_e^2 + \sigma_{g,0}^2)/\sigma_e^2 = h_0^2/(1 -$

$h_0^2$), we can then write the equilibrium heritability in terms of the phenotypic correlation between mates and the generation zero heritability:

$$h_\infty^2 \approx (2r)^{-1}\left((1-h_0^2)^{-1} - \sqrt{(1-h_0^2)^{-2} - 4rh_0^2(1-h_0^2)^{-1}}\right).$$

Similarly, the equilibrium genetic correlation between mates is

$$\begin{aligned} r_{g,\infty} &\approx \frac{r\sigma_{g,0}^2(1-r_{g,\infty})^{-1}}{\sigma_{g,0}^2(1-r_{g,\infty})^{-1}+\sigma_e^2} \\ \implies r_{g,\infty} &= \left(2\sigma_e^2\right)^{-1}\left(\sigma_e^2+\sigma_{g,0}^2 - \sqrt{(2-4r)\,\sigma_e^2\sigma_{g,0}^2+\sigma_e^4+\sigma_{g,0}^4}\right) \\ &= \frac{1}{2}\left((1-h_0^2)^{-1} - \sqrt{(1-h_0^2)^{-2} - 4rh_0^2(1-h_0^2)^{-1}}\right). \end{aligned}$$

Finally, the equilibrium variance is computed

$$\begin{aligned} \sigma_{y,\infty}^2 &\approx \sigma_e^2 + \sigma_{g,0}^2(1-r_{g,\infty})^{-1} \\ &= \left(\frac{1-r_{g,\infty}(1-h_0^2)}{1-r_{g,\infty}}\right)\sigma_{y,0}^2. \end{aligned}$$

Further assumptions are required to make any explicit claims about the equilibrium values of the correlations between haploid causal variants, $\mu_{kl,\infty}$, $k,l=1,\ldots,m$. As in (Nagylaki, 1982), we assume the regressions of individuals' genetic scores on their phenotypes, and on their mates' values, are linear:

$$\mathbb{E}\left[2^{-1/2}\sum_{k=1}^{2m} G_k^*\middle|Y^*\right] = h_t^2 Y^*, \qquad \mathbb{E}\left[2^{-1/2}\sum_{k=1}^{2m} G_k^*\middle|Y^{**}\right] = rh_t^2 Y^{**},$$

and that the same is true of their individual haploid genic values:

$$\mathbb{E}[G_k^*|Y^*] = \nu_{k,t}u_k\sigma_{y,t}^{-1}Y^*, \tag{1}$$

where $\nu_k$ denotes the correlation between the haploid genic value and the phenotype. At equilibrium, we then have

$$\mathbb{E}[G_k^*|Y^{**}] = r\nu_{k,\infty}u_k\sigma_{y,\infty}^{-1}Y^{**}.$$

Thus, by conditional independence of mates' genotypes, we have[2]

[2] We abuse notation here such that $\mu_{kl,\infty}$ refers to $\mu_{\lceil k/2\rceil\lceil l/2\rceil,\infty}$

$$
\begin{aligned}
\mu_{kl,\infty} u_k u_l &= \mathbb{E}[G_k^* G_l^{**}] \\
&= \mathbb{E}\left[\mathbb{E}[G_k^*|Y^{**}]\mathbb{E}[G_l^{**}|Y^{**}]\right] \\
&= r u_k u_l \nu_{k,\infty}\nu_{l,\infty} \\
\implies \mu_{kl,\infty} &= r\nu_{k,\infty}\nu_{l,\infty}.
\end{aligned}
$$

Compute $\nu_{k,\infty}$ as follows:

$$
\begin{aligned}
\nu_{k,\infty}\sigma_{y,\infty} u_k &= \mathbb{E}[G_k Y] \\
&= \sum_{l=1}^{2m} 2^{-1/2} u_k u_l \mathbb{E}[\mathcal{G}_k \mathcal{G}_l] \\
\implies \nu_{k,\infty} &= 2^{-1/2} u_k \sigma_{y,\infty}^{-1}(1 - r\nu_{k,\infty}^2) + 2^{-1/2} r\nu_{k,\infty}\sigma_{y,\infty}^{-1}\sum_{l=1}^{2m}\nu_{l,\infty}u_l.
\end{aligned}
$$

Now, writing

$$
h_\infty^2 Y = \mathbb{E}\left[2^{-1/2}\sum_{l=1}^{2m} G_l \middle| Y\right] = 2^{-1/2}\sigma_{y,\infty}^{-1}\sum_{l=1}^{2m}\nu_{l,\infty}u_l Y,
$$

yields $h_\infty^2 = \sigma_{y,\infty}^{-1} 2^{-1/2}\sum_{l=1}^{2m}\nu_{l,\infty}u_l$, which we then substitute into the previous expression to yield

$$
\nu_{k,\infty} = \sigma_{y,\infty}\left(2^{1/2}u_k r\right)^{-1}\left(\sqrt{\frac{2u_k^2 r}{\sigma_{y,\infty}^2} + (1 - r_{g,\infty})^2} - (1 - r_{g,\infty})\right).
$$

Note that, under exchangeable loci, we then have $\nu_{k,\infty} \equiv \pm\nu_\infty$ for all $k$, which in turn implies that $\mu_{kl,\infty} = \pm r\nu_\infty^2 \equiv \pm\mu_\infty$ for all $k, l$, with $\operatorname{sgn}\mu_{kl,\infty} = \operatorname{sgn} u_k u_l$. That is, the equilibrium correlations between distinct haploid loci are all of equal magnitude, which can be bounded as:

$$
\begin{aligned}
cov_{g,\infty} &= \frac{1}{2}\sum_{k,l=1}^{2m} u_k u_l \mu_\infty \\
&= 4m^2\left(\sigma_{g,0}^2/2m\right)\mu_\infty \\
\implies \mu_\infty &= \pm\frac{r_{g,\infty}\sigma_{g,\infty}^2}{2m\sigma_{g,0}^2} = \mathcal{O}(m^{-1}).
\end{aligned}
$$

In the more general case, for a highly polygenic trait with fixed generation zero conditions, we can employ our assumption that $\max_{k=1,\ldots,m}|\tilde{u}_k| \le c \cdot m^{-1/2}$ to obtain a similar bound. The equilibrium correlations between genic values are $\mu_{kl,\infty} = r\nu_{k,\infty}\nu_{l,\infty}$. Defining $\beta_k = u_k\sigma_{y,\infty}^{-1} = \mathcal{O}(m^{-1/2})$, we compute the equilibrium correlation between mates' haploid genic effects as

$$\begin{aligned}
\mu_{kl,\infty} &= r\nu_{k,\infty}\nu_{l,\infty} \\
&= \sigma^2_{y,\infty}\left(2r^2 u_k u_l\right)^{-1}\left(\sqrt{(1-r_{g,\infty})^2+2u_k^2 r\sigma_{y,\infty}^{-2}}-(1-r_{g,\infty})\right)\left(\sqrt{(1-r_{g,\infty})^2+2u_l^2 r\sigma_{y,\infty}^{-2}}-(1-r_{g,\infty})\right) \\
&= \left(2r^2\beta_k\beta_l\right)^{-1}\left(\sqrt{(1-r_{g,\infty})^2+2\beta_k^2 r}-(1-r_{g,\infty})\right)\left(\sqrt{(1-r_{g,\infty})^2+2\beta_l^2 r}-(1-r_{g,\infty})\right) \qquad (2)\\
&= \tfrac{1}{2}\left(1-r_{g,\infty}\right)^{-2}|\beta_k\beta_l| + \mathcal{o}\left(\beta_k^3,\beta_l^3\right) \\
\implies |\mu_{kl,\infty}| &= \mathcal{O}(m^{-1}).
\end{aligned}$$

The last statement follows from the fact that $(1-r_{g,\infty})^{-2}$ is bounded independent of $m$ via $r_{g,\infty} = rh^2_\infty$ with $r \in (0,1)$, $h^2_\infty \in [0,1]$. In general, we can write the equilibrium covariance between diploid genotypes as

$$\begin{aligned}
\Upsilon_\infty =& \{Cov(\mathcal{Z}_k, \mathcal{Z}_l)\}_{k,l=1}^m \\
=& \begin{pmatrix}
1+r\nu_{1,\infty}^2 & & & \text{Symm} \\
2r\nu_{1,\infty}\nu_{2,\infty} & \ddots & & \\
\vdots & \ddots & \ddots & \\
2r\nu_{1,\infty}\nu_{m,\infty} & \cdots & 2r\nu_{m-1,\infty}\nu_{m,\infty} & 1+r\nu_{m,\infty}^2
\end{pmatrix} \\
:=& D + 2\phi\phi^T,
\end{aligned}$$

where $\phi = (\sqrt{r}\nu_{k,\infty})_{k=1}^m$ and $D = \text{diag}(1-\phi_k^2)_{k=1}^m$.

### S2 Assortative mating and the joint distribution of causal variants; novel results

The results presented in the previous section characterize the first and second moments of causal variants under AM but not moments of higher order. In this section, we employ tools from thermodynamics to directly characterize the equilibrium joint distribution of causal variants and thereby obtain bounds for all higher order moments in terms of the number of causal variants $m$. Then, based on said bounds, we extend previous results in random matrix theory in order the characterize the limiting spectral distribution of a class of covariance matrices generated from random matrices with dependent entries. Finally, we describe the limiting spectral distribution of the genetic relatedness matrix (GRM) under AM.

#### S2.1 A thermodynamic approach

At equilibrium, the variance and covariance of mates' phenotypes are constant across generations. These equilibrium values define the stationary dynamics of assortative mating, and they can be naturally expressed in terms of energy functions suitable for analysis using tools from thermodynamics. This analysis yields a complete characterization of the genetic distribution.

##### S2.1.1 Energy functions and maximum entropy

Begin by defining energy functions

$$
\begin{aligned}
E_1\left(Y^*, Y^{**}\right) &= -Y^{*2} \\
E_2\left(Y^*, Y^{**}\right) &= -Y^{**2} \\
E_3\left(Y^*, Y^{**}\right) &= -Y^* Y^{**}.
\end{aligned} \tag{3}
$$

The intuition from thermodynamics is that states with lower energies are more likely. Thus $E_3$ expresses that mates' phenotype are positively correlated. The signs for $E_1$ and $E_2$ are counterintuitive, but it will be seen below that a countervailing combinatoric effect in going from genotypes to phenotypes leads smaller values of $Y^{*2}$ and $Y^{**2}$ to be more likely, as expected. For now, the signs of these functions can be thought of as purely definitional.

In equilibrium, we have fixed values for the mean energies:

$$
\begin{aligned}
\langle E_1 \rangle &= -\sigma^2_{y,\infty} \\
\langle E_2 \rangle &= -\sigma^2_{y,\infty} \\
\langle E_3 \rangle &= -r\sigma^2_{y,\infty}.
\end{aligned} \tag{4}
$$

Because the dynamics are determined fully at the phenotype level (macroscale), the equilibrium distribution at the genotype level (microscale) will maximize entropy with respect to the macroscale constraints (i.e., (4)). Standard thermodynamic analysis thus implies the genotypes obey a Boltzmann distribution. This enables us to characterize exactly the joint genotypic distribution for mates, and thereby to calculate arbitrary moments and other statistics.

##### S2.1.2 The joint distribution of genotypes under simplifying assumptions

To simplify the initial analysis, take $\sigma_e^2 = 0$, and assume that all SNPs have a minor allele frequency (MAF) of 1/2 and effects $u_i = \sigma_{g,0}/\sqrt{2m}$. Thus $\mathcal{G}_i = \pm 1$ and $G_i = \pm\sigma_{g,0}/\sqrt{2m}$ for all $i$. Denote individuals genetic scores by $\mathscr{G} = \sum_{i=1}^{2m} G_i$, noting that, in this simplified case, $Y = \mathscr{G}$.

The energy functions can be re-expressed as functions of microstates:

$$E_1(\mathcal{G}^*, \mathcal{G}^{**}) = -\sum_{ij=1}^{2m} G_i^* G_j^*.$$

$$E_2(\mathcal{G}^*, \mathcal{G}^{**}) = -\sum_{ij=1}^{2m} G_i^{**} G_j^{**}.$$

$$E_3(\mathcal{G}^*, \mathcal{G}^{**}) = -\sum_{ij=1}^{2m} G_i^* G_j^{**}. \tag{5}$$

Standard thermodynamic analysis then yields the following Boltzmann distribution for mates' genotypes:

$$\begin{aligned}\mathbb{P}[\mathcal{G}^*, \mathcal{G}^{**}] &\propto \exp\left[-\alpha E_1(\mathcal{G}^*, \mathcal{G}^{**}) - \beta E_2(\mathcal{G}^*, \mathcal{G}^{**}) - \gamma E_3(\mathcal{G}^*, \mathcal{G}^{**})\right] \\ &= \exp\left[\alpha \sum_{ij=1}^{2m} G_i^* G_j^* + \beta \sum_{ij=1}^{2m} G_i^{**} G_j^{**} + \gamma \sum_{ij=1}^{2m} G_i^* G_j^{**}\right].\end{aligned} \tag{6}$$

The parameters $\alpha$, $\beta$, and $\gamma$ are inverse temperatures, with values to be determined.

Mathematically, the inverse temperature parameters correspond to Lagrange multipliers for the hard constraints in (4). Typically, these parameters are solved for by substituting the Boltzmann distribution back into the constraint equations (here three constraints for three unknowns). In the present case, there are additional constraints from the assumption of genetic stationarity. Equilibrium at the genetic level implies

$$\mathbb{E}\left[\mathcal{G}_i^* \mathcal{G}_j^*\right] = \mathbb{E}\left[\mathcal{G}_i^* \mathcal{G}_j^{**}\right] = \mathbb{E}\left[\mathcal{G}_i^{**} \mathcal{G}_j^{**}\right] \tag{7}$$

for all $i \neq j$ (Section S1). This symmetry implies $\alpha = \beta$ and $\gamma = 2\beta$. In more detail, the exponent in (6) can be written as $G^T A G$, where $G$ is the concatenation of $G^*$ and $G^{**}$ and the entries of $A$ are all $\alpha$, $\beta$, or $\gamma/2$. (7) implies symmetry under permutation of the components of $G$, which implies all off-diagonal entries of $A$ must be equal[3]. Based on this symmetry argument, the genetic distribution reduces to

$$\mathbb{P}[\mathcal{G}^*, \mathcal{G}^{**}] \propto \exp\left[\beta \left(\sum_{i=1}^{2m} G_i^* + \sum_{i=1}^{2m} G_i^{**}\right)^2\right]. \tag{8}$$

All that remains to specify this distribution is to solve for $\beta$. This is done by requiring the distribution at the phenotype level to satisfy the constraints of (4).

[3]This strong symmetry argument doesn't hold once we allow heterogeneous allelic effects, but symmetry between the components corresponding to $G_i^*$ and $G_i^{**}$, and likewise between $G_j^*$ and $G_j^{**}$, are sufficient to conclude $\alpha = \beta$ and $\gamma = 2\beta$.

##### S2.1.3 The joint distribution of phenotypes under simplifying assumptions

We translate the microstate distribution (8) to a macrostate distribution by coarse-graining. For any value of $Y$, let $n_Y$ be the corresponding number of positive alleles, $|\{i : \mathcal{G}_i = 1\}|$. We have

$$Y = (n_Y - (2m - n_Y)) \frac{\sigma_{g,0}}{\sqrt{2m}} \tag{9}$$

and therefore

$$n_Y = m + \sqrt{\frac{m}{2}} \frac{Y}{\sigma_{g,0}}. \tag{10}$$

The number of genotypes consistent with $Y$ is thus

$$\begin{aligned} \frac{\Gamma(2m+1)}{\Gamma(n_Y+1)\Gamma(2m-n_Y+1)} &\approx \frac{2^{2m}}{\sqrt{\pi m}} \exp\left[-\frac{(n_Y-m)^2}{m}\right] \\ &= \frac{2^{2m}}{\sqrt{\pi m}} \exp\left[-\frac{Y^2}{2\sigma_{g,0}^2}\right]. \end{aligned} \tag{11}$$

The macrostate probabilities can then be obtained by multiplying the microstate probabilities by these counts:

$$\begin{aligned} \mathbb{P}[Y^*, Y^{**}] &= \sum_{\mathcal{G}^*:\mathscr{G}^*=Y^*} \sum_{\mathcal{G}^{**}:\mathscr{G}^{**}=Y^{**}} \mathbb{P}[\mathcal{G}^*, \mathcal{G}^{**}] \\ &\propto \frac{2^{2m}}{\sqrt{\pi m}} \exp\left[-\frac{Y^{*2}}{2\sigma_{g,0}^2}\right] \cdot \frac{2^{2m}}{\sqrt{\pi m}} \exp\left[-\frac{Y^{**2}}{2\sigma_{g,0}^2}\right] \cdot \exp\left[\beta\left(Y^* + Y^{**}\right)^2\right] && (12) \\ &\propto \exp\left[\left(\beta - \frac{1}{2\sigma_{g,0}^2}\right) Y^{*2} + \left(\beta - \frac{1}{2\sigma_{g,0}^2}\right) Y^{**2} + 2\beta Y^* Y^{**}\right]. && (13) \end{aligned}$$

Writing

$$\beta = \frac{r}{2\sigma_{g,0}^2 (1+r)} \tag{14}$$

and

$$\sigma_{Y,\infty}^2 = \frac{\sigma_{g,0}^2}{1-r}, \tag{15}$$

we can re-express the phenotype distribution as

$$\mathbb{P}[Y^*, Y^{**}] \propto \exp\left[-\frac{1}{2\sigma_{Y,\infty}^2 (1-r^2)} \left(Y^{*2} + Y^{**2} - 2rY^*Y^{**}\right)\right], \tag{16}$$

or equivalently

$$(Y^*, Y^{**}) \sim \mathcal{N}\left(0, \sigma^2_{Y,\infty}\begin{bmatrix} 1 & r \\ r & 1 \end{bmatrix}\right). \tag{17}$$

(17) satisfies the constraints of (4). Notice that this analysis also yields an expression for the equilibrium phenotypic variance (15), which was taken as fixed but unknown.[4]

##### S2.1.4 Stationarity

Also useful is to verify that the genetic distribution (Eq. 8) is stationary. Above we used the equilibrium assumption for any two sites, but not for the joint distribution over the full genome. For ease of notation, number loci so that $\{i, m+i\}$ form a diploid pair, and assume without loss of generality that inherited alleles from each parent come from sites 1 through $m$:

$$\tilde{G}_i = \begin{cases} G_i^* & i \leq m \\ G_{i-m}^{**} & i > m. \end{cases} \tag{18}$$

Under these definitions, (8) can be rewritten as

$$\mathbb{P}\left[\mathcal{G}^*, \mathcal{G}^{**}\right] \propto \exp\left[\beta\left(\sum_{i=1}^{2m} \tilde{G}_i + \sum_{i=m+1}^{2m} G_i^* + \sum_{i=m+1}^{2m} G_i^{**}\right)^2\right],$$

and therefore the marginal distribution for the offspring, $\tilde{\mathcal{G}}$, is given by

$$\mathbb{P}\left[\tilde{\mathcal{G}}\right] \propto \sum_{\{\mathcal{G}_i^*\}_{i=m+1}^{2m}} \sum_{\{\mathcal{G}_i^{**}\}_{i=m+1}^{2m}} \exp\left[\beta\left(\sum_{i=1}^{2m} \tilde{G}_i + \sum_{i=m+1}^{2m} G_i^* + \sum_{i=m+1}^{2m} G_i^{**}\right)^2\right].$$

On the other hand, the marginal distribution for one parent is given by

$$\mathbb{P}\left[\mathcal{G}^*\right] \propto \sum_{\{\mathcal{G}_i^{**}\}_{i=1}^{2m}} \exp\left[\beta\left(\sum_{i=1}^{2m} G_i^* + \sum_{i=1}^{m} G_i^{**} + \sum_{i=m+1}^{2m} G_i^{**}\right)^2\right],$$

which is equivalent to the previous expression under the change of variables

$$\left(\{\mathcal{G}_i^*\}_{i=1}^{2m}, \{\mathcal{G}_i^{**}\}_{i=1}^{m}, \{\mathcal{G}_i^{**}\}_{i=m+1}^{2m}\right) \to \left(\left\{\tilde{\mathcal{G}}_i\right\}_{i=1}^{2m}, \{\mathcal{G}_i^*\}_{i=m+1}^{2m}, \{\mathcal{G}_i^{**}\}_{i=m+1}^{2m}\right).$$

[4] Altogether there were four unknowns $(\alpha, \beta, \gamma, \sigma^2_{Y,\infty})$, and four constraints from Eqs. 4 and 7 (where $\mathbb{E}\left[\mathcal{G}_i^*\mathcal{G}_j^*\right] = \mathbb{E}\left[\mathcal{G}_i^{**}\mathcal{G}_j^{**}\right]$ are $\langle E_1\rangle = \langle E_2\rangle$ are redundant).

In conclusion, the joint distribution of mates' genotypes given by (8) (a) reproduces the dynamics of assortative mating as captured in (4) (in particular $cor\left(Y^{*}, Y^{**}\right)=r$), (b) is fully stationary, and (c) maximizes genotype-level entropy conditioned on the constraints of the phenotype-level dynamics. Therefore (8) represents the thermodynamic equilibrium to which the population necessarily converges.

###### S2.1.5 Extension to the general case

The above analysis made three simplifying assumptions: no environmental variance, all allele frequencies are all 1/2, and homogeneous allelic effects (i.e., exchangeable loci). These must be removed to obtain a complete model.

To include environmental variance, the above analysis can be applied to the genetic scores, $\mathscr{G}=\sum_i G_i$, in place of the phenotypes, $Y$. The expression for equilibrium phenotypic variance (15) becomes an expression for equilibrium genetic variance, with the phenotypic correlation ($r$) replaced by the genetic correlation ($r_{g,\infty}$):

$$\sigma_{g,\infty}^{2}=\frac{\sigma_{g,0}^{2}}{1-r_{g,\infty}}.$$

Using the relationship

$$r_{g,\infty}=r\frac{\sigma_{g,\infty}^{2}}{\sigma_{g,\infty}^{2}+\sigma_{e}^{2}},$$

we can derive the same expression for $r_{g,\infty}$ (and hence $\sigma_{g,\infty}^{2}$ and $h_{\infty}^{2}$) as found by previous methods:

$$r_{g,\infty}=\frac{1}{2}\left[\left(1-h_{0}^{2}\right)^{-1}-\sqrt{\left(1-h_{0}^{2}\right)^{-2}-4rh_{0}^{2}\left(1-h_{0}^{2}\right)^{-1}}\right]. \tag{19}$$

Substituting the genetic correlation for the phenotypic correlation in the expression for inverse temperature (14), we obtain

$$\beta=\frac{1+2rh_{0}^{2}-\sqrt{1-4rh_{0}^{2}\left(1-h_{0}^{2}\right)}}{4\sigma_{g,0}^{2}\left(2+rh_{0}^{2}-h_{0}^{2}\right)}. \tag{20}$$

The analysis above did not explicitly constrain allele frequencies, but the symmetry of (8) under sign reversal ($\mathcal{G}\mapsto-\mathcal{G}$) implies all loci have minor allele frequencies (MAF) of 1/2. Arbitrary frequencies values can be imposed by treating them as constraints in the entropy optimization problem, supplementing (4). Writing $p_i$ as the MAF for site $i$ and continuing to code $\mathcal{G}_i$ as $\pm1$ (this is more convenient and conventional in the present framework than standardizing $\mathcal{G}_i$), we have

$$\langle\mathcal{G}_i\rangle=1-2p_i. \tag{21}$$

These constraints manifest as additional terms in the Boltzmann distribution:

$$\mathbb{P}\left[\mathcal{G}^*, \mathcal{G}^{**}\right] \propto \exp\left[\beta\left(\sum_{i=1}^{2m} G_i^* + \sum_{i=1}^{2m} G_i^{**}\right)^2 + \sum_i \lambda_i\left(\mathcal{G}_i^* + \mathcal{G}_i^{**}\right)\right]. \tag{22}$$

The $\lambda_i$ coefficients are determined, together with $\beta$, by solving for the values that lead the distribution to satisfy equations (4) and (21). The system described by (22) is an instance of a spin glass, or generalized Ising model.

Finally, allowing heterogeneous allelic effects, $u_i$, further complicates determination of $\beta$ and $\{\lambda_i\}$. The counting strategy of equations (10) and (11) no longer applies, because the genotypes consistent with a given genetic score no longer arise from a binomial distribution (because of heterogeneous $u_i$), and because they no longer all have equal energy (because of heterogeneous MAF or $\lambda_i$). Nevertheless, the expression for the genetic distribution in (22) is unchanged (only $\beta$ and $\lambda_i$ are changed).

In summary, the complete characterization of the genetic distribution in the general case is given by (22), with $\beta$ and $\{\lambda_i\}$ determined by the constraints in equations (4) and (21). The former constraint amounts to calculating the joint distribution of genetic scores,

$$\mathbb{P}\left[\mathscr{G}^*, \mathscr{G}^{**}\right] \propto \sum_{\mathcal{G}^*: \sum G_i^* = \mathscr{G}^*} \sum_{\mathcal{G}^{**}: \sum G_i^{**} = \mathscr{G}^{**}} \mathbb{P}\left[\mathcal{G}^*, \mathcal{G}^{**}\right],$$

using this distribution to calculate the genetic variance ($\sigma_{g,\infty}^2$) and genetic correlation between mates ($r_{g,\infty}$), and finally calculating the phenotypic correlation between mates:

$$r = \frac{r_{g,\infty}\left(\sigma_{g,\infty}^2 + \sigma_e^2\right)}{\sigma_{g,\infty}^2}.$$

#### S2.2 Moment derivations

Assume a global distribution given by

$$\Pr\left[\mathcal{G}^*, \mathcal{G}^{**}\right] \propto \exp\left[\beta \mathscr{G}^2 + W\right]$$

where $\mathscr{G} = \sum_j \mathcal{G}_j u_j$ and $W = \sum_j \mathcal{G}_j \lambda_j$. Notice that we're concatenating $(\mathcal{G}^*, \mathcal{G}^{**})$ into a single vector $\mathcal{G}$, and similarly for $\mathscr{G}$ and $W$, for economy of notation.

We have shown above that $\beta$ is given by

$$\beta = \frac{1 + 2rh_0^2 - \sqrt{1 - 4rh_0^2\left(1 - h_0^2\right)}}{4\sigma_{g,0}^2\left(2 + rh_0^2 - h_0^2\right)}.$$

The $\lambda$ parameters are unknown and will be approximated below, based on the constraint

$$\mathbb{E}\left[\mathcal{G}_i\right] = q_i.$$

We will make repeated use of the following relation, for various $X$:

$$\begin{aligned}
\frac{\partial}{\partial u_i}\mathbb{E}\left[X\right] &= \frac{\partial}{\partial u_i}\frac{\sum_{\mathcal{G}} X \exp\left[\beta \mathscr{G}^2 + W\right]}{\sum_{\mathcal{G}} \exp\left[\beta \mathscr{G}^2 + W\right]} \\
&= \mathbb{E}\left[\frac{\partial X}{\partial u_i}\right] + \frac{\sum_{\mathcal{G}} X \frac{\partial}{\partial u_i} \exp\left[\beta \mathscr{G}^2 + W\right]}{\sum_{\mathcal{G}} \exp\left[\beta \mathscr{G}^2 + W\right]} - \mathbb{E}\left[X\right] \frac{\sum_{\mathcal{G}} \frac{\partial}{\partial u_i} \exp\left[\beta \mathscr{G}^2 + W\right]}{\sum_{\mathcal{G}} \exp\left[\beta \mathscr{G}^2 + W\right]} \\
&= \mathbb{E}\left[\frac{\partial X}{\partial u_i}\right] + 2\beta \mathbb{E}\left[\mathcal{G}_i \mathscr{G} X\right] - 2\beta \mathbb{E}\left[\mathcal{G}_i \mathscr{G}\right] \mathbb{E}\left[X\right].
\end{aligned}$$

###### S2.2.1 Deriving $\lambda$ from the first moment

Here we expand $\mathbb{E}\left[\mathcal{G}_i\right]$ in $u_i$, holding fixed all other parameters $(u_{-i}, \lambda, \beta)$. We then solve for the value of $\lambda_i$ that yields $q_i$.

Zeroth term:

$$\begin{aligned}
\mathbb{E}\left[\mathcal{G}_i\right]_{u_i=0} &= \frac{\sum_{\mathcal{G}_i} \mathcal{G}_i \exp\left[\mathcal{G}_i \lambda_i\right] \sum_{\mathcal{G}_{-i}} \exp\left[\beta \mathscr{G}_{-i}^2 + W_{-i}\right]}{\sum_{\mathcal{G}_i} \exp\left[\mathcal{G}_i \lambda_i\right] \sum_{\mathcal{G}_{-i}} \exp\left[\beta \mathscr{G}_{-i}^2 + W_{-i}\right]} \\
&= \tanh \lambda_i.
\end{aligned}$$

First derivative:

$$\frac{\partial}{\partial u_i}\mathbb{E}\left[\mathcal{G}_i\right] = 2\beta \mathbb{E}\left[\mathscr{G}\right] - 2\beta \mathbb{E}\left[\mathcal{G}_i \mathscr{G}\right] \mathbb{E}\left[\mathcal{G}_i\right].$$

At $u_i = 0$, $\mathcal{G}_i$ is independent from $\mathcal{G}_{-i}$. Thus the first derivative becomes

$$\frac{\partial}{\partial u_i}_{u_i=0}\mathbb{E}\left[\mathcal{G}_i\right] = 2\beta \left(1 - \tanh^2 \lambda_i\right) \mathbb{E}\left[\mathscr{G}\right]_{u_i=0}.$$

Second derivative:

$$\begin{aligned}
\frac{\partial^2}{\partial u_i^2}\mathbb{E}\left[\mathcal{G}_i\right] &= 2\beta \frac{\mathrm{d}}{\mathrm{d}u_i}\mathbb{E}\left[\mathscr{G}\right] - 2\beta \mathbb{E}\left[\mathcal{G}_i \mathscr{G}\right] \frac{\mathrm{d}}{\mathrm{d}u_i}\mathbb{E}\left[\mathcal{G}_i\right] - 2\beta \mathbb{E}\left[\mathcal{G}_i\right] \frac{\mathrm{d}}{\mathrm{d}u_i}\mathbb{E}\left[\mathcal{G}_i \mathscr{G}\right] \\
&= 4\beta^2 \mathbb{E}\left[\mathcal{G}_i \mathscr{G}^2\right] - 4\beta^2 \mathbb{E}\left[\mathcal{G}_i\right] \mathbb{E}\left[\mathscr{G}^2\right] + 8\beta^2 \mathbb{E}\left[\mathcal{G}_i\right] \mathbb{E}\left[\mathcal{G}_i \mathscr{G}\right]^2 - 8\beta^2 \mathbb{E}\left[\mathcal{G}_i \mathscr{G}\right] \mathbb{E}\left[\mathscr{G}\right].
\end{aligned}$$

At $u_i = 0$ this becomes

$$\frac{\partial^2}{\partial u_i^2}_{u_i=0} \mathbb{E}\left[\mathcal{G}_i\right] = -8\beta^2 \tanh\lambda_i \left(1 - \tanh^2\lambda_i\right) \mathbb{E}\left[\mathscr{G}\right]^2_{u_i=0}.$$

Therefore we have the expansion

$$\mathbb{E}\left[\mathcal{G}_i\right] = \tanh\lambda_i + 2\beta\left(1 - \tanh^2\lambda_i\right)\mathbb{E}\left[\mathscr{G}\right]_{u_i=0} u_i - 4\beta^2 \tanh\lambda_i \left(1 - \tanh^2\lambda_i\right)\mathbb{E}\left[\mathscr{G}\right]^2_{u_i=0} u_i^2 + \mathcal{O}\left(u_i^3\right). \quad (23)$$

Next we expand $\mathbb{E}\left[\mathscr{G}\right]$ in terms of $u_i$:

$$\frac{\partial}{\partial u_i}\mathbb{E}\left[\mathscr{G}\right] = \mathbb{E}\left[\mathcal{G}_i\right] + 2\beta\mathbb{E}\left[\mathcal{G}_i\mathscr{G}^2\right] - 2\beta\mathbb{E}\left[\mathcal{G}_i\mathscr{G}\right]\mathbb{E}\left[\mathscr{G}\right].$$

At $u_i = 0$ this becomes

$$\frac{\partial}{\partial u_i}_{u_i=0}\mathbb{E}\left[\mathscr{G}\right] = \tanh\lambda_i + 2\beta\tanh\lambda_i\left(\mathbb{E}\left[\mathscr{G}^2\right]_{u_i=0} - \mathbb{E}\left[\mathscr{G}\right]^2_{u_i=0}\right).$$

Therefore

$$\mathbb{E}\left[\mathscr{G}\right]_{u_i=0} = \mathbb{E}\left[\mathscr{G}\right] - \tanh\lambda_i u_i - 2\beta\tanh\lambda_i\left(\mathbb{E}\left[\mathscr{G}^2\right] - \mathbb{E}\left[\mathscr{G}\right]^2\right)u_i + \mathcal{O}\left(u_i^2\right) \quad (24)$$

where we have dropped the $u_i = 0$ conditions on the RHS and absorbed the resulting error into the $\mathcal{O}\left(u_i^2\right)$ term.

Substituting (24) into (23) yields

$$\begin{aligned}\mathbb{E}\left[\mathcal{G}_i\right] =& \tanh\lambda_i + 2\beta\left(1 - \tanh^2\lambda_i\right)\mathbb{E}\left[\mathscr{G}\right]u_i - 2\beta\tanh\lambda_i\left(1 - \tanh^2\lambda_i\right)u_i^2 \\ & - 4\beta^2\tanh\lambda_i\left(1 - \tanh^2\lambda_i\right)\mathbb{E}\left[\mathscr{G}^2\right]u_i^2 + \mathcal{O}\left(u_i^3\right).\end{aligned}$$

Writing $\tanh\lambda_i = A + Bu_i + Cu_i^2 + \mathcal{O}\left(u_i^3\right)$ and substituting $q_i$ for $\mathbb{E}\left[\mathcal{G}_i\right]$ yields

$$\begin{aligned}q_i =& \mathcal{O}\left(u_i^3\right) + A + \left(B + 2\beta\left(1 - A^2\right)\mathbb{E}\left[\mathscr{G}\right]\right)u_i \\ & + \left(C - 4\beta AB\mathbb{E}\left[\mathscr{G}\right] - 2\beta A\left(1 - A^2\right) - 4\beta^2 A\left(1 - A^2\right)\mathbb{E}\left[\mathscr{G}^2\right]\right)u_i^2\end{aligned}$$

implying

$$\begin{aligned}A &= q_i \\ B &= -2\beta\left(1 - q_i^2\right)\mathbb{E}\left[\mathscr{G}\right] \\ C &= 2\beta q_i\left(1 - q_i^2\right) + 4\beta^2 q_i\left(1 - q_i^2\right)\left(\mathbb{E}\left[\mathscr{G}^2\right] - 2\mathbb{E}\left[\mathscr{G}\right]^2\right)\end{aligned}$$

and therefore

$$\tanh\lambda_i = q_i - 2\beta\left(1-q_i^2\right)\mathbb{E}\left[\mathscr{G}\right]u_i + 2\beta q_i\left(1-q_i^2\right)u_i^2 + 4\beta^2 q_i\left(1-q_i^2\right)\left(\mathbb{E}\left[\mathscr{G}^2\right] - 2\mathbb{E}\left[\mathscr{G}\right]^2\right)u_i^2 + \mathcal{O}\left(u_i^3\right). \tag{25}$$

###### S2.2.2 Second moments

Here we compute the second order expansion of he second moment in $u_i$ and $u_j$ about zero:

$$\begin{aligned}
\mathbb{E}\left[\mathcal{G}_i\mathcal{G}_j\right]\big|_{u_i,u_j=0} &= \frac{\sum_{\mathcal{G}_i}\mathcal{G}_i\exp\left[\mathcal{G}_i\lambda_i\right]\cdot\sum_{\mathcal{G}_j}\mathcal{G}_j\exp\left[\mathcal{G}_j\lambda_j\right]\cdot\sum_{\mathcal{G}_{-ij}}\exp\left[\beta\mathscr{G}_{-ij}^2 + W_{-ij}\right]}{\sum_{\mathcal{G}_i}\exp\left[\mathcal{G}_i\lambda_i\right]\cdot\sum_{\mathcal{G}_j}\exp\left[\mathcal{G}_j\lambda_j\right]\cdot\sum_{\mathcal{G}_{-ij}}\exp\left[\beta\mathscr{G}_{-ij}^2 + W_{-ij}\right]} \\
&= \tanh\lambda_i\tanh\lambda_j, \\
\frac{\partial}{\partial u_i}\mathbb{E}\left[\mathcal{G}_i\mathcal{G}_j\right]\Big|_{u_i,u_j=0} &= \beta^2\mathbb{E}\left[\mathcal{G}_i\mathcal{G}_j\mathscr{G}^2\right] - 4\beta^2\mathbb{E}\left[\mathcal{G}_i\mathcal{G}_j\right]\mathbb{E}\left[\mathscr{G}^2\right] + 8\beta^2\mathbb{E}\left[\mathcal{G}_i\mathcal{G}_j\right]\mathbb{E}\left[\mathcal{G}_i\mathscr{G}\right]^2 - 8\beta^2\mathbb{E}\left[\mathcal{G}_i\mathscr{G}\right]\mathbb{E}\left[\mathcal{G}_j\mathscr{G}\right]\Big|_{u_i,u_j=0} \\
&= -8\beta^2\tanh\lambda_i\tanh\lambda_j\left(1-\tanh^2\lambda_i\right)\mathbb{E}\left[\mathscr{G}\right]_{u_i,u_j=0}^2, \\
\frac{\partial^2}{\partial u_i\partial u_j}\mathbb{E}\left[\mathcal{G}_i\mathcal{G}_j\right]\Big|_{u_i,u_j=0} &= 2\beta\left(1-\mathbb{E}\left[\mathcal{G}_i\mathcal{G}_j\right]^2\right) + 4\beta^2\mathbb{E}\left[\mathscr{G}^2\right] - 4\beta^2\mathbb{E}\left[\mathcal{G}_i\mathscr{G}\right]^2 - 4\beta^2\mathbb{E}\left[\mathcal{G}_j\mathscr{G}\right]^2 \\
&\quad +8\beta^2\mathbb{E}\left[\mathcal{G}_i\mathscr{G}\right]\mathbb{E}\left[\mathcal{G}_j\mathscr{G}\right]\mathbb{E}\left[\mathcal{G}_i\mathcal{G}_j\right] - 4\beta^2\mathbb{E}\left[\mathcal{G}_i\mathcal{G}_j\right]\mathbb{E}\left[\mathcal{G}_i\mathcal{G}_j\mathscr{G}^2\right]\big|_{u_i,u_j=0} \\
&= 2\beta\left(1-\tanh^2\lambda_i\tanh^2\lambda_j\right) + 4\beta^2\left(1-\tanh^2\lambda_i\tanh^2\lambda_j\right)\mathbb{E}\left[\mathscr{G}^2\right]_{u_i,u_j=0} \\
&\quad + 4\beta^2\left(2\tanh^2\lambda_i\tanh^2\lambda_j - \tanh^2\lambda_i - \tanh^2\lambda_j\right)\mathbb{E}\left[\mathscr{G}\right]_{u_i,u_j=0}^2.
\end{aligned}$$

We now have the following expansion of the second moment:

$$\begin{aligned}
\mathbb{E}\left[\mathcal{G}_i\mathcal{G}_j\right] =& \mathcal{O}\left(u_i^3,u_j^3\right) + \tanh\lambda_i\tanh\lambda_j + 2\beta\tanh\lambda_j\left(1-\tanh^2\lambda_i\right)\mathbb{E}\left[\mathscr{G}\right]_{u_i,u_j=0}u_i \\
&+ 2\beta\tanh\lambda_i\left(1-\tanh^2\lambda_j\right)\mathbb{E}\left[\mathscr{G}\right]_{u_i,u_j=0}u_j - 4\beta^2\tanh\lambda_i\tanh\lambda_j\left(1-\tanh^2\lambda_i\right)\mathbb{E}\left[\mathscr{G}\right]_{u_i,u_j=0}^2u_i^2 \\
&- 4\beta^2\tanh\lambda_i\tanh\lambda_j\left(1-\tanh^2\lambda_j\right)\mathbb{E}\left[\mathscr{G}\right]_{u_i,u_j=0}^2u_j^2 + 2\beta\left(1-\tanh^2\lambda_i\tanh^2\lambda_j\right)u_iu_j \\
&+ 4\beta^2\left(1-\tanh^2\lambda_i\tanh^2\lambda_j\right)\mathbb{E}\left[\mathscr{G}^2\right]_{u_i,u_j=0}u_iu_j \\
&+ 4\beta^2\left(2\tanh^2\lambda_i\tanh^2\lambda_j - \tanh^2\lambda_i - \tanh^2\lambda_j\right)\mathbb{E}\left[\mathscr{G}\right]_{u_i,u_j=0}^2u_iu_j.
\end{aligned} \tag{26}$$

Generalizing the derivation of (24), we have

$$\begin{aligned}
\mathbb{E}\left[\mathscr{G}\right]_{u_i,u_j=0} =& \mathbb{E}\left[\mathscr{G}\right] - \tanh\lambda_iu_i - \tanh\lambda_ju_j - 2\beta\tanh\lambda_i\left(\mathbb{E}\left[\mathscr{G}^2\right] - \mathbb{E}\left[\mathscr{G}\right]^2\right)u_i \\
&- 2\beta\tanh\lambda_j\left(\mathbb{E}\left[\mathscr{G}^2\right] - \mathbb{E}\left[\mathscr{G}\right]^2\right)u_j + \mathcal{O}\left(u_i^2,u_j^2\right).
\end{aligned}$$

Substituting this into (26) yields

$$\begin{aligned}\mathbb{E}\left[\mathcal{G}_i\mathcal{G}_j\right] = {} & \tanh\lambda_i\tanh\lambda_j + 2\beta\tanh\lambda_j\left(1-\tanh^2\lambda_i\right)\mathbb{E}\left[\mathscr{G}\right]u_i + 2\beta\tanh\lambda_i\left(1-\tanh^2\lambda_j\right)\mathbb{E}\left[\mathscr{G}\right]u_j\\ & - 2\beta\tanh\lambda_i\tanh\lambda_j\left(1-\tanh^2\lambda_i\right)u_i^2 - 4\beta^2\tanh\lambda_i\tanh\lambda_j\left(1-\tanh^2\lambda_i\right)\mathbb{E}\left[\mathscr{G}^2\right]u_i^2\\ & - 2\beta\tanh\lambda_i\tanh\lambda_j\left(1-\tanh^2\lambda_j\right)u_j^2 - 4\beta^2\tanh\lambda_i\tanh\lambda_j\left(1-\tanh^2\lambda_j\right)\mathbb{E}\left[\mathscr{G}^2\right]u_j^2\\ & + 2\beta\left(1-\tanh^2\lambda_i\right)\left(1-\tanh^2\lambda_j\right)u_iu_j + 4\beta^2\left(1-\tanh^2\lambda_i\right)\left(1-\tanh^2\lambda_j\right)\mathbb{E}\left[\mathscr{G}^2\right]u_iu_j + \mathcal{O}\left(u_i^3,u_j^3\right).\end{aligned}$$

Finally, substituting (25) yields

$$\mathbb{E}\left[\mathcal{G}_i\mathcal{G}_j\right] = q_iq_j + \left(2\beta + 4\beta^2\operatorname{var}\left(\mathscr{G}\right)\right)\left(1-q_i^2\right)\left(1-q_j^2\right)u_iu_j + \mathcal{O}\left(u_i^3,u_j^3\right). \tag{27}$$

This can also be written

$$\operatorname{corr}\left(\mathcal{G}_i,\mathcal{G}_j\right) = \alpha_i\alpha_j$$

with

$$\alpha_i = \sqrt{\left(2\beta + 4\beta^2\operatorname{var}\left(\mathscr{G}\right)\right)\left(1-q_i^2\right)}u_i + \mathcal{O}\left(u_i^2\right).$$

###### S2.2.3 Third moments

**Three distinct indices ($\mathbb{E}[\mathcal{G}_i\mathcal{G}_j\mathcal{G}_k]$)** We compute the second order expansion of $\mathbb{E}\left[\mathcal{G}_i\mathcal{G}_j\mathcal{G}_k\right]$ in $u_i, u_j, u_k$ about 0:

Zeroth term:

$$\begin{aligned}\left.\mathbb{E}\left[\mathcal{G}_i\mathcal{G}_j\mathcal{G}_k\right]\right|_{u_i,u_j,u_k=0} &= \tanh\lambda_i\tanh\lambda_j\tanh\lambda_k,\\ \left.\frac{\partial}{\partial u_i}\mathbb{E}\left[\mathcal{G}_i\mathcal{G}_j\mathcal{G}_k\right]\right|_{u_i,u_j,u_k=0} &= 2\beta\mathbb{E}\left[\mathcal{G}_j\mathcal{G}_k\mathscr{G}\right] - 2\beta\mathbb{E}\left[\mathcal{G}_i\mathscr{G}\right]\mathbb{E}\left[\mathcal{G}_i\mathcal{G}_j\mathcal{G}_k\right]\Big|_{u_i,u_j,u_k=0}\\ &= 2\beta\tanh\lambda_j\tanh\lambda_k\left(1-\tanh^2\lambda_i\right)\mathbb{E}\left[\mathscr{G}\right]\Big|_{u_i,u_j,u_k=0},\\ \left.\frac{\partial^2}{\partial u_i^2}\mathbb{E}\left[\mathcal{G}_i\mathcal{G}_j\mathcal{G}_k\right]\right|_{u_i,u_j,u_k=0} &= 4\beta^2\Big(\mathbb{E}\left[\mathcal{G}_i\mathcal{G}_j\mathcal{G}_k\mathscr{G}^2\right] - \mathbb{E}\left[\mathcal{G}_i\mathcal{G}_j\mathcal{G}_k\right]\mathbb{E}\left[\mathscr{G}^2\right]\\ &\quad\left. + 2\mathbb{E}\left[\mathcal{G}_i\mathcal{G}_j\mathcal{G}_k\right]\mathbb{E}\left[\mathcal{G}_i\mathscr{G}\right]^2 - 2\mathbb{E}\left[\mathcal{G}_i\mathscr{G}\right]\mathbb{E}\left[\mathcal{G}_j\mathcal{G}_k\mathscr{G}\right]\Big)\right|_{u_i,u_j,u_k=0}\\ &= -8\beta^2\tanh\lambda_i\tanh\lambda_j\tanh\lambda_k\left(1-\tanh^2\lambda_i\right)\left.\mathbb{E}\left[\mathscr{G}\right]^2\right|_{u_i,u_j,u_k=0},\\ \left.\frac{\partial^2}{\partial u_i\partial u_j}\mathbb{E}\left[\mathcal{G}_i\mathcal{G}_j\mathcal{G}_k\right]\right|_{u_i,u_j,u_k=0} &= 2\beta\tanh\lambda_k\left(1-\tanh^2\lambda_i\tanh^2\lambda_j\right) + 4\beta^2\tanh\lambda_k\left(1-\tanh^2\lambda_i\tanh^2\lambda_j\right)\mathbb{E}\left[\mathscr{G}^2\right]\\ &\quad + 4\beta^2\tanh\lambda_k\left(2\tanh^2\lambda_i\tanh^2\lambda_j - \tanh^2\lambda_i - \tanh^2\lambda_j\right)\left.\mathbb{E}\left[\mathscr{G}\right]^2\right|_{u_i,u_j,u_k=0}.\end{aligned}$$

We now have the following expansion of the third moment:

$$\begin{aligned}\mathbb{E}\left[\mathcal{G}_i\mathcal{G}_j\mathcal{G}_k\right] = &\mathcal{O}\left(u_i^3, u_j^3, u_k^3\right) + \tanh\lambda_i \tanh\lambda_j \tanh\lambda_k \\ &+ 2\beta \tanh\lambda_j \tanh\lambda_k \left(1-\tanh^2\lambda_i\right) \mathbb{E}\left[\mathscr{G}\right]_{u_i,u_j,u_k=0} u_i \\ &+ 2\beta \tanh\lambda_i \tanh\lambda_k \left(1-\tanh^2\lambda_j\right) \mathbb{E}\left[\mathscr{G}\right]_{u_i,u_j,u_k=0} u_j \\ &+ 2\beta \tanh\lambda_i \tanh\lambda_j \left(1-\tanh^2\lambda_k\right) \mathbb{E}\left[\mathscr{G}\right]_{u_i,u_j,u_k=0} u_k \\ &- 4\beta^2 \tanh\lambda_i \tanh\lambda_j \tanh\lambda_k \left(1-\tanh^2\lambda_i\right) \mathbb{E}\left[\mathscr{G}\right]^2_{u_i,u_j,u_k=0} u_i^2 \\ &- 4\beta^2 \tanh\lambda_i \tanh\lambda_j \tanh\lambda_k \left(1-\tanh^2\lambda_j\right) \mathbb{E}\left[\mathscr{G}\right]^2_{u_i,u_j,u_k=0} u_j^2 \\ &- 4\beta^2 \tanh\lambda_i \tanh\lambda_j \tanh\lambda_k \left(1-\tanh^2\lambda_k\right) \mathbb{E}\left[\mathscr{G}\right]^2_{u_i,u_j,u_k=0} u_k^2 \\ &+ 2\beta \tanh\lambda_k \left(1-\tanh^2\lambda_i \tanh^2\lambda_j\right) u_i u_j \\ &+ 4\beta^2 \tanh\lambda_k \left(1-\tanh^2\lambda_i \tanh^2\lambda_j\right) \mathbb{E}\left[\mathscr{G}^2\right]_{u_i,u_j,u_k=0} u_i u_j \\ &+ 4\beta^2 \tanh\lambda_k \left(2\tanh^2\lambda_i \tanh^2\lambda_j - \tanh^2\lambda_i - \tanh^2\lambda_j\right) \mathbb{E}\left[\mathscr{G}\right]^2_{u_i,u_j,u_k=0} u_i u_j \\ &+ 2\beta \tanh\lambda_j \left(1-\tanh^2\lambda_i \tanh^2\lambda_k\right) u_i u_k \\ &+ 4\beta^2 \tanh\lambda_j \left(1-\tanh^2\lambda_i \tanh^2\lambda_k\right) \mathbb{E}\left[\mathscr{G}^2\right]_{u_i,u_j,u_k=0} u_i u_k \\ &+ 4\beta^2 \tanh\lambda_j \left(2\tanh^2\lambda_i \tanh^2\lambda_k - \tanh^2\lambda_i - \tanh^2\lambda_k\right) \mathbb{E}\left[\mathscr{G}\right]^2_{u_i,u_j,u_k=0} u_i u_k \\ &+ 2\beta \tanh\lambda_i \left(1-\tanh^2\lambda_j \tanh^2\lambda_k\right) u_j u_k \\ &+ 4\beta^2 \tanh\lambda_i \left(1-\tanh^2\lambda_j \tanh^2\lambda_k\right) \mathbb{E}\left[\mathscr{G}^2\right]_{u_i,u_j,u_k=0} u_j u_k \\ &+ 4\beta^2 \tanh\lambda_i \left(2\tanh^2\lambda_j \tanh^2\lambda_k - \tanh^2\lambda_j - \tanh^2\lambda_k\right) \mathbb{E}\left[\mathscr{G}\right]^2_{u_i,u_j,u_k=0} u_j u_k. \end{aligned} \tag{28}$$

Generalizing the derivation of (24), we have

$$\mathbb{E}\left[\mathscr{G}\right]_{u_i,u_j,u_k=0} = \mathbb{E}\left[\mathscr{G}\right] - \left(1+2\beta\left(\mathbb{E}\left[\mathscr{G}^2\right]-\mathbb{E}\left[\mathscr{G}\right]^2\right)\right)\left(\tanh\lambda_i u_i + \tanh\lambda_j u_j + \tanh\lambda_k u_k\right) + \mathcal{O}\left(u_i^2, u_j^2, u_k^2\right).$$

Substituting this into (28) yields

$$\begin{aligned}\mathbb{E}\left[\mathcal{G}_i\mathcal{G}_j\mathcal{G}_k\right] = &\tanh\lambda_i \tanh\lambda_j \tanh\lambda_k \\ &+ 2\beta \tanh\lambda_j \tanh\lambda_k \left(1-\tanh^2\lambda_i\right) \mathbb{E}\left[\mathscr{G}\right] u_i \\ &+ 2\beta \tanh\lambda_i \tanh\lambda_k \left(1-\tanh^2\lambda_j\right) \mathbb{E}\left[\mathscr{G}\right] u_j \\ &+ 2\beta \tanh\lambda_i \tanh\lambda_j \left(1-\tanh^2\lambda_k\right) \mathbb{E}\left[\mathscr{G}\right] u_k \\ &- 2\beta \tanh\lambda_i \tanh\lambda_j \tanh\lambda_k \left(1-\tanh^2\lambda_i\right)\left(1+2\beta\mathbb{E}\left[\mathscr{G}^2\right]\right) u_i^2 \\ &- 2\beta \tanh\lambda_i \tanh\lambda_j \tanh\lambda_k \left(1-\tanh^2\lambda_j\right)\left(1+2\beta\mathbb{E}\left[\mathscr{G}^2\right]\right) u_j^2 \\ &- 2\beta \tanh\lambda_i \tanh\lambda_j \tanh\lambda_k \left(1-\tanh^2\lambda_k\right)\left(1+2\beta\mathbb{E}\left[\mathscr{G}^2\right]\right) u_k^2 \end{aligned}$$

$$
\begin{aligned}
&+ 2\beta \tanh \lambda_k \left(1 - \tanh^2 \lambda_i\right) \left(1 - \tanh^2 \lambda_j\right) \left(1 + 2\beta \mathbb{E}\left[\mathscr{G}^2\right]\right) u_i u_j \\
&+ 2\beta \tanh \lambda_j \left(1 - \tanh^2 \lambda_i\right) \left(1 - \tanh^2 \lambda_k\right) \left(1 + 2\beta \mathbb{E}\left[\mathscr{G}^2\right]\right) u_i u_k \\
&+ 2\beta \tanh \lambda_i \left(1 - \tanh^2 \lambda_j\right) \left(1 - \tanh^2 \lambda_k\right) \left(1 + 2\beta \mathbb{E}\left[\mathscr{G}^2\right]\right) u_j u_k \\
&+ \mathcal{O}\left(u_i^3, u_j^3, u_k^3\right).
\end{aligned}
$$

Substituting (25) yields

$$
\begin{aligned}
\mathbb{E}\left[\mathcal{G}_i \mathcal{G}_j \mathcal{G}_k\right] =& \mathcal{O}\left(u_i^3, u_j^3, u_k^3\right) + q_i q_j q_k + q_k \left(1 - q_i^2\right) \left(1 - q_j^2\right) \left(2\beta + 4\beta^2 \mathrm{var}\left(\mathscr{G}\right)\right) u_i u_j \\
&+ q_j \left(1 - q_i^2\right) \left(1 - q_k^2\right) \left(2\beta + 4\beta^2 \mathrm{var}\left(\mathscr{G}\right)\right) u_i u_k + q_i \left(1 - q_j^2\right) \left(1 - q_k^2\right) \left(2\beta + 4\beta^2 \mathrm{var}\left(\mathscr{G}\right)\right) u_j u_k.
\end{aligned} \tag{29}
$$

Notice the nontrivial terms match the expression for the second moment from (27). Thus the third central moment is zero to the second degree in $u$:

$$
\begin{aligned}
\mathbb{E}\left[\left(\mathcal{G}_i - q_i\right)\left(\mathcal{G}_j - q_j\right)\left(\mathcal{G}_k - q_k\right)\right] &= \mathbb{E}\left[\mathcal{G}_i \mathcal{G}_j \mathcal{G}_k\right] - q_i \mathbb{E}\left[\mathcal{G}_j \mathcal{G}_k\right] - q_j \mathbb{E}\left[\mathcal{G}_i \mathcal{G}_k\right] - q_k \mathbb{E}\left[\mathcal{G}_i \mathcal{G}_j\right] + 2 q_i q_j q_k \\
&= \mathcal{O}\left(u_i^3, u_j^3, u_k^3\right).
\end{aligned} \tag{30}
$$

For all values of $u_i, u_j, u_k$, the distribution of $\mathcal{G}_i, \mathcal{G}_j, \mathcal{G}_k$ is fully determined by the distribution of $\mathscr{G}_{-i,j,k}$, independent of $m$ (Eq. (6)). Assuming the higher moments of $\mathscr{G}_{-i,j,k}$ do not grow with $m$, an assumption we make going forward, we then have

$$
\mathbb{E}\left[\left(\mathcal{G}_i - q_i\right)\left(\mathcal{G}_j - q_j\right)\left(\mathcal{G}_k - q_k\right)\right] = \mathcal{O}\left(m^{-3/2}\right).
$$

**Two distinct indices ($\mathbb{E}[\mathcal{G}_i^2 \mathcal{G}_j]$)** Let $i, j$ be distinct indices. Using $\mathcal{G}_i^2 = 1$ and substituting (27) yields

$$
\begin{aligned}
\mathbb{E}\left[\left(\mathcal{G}_i - q_i\right)^2 \left(\mathcal{G}_j - q_j\right)\right] &= \mathbb{E}\left[\mathcal{G}_i^2 \mathcal{G}_j\right] - 2 q_i \mathbb{E}\left[\mathcal{G}_i \mathcal{G}_j\right] + q_i^2 \mathbb{E}\left[\mathcal{G}_j\right] - q_j \mathbb{E}\left[\mathcal{G}_i^2\right] + 2 q_i q_j \mathbb{E}\left[\mathcal{G}_i\right] - q_i^2 q_j \\
&= -2 q_i \left(\mathbb{E}\left[\mathcal{G}_i \mathcal{G}_j\right] - q_i q_j\right) \\
&= -2 q_i \left(1 - q_i^2\right)\left(1 - q_j^2\right)\left(2\beta + 4\beta^2 \mathrm{var}\left(\mathscr{G}\right)\right) u_i u_j + \mathcal{O}\left(u_i^3, u_j^3\right) \\
&= \mathcal{O}\left(m^{-1}\right).
\end{aligned}
$$

##### S2.2.4 Fourth moments

**Four distinct indices ($\mathbb{E}[\mathcal{G}_i \mathcal{G}_j \mathcal{G}_k \mathcal{G}_l]$)** Let $i, j, k, l$ be four distinct indices. We again expand $\mathbb{E}\left[\mathcal{G}_i \mathcal{G}_j \mathcal{G}_k \mathcal{G}_l\right]$ in $u_i, u_j, u_k, u_l$.

Zeroth term:

$$
\begin{aligned}
\mathbb{E}\left[\mathcal{G}_i\mathcal{G}_j\mathcal{G}_k\mathcal{G}_l\right]\big|_{u_i,\ldots,u_l=0} &= \tanh\lambda_i\tanh\lambda_j\tanh\lambda_k\tanh\lambda_l, \\
\frac{\partial}{\partial u_i}\mathbb{E}\left[\mathcal{G}_i\mathcal{G}_j\mathcal{G}_k\mathcal{G}_l\right]\Big|_{u_i,\ldots,u_l=0} &= 2\beta\tanh\lambda_j\tanh\lambda_k\tanh\lambda_l\left(1-\tanh^2\lambda_i\right)\mathbb{E}\left[\mathscr{G}\right]_{u_i,\ldots,u_l=0}, \\
\frac{\partial^2}{\partial u_i^2}\mathbb{E}\left[\mathcal{G}_i\mathcal{G}_j\mathcal{G}_k\mathcal{G}_l\right]\Big|_{u_i,\ldots,u_l=0} &= -8\beta^2\tanh\lambda_i\tanh\lambda_j\tanh\lambda_k\tanh\lambda_l\left(1-\tanh^2\lambda_i\right)\mathbb{E}\left[\mathscr{G}\right]^2_{u_i,\ldots,u_l=0}, \\
\frac{\partial^2}{\partial u_i\partial u_j}\mathbb{E}\left[\mathcal{G}_i\mathcal{G}_j\mathcal{G}_k\mathcal{G}_l\right]\Big|_{u_i,\ldots,u_l=0} &= 2\beta\tanh\lambda_k\tanh\lambda_l\left(1-\tanh^2\lambda_i\tanh^2\lambda_j\right) \\
&\quad + 4\beta^2\tanh\lambda_k\tanh\lambda_l\left(1-\tanh^2\lambda_i\tanh^2\lambda_j\right)\mathbb{E}\left[\mathscr{G}^2\right]_{u_i,\ldots,u_l=0} \\
&\quad + 4\beta^2\tanh\lambda_k\tanh\lambda_l\left(2\tanh^2\lambda_i\tanh^2\lambda_j-\tanh^2\lambda_i-\tanh^2\lambda_j\right)\mathbb{E}\left[\mathscr{G}\right]^2_{u_i,\ldots,u_l=0}.
\end{aligned}
$$

We now have the following expansion of the fourth moment:

$$
\begin{aligned}
\mathbb{E}\left[\mathcal{G}_i\mathcal{G}_j\mathcal{G}_k\mathcal{G}_l\right] = {} & \mathcal{O}\left(u_i^3,u_j^3,u_k^3,u_l^3\right) + \tanh\lambda_i\tanh\lambda_j\tanh\lambda_k\tanh\lambda_l \\
& + 2\beta\tanh\lambda_j\tanh\lambda_k\tanh\lambda_l\left(1-\tanh^2\lambda_i\right)\mathbb{E}\left[\mathscr{G}\right]_{u_i,\ldots,u_l=0}u_i \\
& + 2\beta\tanh\lambda_i\tanh\lambda_k\tanh\lambda_l\left(1-\tanh^2\lambda_j\right)\mathbb{E}\left[\mathscr{G}\right]_{u_i,\ldots,u_l=0}u_j \\
& + 2\beta\tanh\lambda_i\tanh\lambda_j\tanh\lambda_l\left(1-\tanh^2\lambda_k\right)\mathbb{E}\left[\mathscr{G}\right]_{u_i,\ldots,u_l=0}u_k \\
& + 2\beta\tanh\lambda_i\tanh\lambda_j\tanh\lambda_k\left(1-\tanh^2\lambda_l\right)\mathbb{E}\left[\mathscr{G}\right]_{u_i,\ldots,u_l=0}u_l \\
& - 4\beta^2\tanh\lambda_i\tanh\lambda_j\tanh\lambda_k\tanh\lambda_l\left(1-\tanh^2\lambda_i\right)\mathbb{E}\left[\mathscr{G}\right]^2_{u_i,\ldots,u_l=0}u_i^2 \\
& - 4\beta^2\tanh\lambda_i\tanh\lambda_j\tanh\lambda_k\tanh\lambda_l\left(1-\tanh^2\lambda_j\right)\mathbb{E}\left[\mathscr{G}\right]^2_{u_i,\ldots,u_l=0}u_j^2 \\
& - 4\beta^2\tanh\lambda_i\tanh\lambda_j\tanh\lambda_k\tanh\lambda_l\left(1-\tanh^2\lambda_k\right)\mathbb{E}\left[\mathscr{G}\right]^2_{u_i,\ldots,u_l=0}u_k^2 \\
& - 4\beta^2\tanh\lambda_i\tanh\lambda_j\tanh\lambda_k\tanh\lambda_l\left(1-\tanh^2\lambda_l\right)\mathbb{E}\left[\mathscr{G}\right]^2_{u_i,\ldots,u_l=0}u_l^2 \\
& + 2\beta\tanh\lambda_k\tanh\lambda_l\left(1-\tanh^2\lambda_i\tanh^2\lambda_j\right)u_iu_j \\
& + 4\beta^2\tanh\lambda_k\tanh\lambda_l\left(1-\tanh^2\lambda_i\tanh^2\lambda_j\right)\mathbb{E}\left[\mathscr{G}^2\right]_{u_i,\ldots,u_l=0}u_iu_j \\
& + 4\beta^2\tanh\lambda_k\tanh\lambda_l\left(2\tanh^2\lambda_i\tanh^2\lambda_j-\tanh^2\lambda_i-\tanh^2\lambda_j\right)\mathbb{E}\left[\mathscr{G}\right]^2_{u_i,\ldots,u_l=0}u_iu_j \\
& + 2\beta\tanh\lambda_j\tanh\lambda_l\left(1-\tanh^2\lambda_i\tanh^2\lambda_k\right)u_iu_k \\
& + 4\beta^2\tanh\lambda_j\tanh\lambda_l\left(1-\tanh^2\lambda_i\tanh^2\lambda_k\right)\mathbb{E}\left[\mathscr{G}^2\right]_{u_i,\ldots,u_l=0}u_iu_k \\
& + 4\beta^2\tanh\lambda_j\tanh\lambda_l\left(2\tanh^2\lambda_i\tanh^2\lambda_k-\tanh^2\lambda_i-\tanh^2\lambda_k\right)\mathbb{E}\left[\mathscr{G}\right]^2_{u_i,\ldots,u_l=0}u_iu_k \\
& + 2\beta\tanh\lambda_j\tanh\lambda_k\left(1-\tanh^2\lambda_i\tanh^2\lambda_l\right)u_iu_l \\
& + 4\beta^2\tanh\lambda_j\tanh\lambda_k\left(1-\tanh^2\lambda_i\tanh^2\lambda_l\right)\mathbb{E}\left[\mathscr{G}^2\right]_{u_i,\ldots,u_l=0}u_iu_l \\
& + 4\beta^2\tanh\lambda_j\tanh\lambda_k\left(2\tanh^2\lambda_i\tanh^2\lambda_l-\tanh^2\lambda_i-\tanh^2\lambda_l\right)\mathbb{E}\left[\mathscr{G}\right]^2_{u_i,\ldots,u_l=0}u_iu_l \\
& + 2\beta\tanh\lambda_i\tanh\lambda_l\left(1-\tanh^2\lambda_j\tanh^2\lambda_k\right)u_ju_k
\end{aligned}
$$

$$+ 4\beta^2 \tanh\lambda_i \tanh\lambda_l \left(1 - \tanh^2\lambda_j \tanh^2\lambda_k\right) \mathbb{E}\left[\mathscr{G}^2\right]_{u_i,\ldots,u_l=0} u_j u_k$$

$$+ 4\beta^2 \tanh\lambda_i \tanh\lambda_l \left(2\tanh^2\lambda_j \tanh^2\lambda_k - \tanh^2\lambda_j - \tanh^2\lambda_k\right) \mathbb{E}\left[\mathscr{G}\right]^2_{u_i,\ldots,u_l=0} u_j u_k$$

$$+ 2\beta \tanh\lambda_i \tanh\lambda_k \left(1 - \tanh^2\lambda_j \tanh^2\lambda_l\right) u_j u_l$$

$$+ 4\beta^2 \tanh\lambda_i \tanh\lambda_k \left(1 - \tanh^2\lambda_j \tanh^2\lambda_l\right) \mathbb{E}\left[\mathscr{G}^2\right]_{u_i,\ldots,u_l=0} u_j u_l$$

$$+ 4\beta^2 \tanh\lambda_i \tanh\lambda_k \left(2\tanh^2\lambda_j \tanh^2\lambda_l - \tanh^2\lambda_j - \tanh^2\lambda_l\right) \mathbb{E}\left[\mathscr{G}\right]^2_{u_i,\ldots,u_l=0} u_j u_l$$

$$+ 2\beta \tanh\lambda_i \tanh\lambda_j \left(1 - \tanh^2\lambda_k \tanh^2\lambda_l\right) u_k u_l$$

$$+ 4\beta^2 \tanh\lambda_i \tanh\lambda_j \left(1 - \tanh^2\lambda_k \tanh^2\lambda_l\right) \mathbb{E}\left[\mathscr{G}^2\right]_{u_i,\ldots,u_l=0} u_k u_l$$

$$+ 4\beta^2 \tanh\lambda_i \tanh\lambda_j \left(2\tanh^2\lambda_k \tanh^2\lambda_l - \tanh^2\lambda_k - \tanh^2\lambda_l\right) \mathbb{E}\left[\mathscr{G}\right]^2_{u_i,\ldots,u_l=0} u_k u_l. \tag{31}$$

Generalizing the derivation of (24), we have

$$\begin{aligned}\mathbb{E}\left[\mathscr{G}\right]_{u_i,\ldots,u_l=0} =& \mathcal{O}\left(u_i^2, u_j^2, u_k^2, u_l^2\right) + \mathbb{E}\left[\mathscr{G}\right] \\ &- \left(1 + 2\beta\left(\mathbb{E}\left[\mathscr{G}^2\right] - \mathbb{E}\left[\mathscr{G}\right]^2\right)\right)\left(\tanh\lambda_i u_i + \tanh\lambda_j u_j + \tanh\lambda_k u_k + \tanh\lambda_l u_l\right).\end{aligned}$$

Substituting this into (31) yields

$$\begin{aligned}\mathbb{E}\left[\mathcal{G}_i\mathcal{G}_j\mathcal{G}_k\mathcal{G}_l\right] = & \mathcal{O}\left(u_i^3, u_j^3, u_k^3, u_l^3\right) + \tanh\lambda_i \tanh\lambda_j \tanh\lambda_k \tanh\lambda_l \\
&+ 2\beta \tanh\lambda_j \tanh\lambda_k \tanh\lambda_l \left(1 - \tanh^2\lambda_i\right) \mathbb{E}\left[\mathscr{G}\right] u_i \\
&+ 2\beta \tanh\lambda_i \tanh\lambda_k \tanh\lambda_l \left(1 - \tanh^2\lambda_j\right) \mathbb{E}\left[\mathscr{G}\right] u_j \\
&+ 2\beta \tanh\lambda_i \tanh\lambda_j \tanh\lambda_l \left(1 - \tanh^2\lambda_k\right) \mathbb{E}\left[\mathscr{G}\right] u_k \\
&+ 2\beta \tanh\lambda_i \tanh\lambda_j \tanh\lambda_k \left(1 - \tanh^2\lambda_l\right) \mathbb{E}\left[\mathscr{G}\right] u_l \\
&- 2\beta \tanh\lambda_i \tanh\lambda_j \tanh\lambda_k \tanh\lambda_l \left(1 - \tanh^2\lambda_i\right)\left(1 + 2\beta\mathbb{E}\left[\mathscr{G}^2\right]\right) u_i^2 \\
&- 2\beta \tanh\lambda_i \tanh\lambda_j \tanh\lambda_k \tanh\lambda_l \left(1 - \tanh^2\lambda_j\right)\left(1 + 2\beta\mathbb{E}\left[\mathscr{G}^2\right]\right) u_j^2 \\
&- 2\beta \tanh\lambda_i \tanh\lambda_j \tanh\lambda_k \tanh\lambda_l \left(1 - \tanh^2\lambda_k\right)\left(1 + 2\beta\mathbb{E}\left[\mathscr{G}^2\right]\right) u_k^2 \\
&- 2\beta \tanh\lambda_i \tanh\lambda_j \tanh\lambda_k \tanh\lambda_l \left(1 - \tanh^2\lambda_l\right)\left(1 + 2\beta\mathbb{E}\left[\mathscr{G}^2\right]\right) u_l^2 \\
&+ 2\beta \tanh\lambda_k \tanh\lambda_l \left(1 - \tanh^2\lambda_i\right)\left(1 - \tanh^2\lambda_j\right)\left(1 + 2\beta\mathbb{E}\left[\mathscr{G}^2\right]\right) u_i u_j \\
&+ 2\beta \tanh\lambda_j \tanh\lambda_l \left(1 - \tanh^2\lambda_i\right)\left(1 - \tanh^2\lambda_k\right)\left(1 + 2\beta\mathbb{E}\left[\mathscr{G}^2\right]\right) u_i u_k \\
&+ 2\beta \tanh\lambda_j \tanh\lambda_k \left(1 - \tanh^2\lambda_i\right)\left(1 - \tanh^2\lambda_l\right)\left(1 + 2\beta\mathbb{E}\left[\mathscr{G}^2\right]\right) u_i u_l \\
&+ 2\beta \tanh\lambda_i \tanh\lambda_l \left(1 - \tanh^2\lambda_j\right)\left(1 - \tanh^2\lambda_k\right)\left(1 + 2\beta\mathbb{E}\left[\mathscr{G}^2\right]\right) u_j u_k \\
&+ 2\beta \tanh\lambda_i \tanh\lambda_k \left(1 - \tanh^2\lambda_j\right)\left(1 - \tanh^2\lambda_l\right)\left(1 + 2\beta\mathbb{E}\left[\mathscr{G}^2\right]\right) u_j u_l \\
&+ 2\beta \tanh\lambda_i \tanh\lambda_j \left(1 - \tanh^2\lambda_k\right)\left(1 - \tanh^2\lambda_l\right)\left(1 + 2\beta\mathbb{E}\left[\mathscr{G}^2\right]\right) u_k u_l.\end{aligned}$$

Substituting (25) yields

$$
\begin{aligned}
\mathbb{E}\left[\mathcal{G}_i\mathcal{G}_j\mathcal{G}_k\mathcal{G}_l\right] = \mathcal{O}\left(u_i^3, u_j^3, u_k^3, u_l^3\right) &+ q_iq_jq_kq_l \\
&+ 2\beta q_kq_l\left(1-q_i^2\right)\left(1-q_j^2\right)\left(1+2\beta\mathrm{var}\left(\mathscr{G}^2\right)\right)u_iu_j \\
&+ 2\beta q_jq_l\left(1-q_i^2\right)\left(1-q_k^2\right)\left(1+2\beta\mathrm{var}\left(\mathscr{G}^2\right)\right)u_iu_k \\
&+ 2\beta q_jq_k\left(1-q_i^2\right)\left(1-q_l^2\right)\left(1+2\beta\mathrm{var}\left(\mathscr{G}^2\right)\right)u_iu_l \\
&+ 2\beta q_iq_l\left(1-q_j^2\right)\left(1-q_k^2\right)\left(1+2\beta\mathrm{var}\left(\mathscr{G}^2\right)\right)u_ju_k \\
&+ 2\beta q_iq_k\left(1-q_j^2\right)\left(1-q_l^2\right)\left(1+2\beta\mathrm{var}\left(\mathscr{G}^2\right)\right)u_ju_l \\
&+ 2\beta q_iq_j\left(1-q_k^2\right)\left(1-q_l^2\right)\left(1+2\beta\mathrm{var}\left(\mathscr{G}^2\right)\right)u_ku_l.
\end{aligned}
$$

As with the third moment, the nontrivial terms here match the expression for the second moment from (27). Together with (30), this implies the fourth central moment is zero to second degree in $u$:

$$
\begin{aligned}
\mathbb{E}\left[\left(\mathcal{G}_i-q_i\right)\left(\mathcal{G}_j-q_j\right)\left(\mathcal{G}_k-q_k\right)\left(\mathcal{G}_l-q_l\right)\right] =& \mathbb{E}\left[\mathcal{G}_i\mathcal{G}_j\mathcal{G}_k\mathcal{G}_l\right] - q_i\mathbb{E}\left[\left(\mathcal{G}_j-q_j\right)\left(\mathcal{G}_k-q_k\right)\left(\mathcal{G}_l-q_l\right)\right] - q_j\mathbb{E}\left[\left(\mathcal{G}_i-q_i\right)\left(\mathcal{G}_k-q_k\right)\left(\mathcal{G}_l-q_l\right)\right] \\
&- q_k\mathbb{E}\left[\left(\mathcal{G}_i-q_i\right)\left(\mathcal{G}_j-q_j\right)\left(\mathcal{G}_l-q_l\right)\right] - q_l\mathbb{E}\left[\left(\mathcal{G}_i-q_i\right)\left(\mathcal{G}_j-q_j\right)\left(\mathcal{G}_k-q_k\right)\right] \\
&- q_iq_j\mathbb{E}\left[\mathcal{G}_k\mathcal{G}_l\right] - q_iq_k\mathbb{E}\left[\mathcal{G}_j\mathcal{G}_l\right] - q_iq_l\mathbb{E}\left[\mathcal{G}_k\mathcal{G}_j\right] - q_jq_k\mathbb{E}\left[\mathcal{G}_i\mathcal{G}_l\right] - q_jq_l\mathbb{E}\left[\mathcal{G}_i\mathcal{G}_k\right] \\
&- q_kq_l\mathbb{E}\left[\mathcal{G}_i\mathcal{G}_j\right] + 5q_iq_jq_kq_l \\
=& \mathcal{O}\left(u_i^3, u_j^3, u_k^3, u_l^3\right) = \mathcal{O}\left(m^{-3/2}\right).
\end{aligned}
$$

**Mixed fourth moments** Here we consider central fourth moments with repeated indices. These reduce to lower-order moments using $\mathcal{G}_i^2 = 1$.

Let $i, j, k$ be any three distinct indices. Simplifying and then substituting (27) and (29) yields

$$
\begin{aligned}
\mathbb{E}\left[\left(\mathcal{G}_i-q_i\right)^2\left(\mathcal{G}_j-q_j\right)\left(\mathcal{G}_k-q_k\right)\right] =& \mathbb{E}\left[\mathcal{G}_i^2\mathcal{G}_j\mathcal{G}_k\right] - 2q_i\mathbb{E}\left[\mathcal{G}_i\mathcal{G}_j\mathcal{G}_k\right] + q_i^2\mathbb{E}\left[\mathcal{G}_j\mathcal{G}_k\right] - q_j\mathbb{E}\left[\mathcal{G}_i^2\mathcal{G}_k\right] \\
&+ 2q_iq_j\mathbb{E}\left[\mathcal{G}_i\mathcal{G}_k\right] - q_k\mathbb{E}\left[\mathcal{G}_i^2\mathcal{G}_j\right] + 2q_iq_k\mathbb{E}\left[\mathcal{G}_i\mathcal{G}_j\right] + q_jq_k\mathbb{E}\left[\mathcal{G}_i^2\right] - 3q_i^2q_jq_k \\
=& \left(1-q_i^2\right)\left(1-q_j^2\right)\left(1-q_k^2\right)\left(2\beta+4\beta^2\mathrm{var}\left(\mathscr{G}\right)\right)u_ju_k + \mathcal{O}\left(u_i^3, u_j^3, u_k^3\right) \\
=& \mathcal{O}\left(m^{-1}\right).
\end{aligned}
$$

Now consider $\left(\mathcal{G}_i-q_i\right)^3\left(\mathcal{G}_j-q_j\right)$:

$$
\begin{aligned}
\mathbb{E}\left[\left(\mathcal{G}_i-q_i\right)^3\left(\mathcal{G}_j-q_j\right)\right] &= \mathbb{E}\left[\mathcal{G}_i^3\mathcal{G}_j\right] - 3q_i\mathbb{E}\left[\mathcal{G}_i^2\mathcal{G}_j\right] + 3q_i^2\mathbb{E}\left[\mathcal{G}_i\mathcal{G}_j\right] - q_i^3\mathbb{E}\left[\mathcal{G}_j\right] \\
&\quad - q_j\mathbb{E}\left[\mathcal{G}_i^3\right] + 3q_iq_j\mathbb{E}\left[\mathcal{G}_i^2\right] - 3q_i^2q_j\mathbb{E}\left[\mathcal{G}_i\right] + q_i^3q_j \\
&= \left(1+3q_i^2\right)\left(\mathbb{E}\left[\mathcal{G}_i\mathcal{G}_j\right] - q_iq_j\right)
\end{aligned}
$$

$$= \left(1-q_i^2\right)\left(1-q_j^2\right)\left(1+3q_i^2\right)\left(2\beta+4\beta^2\mathrm{var}\left(\mathscr{G}\right)\right)u_iu_j+\mathcal{O}\left(u_i^3,u_j^3\right)$$
$$= \mathcal{O}\left(m^{-1}\right).$$

Last, consider $\left(\mathcal{G}_i-q_i\right)^2\left(\mathcal{G}_j-q_j\right)^2$:

$$\begin{aligned}
\mathbb{E}\left[\left(\mathcal{G}_i-q_i\right)^2\left(\mathcal{G}_j-q_j\right)^2\right] &= \mathbb{E}\left[\mathcal{G}_i^2\mathcal{G}_j^2\right]-2q_j\mathbb{E}\left[\mathcal{G}_i^2\mathcal{G}_j\right]+q_j^2\mathbb{E}\left[\mathcal{G}_i^2\right]-2q_i\mathbb{E}\left[\mathcal{G}_i\mathcal{G}_j^2\right]\\
&\quad+4q_iq_j\mathbb{E}\left[\mathcal{G}_i\mathcal{G}_j\right]-2q_iq_j^2\mathbb{E}\left[\mathcal{G}_i\right]+q_i^2\mathbb{E}\left[\mathcal{G}_j^2\right]-2q_i^2q_j\mathbb{E}\left[\mathcal{G}_j\right]+q_i^2q_j^2\\
&= \left(1-q_i^2\right)\left(1-q_j^2\right)+4q_iq_j\left(\mathbb{E}\left[\mathcal{G}_i\mathcal{G}_j\right]-q_iq_j\right)\\
&= \left(1-q_i^2\right)\left(1-q_j^2\right)+4q_iq_j\left(1-q_i^2\right)\left(1-q_j^2\right)\left(2\beta+4\beta^2\mathrm{var}\left(\mathscr{G}\right)\right)u_iu_j+\mathcal{O}\left(u_i^3,u_j^3\right).
\end{aligned}$$

Therefore the corresponding standardized moment is bound as

$$\begin{aligned}
\mathbb{E}\left[\frac{\left(\mathcal{G}_i-q_i\right)^2}{1-q_i^2}\frac{\left(\mathcal{G}_j-q_j\right)^2}{1-q_j^2}\right] &= 1+4q_iq_j\left(2\beta+4\beta^2\mathrm{var}\left(\mathscr{G}\right)\right)u_iu_j+\mathcal{O}\left(u_i^3,u_j^3\right)\\
&= 1+\mathcal{O}\left(m^{-1}\right).
\end{aligned}$$

#### S2.3 The limiting spectral distribution of covariance matrices for a class of random matrices with dependent entries

Here we establish the limiting spectral distribution for a class of random matrices with dependent entries to which our matrix of standardized genotypes $Z$ belongs. For simplicity, we will assume that the equilibrium covariances among all causal variants is constant, though our results are readily extended to the general case where $\mathbb{E}[\{n^{-1}Z^TZ\}_{ij}]=\mathcal{O}(1/m)$ for all $i, j\neq i$.

##### S2.3.1 Introduction and primary result

For an $n\times n$ Hermitian matrix $A$, we let $\lambda_1(A),\ldots,\lambda_n(A)\in\mathbb{R}$ denote the eigenvalues of $A$. The empirical spectral distribution (ESD) $F^A$ of $A$ is defined as

$$F^A(x) := \frac{1}{n}\left|\{1\leq i\leq n:\lambda_i(A)\leq x\}\right|,$$

where $|E|$ denotes the cardinality of the finite set $E$. The Marčenko–Pastur distribution $F_\tau^{\mathrm{MP}}$ with parameter $\tau>0$ is the probability distribution function with density

$$f_\tau^{\mathrm{MP}}(x) := \begin{cases} \frac{1}{2\pi\tau x}\sqrt{(b-x)(x-a)}, & \text{if } a\leq x\leq b,\\ 0, & \text{otherwise,}\end{cases}$$

and a point mass $1 - 1/\tau$ at the origin if $\tau > 1$, where $a := (1-\sqrt{\tau})^2$ and $b := (1+\sqrt{\tau})^2$.

Let $z \in \mathbb{R}^p$ be a random vector. We let $Z$ be an $n \times p$ matrix whose rows are independent and identically distributed (iid) copies of $z$. We will be interested in the eigenvalues of $\frac{1}{p}ZZ^T$. We view $n$ as a large asymptotic parameter tending to infinity; $p$ and $m$ will be two additional parameters which we assume tend to infinity with $n$. Asymptotic notation (such as $\mathcal{O}$, $o$) will be used under the assumption that $n, m, p \to \infty$.

For the model under consideration, we make the following assumptions regarding the random vector $z$. The random vector $z$ has mean zero and covariance matrix

$$\Sigma := \begin{bmatrix} \Sigma' & 0 \\ 0 & I_{p-m} \end{bmatrix},$$

where $\Sigma'$ is an $m \times m$ matrix with entries

$$\Sigma'_{ij} := \begin{cases} 2\mu(m), & \text{if } i \neq j, \\ 1 + \mu(m), & \text{if } i = j. \end{cases} \tag{32}$$

Here $I_n$ denotes the $n \times n$ identity matrix. Additionally, we assume $\mu(m) = \mathcal{O}(1/m)$. We also assume the following about the entries of $z = (z_i)_{i=1}^p$:

1. There exists a constant $\kappa > 0$ so that $\sup_{1 \le i \le p} |z_i| \le \kappa$ with probability 1.
2. $\mathbb{E}[z_i^2 z_j^2] = 1 + o(1)$ uniformly for all distinct $i, j$.
3. $\mathbb{E}[z_i^3 z_j] = o(1)$ uniformly for all distinct $i, j$.
4. One has
$$\mathbb{E}[z_i z_j z_k z_l] = \begin{cases} o\left(\frac{1}{m}\right), & \text{if } |\{i,j,k,l\}| = 4, \\ o\left(\frac{1}{\sqrt{m}}\right), & \text{if } |\{i,j,k,l\}| = 3, \end{cases} \tag{33}$$
uniformly in $i, j, k, l$.

**Theorem 1.** *Suppose $\frac{n}{p} \to \tau \in (0, \infty)$ as $n \to \infty$ and $m \ge cp$ for some constant $c > 0$. Then, under the assumptions above, the ESD $F^{\frac{1}{p}ZZ^T}$ of $\frac{1}{p}ZZ^T$ converges almost surely to the Marčenko–Pastur distribution $F_\tau^{\mathrm{MP}}$ as $n \to \infty$.*

The remainder of this section is devoted to the proof of Theorem 1. Throughout, we assume $m$, $n$, $p$, $z$, and $Z$ satisfy the assumptions of Theorem 1.

##### S2.3.2 Notation and overview

$I_n$ denotes the $n \times n$ identity matrix. Often we will simply write $I$ when the size can be deduced from context. For a matrix $A$, we let $\|A\|$ denote the spectral (operator) norm of $A$ and $\|A\|_2$ denote the Hilbert–Schmidt

(Frobenius) norm of $A$ defined by

$$\|A\|_2 := \sqrt{\operatorname{tr}(AA^*)} = \sqrt{\operatorname{tr}(A^*A)},$$

where $A^*$ denotes the conjugate transpose of $A$. We will exploit the fact that the spectral norm of $A$ is an upper bound for the spectral norm of any sub-matrix of $A$.

For $\alpha \in \mathbb{C}^+ := \{w \in \mathbb{C} : \Im(w) > 0\}$, we define the resolvent of $\frac{1}{p}ZZ^T$ as

$$R(\alpha) := \left(\frac{1}{p}ZZ^T - \alpha I\right)^{-1}.$$

$Z^{(k)}$ is the $(n-1) \times p$ matrix constructed from $Z$ by removing the $k$-th row. Let $R^{(k)}$ be the resolvent of $\frac{1}{p}Z^{(k)}Z^{(k)T}$, i.e.,

$$R^{(k)}(\alpha) := \left(\frac{1}{p}Z^{(k)}Z^{(k)T} - \alpha I\right)^{-1}, \qquad \alpha \in \mathbb{C}^+.$$

Define $s_n$ to be the Stieltjes transform of the ESD $F^{\frac{1}{p}ZZ^T}$. In other words,

$$s_n(\alpha) := \frac{1}{n}\operatorname{tr} R(\alpha), \qquad \alpha \in \mathbb{C}^+. \tag{34}$$

We let $s$ be the Stieltjes transform of $F_\tau^{\mathrm{MP}}$. From (Bai *et al.*, 2010, Chapter 3) it follows that $s$ satisfies the equation

$$s(\alpha) = \frac{1}{1 - \alpha - \tau - \tau\alpha s(\alpha)}, \qquad \alpha \in \mathbb{C}^+. \tag{35}$$

We prove Theorem 1 by showing that $s_n$ converges to $s$ as $n$ tends to infinity. This is a standard proof method in random matrix theory (see e.g. (Bai *et al.*, 2010) for details). However, the dependence between the entries introduces new difficulties in establishing this convergence. The main technical advance in this work involves dealing with the dependence amongst the entries.

In Section S2.3.3 we present the main tools from linear algebra and probability theory that are required for the proof. A concentration inequality is presented in Section S2.3.4 which allows us to pass between the Stieltjes transform and its expected value. Section S2.3.5 contains a version of the Hanson–Wright inequality for the rows of $Z$; this section is the main technical tool for dealing with the dependence amongst the entries. We finally complete the proof of Theorem 1 in Section S2.3.6

###### S2.3.3 Tools from probability theory and linear algebra

We begin by introducing some tools from probability theory and linear algebra.

**Lemma 2.** *For any $\alpha \in \mathbb{C}^+$ and any $1 \le k \le n$,*

$$\left|\operatorname{tr} R(\alpha) - \operatorname{tr} R^{(k)}(\alpha)\right| \le \frac{1}{\Im(\alpha)}.$$

*Proof.* This follows from Eq. (A.1.12) in (Bai *et al.*, 2010). □

**Lemma 3.** *If $\mu$ is a probability measure supported on $[0, \infty)$ and $s_\mu$ is its Stieltjes transform defined by*

$$s_\mu(\alpha) := \int_0^\infty \frac{1}{x - \alpha} d\mu(x), \qquad \alpha \in \mathbb{C}^+, \tag{36}$$

*then $|s_\mu(\alpha)| \le \frac{1}{\Im(\alpha)}$ and*

$$\Im s_\mu(\alpha) > 0, \qquad \Im(\alpha s_\mu(\alpha)) \ge 0$$

*for all $\alpha \in \mathbb{C}^+$.*

*Proof.* The first bound follows from the triangle inequality:

$$|s_\mu(\alpha)| \le \int_0^\infty \frac{1}{|x - \alpha|} d\mu(x) \le \int_0^\infty \frac{1}{\Im \alpha} d\mu(x) = \frac{1}{\Im \alpha}.$$

For the other bounds, since

$$s_\mu(\alpha) = \int_0^\infty \frac{x - \bar{\alpha}}{|x - \alpha|^2} d\mu(x),$$

it follows that $\Im s_\mu(\alpha) > 0$. Similarly,

$$\Im(\alpha s_\mu(\alpha)) = \Im(\alpha) \int_0^\infty \frac{x}{|x - \alpha|^2} d\mu(x) \ge 0,$$

which completes the proof. □

**Lemma 4** (Stability)**.** *Suppose $\mu$ is a probability measure supported on $[0, \infty)$ and $s_\mu$ is its Stieltjes transform defined in (36). If*

$$s_\mu(\alpha) = \frac{1}{1 - \alpha - \tau - \tau \alpha s_\mu(\alpha)} + \varepsilon \tag{37}$$

*for some $\alpha \in \mathbb{C}^+$, some $\varepsilon \in \mathbb{C}$, and some $\tau > 0$, then*

$$|s_\mu(\alpha) - s(\alpha)| \le \frac{|\varepsilon|}{\Im \alpha} |1 - \alpha - \tau - \tau \alpha s_\mu(\alpha)|,$$

*where $s$ is the Stieltjes transform of the Marčenko–Pastur distribution function $F_y^{\mathrm{MP}}$ (see (35)).*

*Proof.* Using (35) and (37), we have

$$-\tau \alpha s_\mu^2(\alpha) + s_\mu(\alpha)(1 - \alpha - \tau) - 1 = \varepsilon(1 - \alpha - \tau - \tau \alpha s_\mu(\alpha))$$

and

$$-\tau\alpha s^2(\alpha) + s(\alpha)(1-\alpha-\tau) - 1 = 0.$$

Subtracting the two equations gives

$$|s_\mu(\alpha) - s(\alpha)||-\tau\alpha(s_\mu(\alpha)+s(\alpha)) + 1 - \alpha - \tau| = |\varepsilon||1-\alpha-\tau-\tau\alpha s_\mu(\alpha)|.$$

In view of Lemma 3,

$$|-\tau\alpha(s_\mu(\alpha)+s(\alpha)) + 1 - \alpha - \tau| \geq \tau\Im(\alpha(s_\mu(\alpha)+s(\alpha))) + \Im(\alpha) \geq \Im(\alpha),$$

and the conclusion follows. □

Throughout the proof, we will utilize the resolvent identity:

$$A^{-1} - B^{-1} = A^{-1}(B-A)B^{-1}, \tag{38}$$

which holds for invertible matrices $A$ and $B$. We will also use the bound

$$\|(A-\alpha I)^{-1}\| \leq \frac{1}{\Im(\alpha)} \tag{39}$$

which follows from the spectral theorem and holds for any Hermitian matrix $A$ and any $\alpha \in \mathbb{C}^+$. We will also need the following bound.

**Lemma 5.** *Let $A$ be an $n \times p$ matrix. Then*

$$A^*(AA^* - \alpha I)^{-1}A = (A^*A - \alpha I)^{-1}A^*A \tag{40}$$

*for all $\alpha \in \mathbb{C}^+$. In addition,*

$$\|A^*(AA^* - \alpha I)^{-1}A\| \leq 1 + \frac{|\alpha|}{\Im\alpha} \tag{41}$$

*for all $\alpha \in \mathbb{C}^+$.*

*Proof.* For $|\alpha| > \|AA^*\|$ the Neumann series for the resolvent $(AA^* - \alpha I)^{-1}$ gives

$$(AA^* - \alpha I)^{-1} = -\frac{1}{\alpha}I - \sum_{k=1}^{\infty}\frac{(AA^*)^k}{\alpha^{k+1}}.$$

It follows that

$$A^*(AA^* - \alpha I)^{-1}A = -\frac{A^*A}{\alpha} - \sum_{k=1}^{\infty} \frac{(A^*A)^k A^*A}{\alpha^{k+1}} = (A^*A - \alpha I)^{-1}A^*A.$$

This establishes (40) for $\alpha \in \mathbb{C}^+$ with $|\alpha| > \|AA^*\|$. The identity can be extended to all of $\mathbb{C}^+$ by analytic continuation since the entries of the resolvent are analytic in $\mathbb{C}^+$. Alternatively, (40) can be derived using the singular value decomposition for $A$.

The bound in (41) can be established using (40). Indeed, by (40)

$$\begin{aligned} A^*(AA^* - \alpha I)^{-1}A &= (A^*A - \alpha I)^{-1}A^*A \\ &= (A^*A - \alpha I)^{-1}(A^*A - \alpha I + \alpha I) \\ &= I + \alpha(A^*A - \alpha I)^{-1}. \end{aligned}$$

We conclude that

$$\|A^*(AA^* - \alpha I)^{-1}A\| \leq 1 + \frac{\alpha}{\Im(\alpha)}$$

by the triangle inequality and (39). $\square$

We will also take advantage of the bound

$$|\mathrm{tr}\,(A)| \leq \mathrm{rank}\,(A)\|A\|, \tag{42}$$

which holds for any square matrix $A$ and follows from (Bai *et al.*, 2010, Corollary A.12).

###### S2.3.4 Concentration

Recall the Stieltjes transform $s_n$ defined in (34). The following results show that the Stieltjes transform $s_n$ concentrates around its expected value $\mathbb{E}[s_n]$.

**Lemma 6.** *For any $\alpha \in \mathbb{C}^+$ and any $t > 0$,*

$$\mathbb{P}(|s_n(\alpha) - \mathbb{E}[s_n(\alpha)]| > t) \leq C \exp\left(-cnt^2(\Im\alpha)^2\right),$$

*where $C, c > 0$ are absolute constants.*

*Proof.* Let $\tilde{Z}^{(k)}$ be formed from the matrix $Z$ by replacing the $k$-th row with zeros. Let $\tilde{R}^{(k)}$ be the resolvent of $\tilde{Z}^{(k)}\tilde{Z}^{(k)T}$ defined by

$$\tilde{R}^{(k)}(\alpha) := \left(\tilde{Z}^{(k)}\tilde{Z}^{(k)T} - \alpha I\right)^{-1}, \qquad \alpha \in \mathbb{C}^+.$$

Then for any $1 \le k \le n$,

$$\left| \operatorname{tr} R(\alpha) - \operatorname{tr} \tilde{R}^{(k)}(\alpha) \right| \le \left| \operatorname{tr} R(\alpha) - \operatorname{tr} R^{(k)}(\alpha) \right| + \left| \operatorname{tr} R^{(k)}(\alpha) - \operatorname{tr} \tilde{R}^{(k)}(\alpha) \right|.$$

The first term on the right-hand side can be bounded using Lemma 2. The second term can be controlled using the fact that $Z^{(k)}Z^{(k)T}$ and $\tilde{Z}^{(k)}\tilde{Z}^{(k)T}$ have the same eigenvalues except $\tilde{Z}^{(k)}\tilde{Z}^{(k)T}$ has one additional zero eigenvalue. In other words,

$$\left| \operatorname{tr} R^{(k)}(\alpha) - \operatorname{tr} \tilde{R}^{(k)}(\alpha) \right| = \left| \frac{1}{\alpha} \right| \le \frac{1}{\Im(\alpha)}.$$

We conclude that

$$\left| \operatorname{tr} R(\alpha) - \operatorname{tr} \tilde{R}^{(k)}(\alpha) \right| \le \frac{2}{\Im(\alpha)}$$

for any $1 \le k \le n$. Now since $\operatorname{tr} R(\alpha)$ is a function of the independent random vectors $z_1, \ldots, z_n$, McDiarmid's inequality (see (McDiarmid, 1989) or Theorem 2.1.10 in (Tao, 2011)) implies that

$$\mathbb{P}(|\operatorname{tr} R(\alpha) - \mathbb{E}\operatorname{tr} R(\alpha)| > t) \le C \exp\left( -c(\Im\alpha)^2 \frac{t^2}{n} \right)$$

for any $t > 0$, where $C, c > 0$ are absolute constants. The claim now follows by rescaling. □

**Lemma 7.** *Define the domain*

$$S := \left\{ \alpha \in \mathbb{C}^+ : 1 \ge \Im(\alpha) \ge \frac{\log n}{n^{1/4}}, -n \le \Re(\alpha) \le n \right\}.$$

*One has*

$$\mathbb{P}\left( \sup_{\alpha \in S} |s_n(\alpha) - \mathbb{E}[s_n(\alpha)]| > 2n^{-1/4} \right) \le C \exp\left( -c \log^2 n \right),$$

*where $C, c > 0$ are absolute constants.*

We will utilize an $\varepsilon$-net in order to prove Lemma 7.s

**Definition** (Nets)**.** Let $X$ be a subset of $\mathbb{C}$ and $\varepsilon > 0$. A subset $\mathcal{N}$ of $X$ is called an $\varepsilon$-net of $X$ if every point $x \in X$ can be approximated within $\varepsilon$ by some point $y \in \mathcal{N}$, i.e. so that $|x - y| \le \varepsilon$.

*Proof of Lemma 7.* By Lemma 6,

$$\sup_{\alpha \in S} \mathbb{P}\left( |s_n(\alpha) - \mathbb{E}[s_n(\alpha)]| > n^{-1/4} \right) \le C' \exp\left( -c' \log^2 n \right), \tag{43}$$

where $C', c' > 0$ are absolute constants. The goal is to extend this bound to simultaneously hold for all $\alpha \in S$. We will use a net to do so. Let $\mathcal{N}$ be a $n^{-1}$-net of $S$. A simple volume argument (see for instance

(O'Rourke *et al.*, 2019, Lemma 3.3)) shows that $\mathcal{N}$ can be chosen so that

$$|\mathcal{N}| \leq C_0 n^{C_0}$$

for an absolute constant $C_0 > 0$. By the union bound and (43), we find

$$\mathbb{P}\left(\sup_{\alpha\in\mathcal{N}} |s_n(\alpha) - \mathbb{E}[s_n(\alpha)]| > n^{-1/4}\right) \leq C\exp(-c\log^2 n), \tag{44}$$

where $C, c > 0$ are absolute constants.

We now wish to extend this bound from $\mathcal{N}$ to $S$. To this end, suppose there exists $\alpha \in S$ so that $|s_n(\alpha) - \mathbb{E}s_n(\alpha)| > 2n^{-1/4}$. Then there exists $\alpha' \in \mathcal{N}$ such that $|\alpha - \alpha'| \leq \frac{1}{n}$. Thus, by the resolvent identity (38), (42), and (39), we have

$$\left|\frac{1}{n}\operatorname{tr} R(\alpha) - \frac{1}{n}\operatorname{tr} R(\alpha')\right| \leq |\alpha - \alpha'|\|R(\alpha)\|\|R(\alpha')\| \leq \frac{|\alpha-\alpha'|}{\Im(\alpha)\Im(\alpha')}.$$

The same bound holds for the expected value (by the same argument) and so we deduce that

$$|(s_n(\alpha) - \mathbb{E}[s_n(\alpha))] - (s_n(\alpha') - \mathbb{E}[s_n(\alpha'))]| \leq 2\frac{\sqrt{n}}{\log^2 n}|\alpha - \alpha'| < n^{-1/4}$$

for $n$ sufficiently large since $|\alpha - \alpha'| \leq n^{-1}$. Here we bounded the imaginary parts of $\alpha$ and $\alpha'$ from below by $n^{-1/4}\log n$ using the definition of the set $S$. Since $|s_n(\alpha) - \mathbb{E}s_n(\alpha)| > 2n^{-1/4}$, it follows that $|s_n(\alpha') - \mathbb{E}s_n(\alpha')| > n^{-1/4}$. Summarizing, we have shown that

$$\mathbb{P}\left(\sup_{\alpha\in S} |s_n(\alpha) - \mathbb{E}[s_n(\alpha)]| > 2n^{-1/4}\right) \leq \mathbb{P}\left(\sup_{\alpha\in\mathcal{N}} |s_n(\alpha) - \mathbb{E}[s_n(\alpha)]| > n^{-1/4}\right).$$

Thus, the claim follows from (44). □

##### S2.3.5 Hanson–Wright inequality

Recall that $z$ is a random vector in $\mathbb{R}^p$ with mean zero, and the rows of $Z$ are iid copies of $z$. In addition, $\Sigma$ is the covariance matrix of $z$. The Hanson–Wright inequality is a concentration inequality which shows that the quadratic form $z^T Az$ concentrates around its expected value, where $A$ here is an arbitrary deterministic $p \times p$ matrix. If the entries of $z$ were independent, the bound would follow from more classical results such as (Wright, 1973; Hanson *et al.*, 1971; Rudelson *et al.*, 2013). In the model under consideration the entries of $z$ are dependent, and this dependence requires different techniques to deal with the quadratic form $z^T Az$. We begin first with its expected value.

**Lemma 8.** *For a deterministic $p \times p$ matrix $A$,*

$$\frac{1}{p}\mathbb{E}[z^T A z] = \frac{1}{p}\mathrm{tr}\,(A\Sigma) = \frac{1}{p}\mathrm{tr}\,(A) + \mathcal{O}(m^{-1}\|A\|). \tag{45}$$

*Proof.* We denote $z = (z_i)_{i=1}^p$. Then

$$\mathbb{E}[z^T A z] = \sum_{i,j=1}^{p} A_{ij}\mathbb{E}[z_i z_j] = \sum_{i,j=1}^{p} A_{ij}\Sigma_{ij} = \mathrm{tr}\,(A\Sigma^T) = \mathrm{tr}\,(A\Sigma).$$

This establishes the first equality in (45).

We now turn to the second equality in (45). Recall that

$$\Sigma = \begin{bmatrix} \Sigma' & 0 \\ 0 & I_{p-m} \end{bmatrix},$$

where $\Sigma'$ is the $m \times m$ matrix defined in (32). We similarly decompose $A$ in block form as

$$A = \begin{bmatrix} A_1 & A_2 \\ A_3 & A_4 \end{bmatrix},$$

where $A_1$ is $m \times m$ and $A_4$ is $(p-m) \times (p-m)$. Then $\mathrm{tr}\,(A\Sigma) = \mathrm{tr}\,(A_1\Sigma') + \mathrm{tr}\,(A_4)$. We rewrite $\Sigma' = 2\mu(m)J + (1-\mu(m))I$, where $J$ is the all-ones matrix. So, we obtain

$$\mathrm{tr}\,(A_1\Sigma') = (1-\mu(m))\mathrm{tr}\,(A_1) + 2\mu(m)\mathrm{tr}\,(A_1 J).$$

Since $J$ is rank one, we can use (42) to bound $|\mathrm{tr}\,(A_1 J)| \leq \|A_1\|\|J\| \leq \|A\|\|J\|$. A simple computation shows that $\|J\| = m$, and using $\mu(m) = \mathcal{O}(m^{-1})$, we conclude that

$$\frac{1}{p}\mathrm{tr}\,(A_1\Sigma') = \frac{1}{p}\mathrm{tr}\,(A_1) + \mathcal{O}(m^{-1}\|A\|),$$

where we used that $m \leq p$. Putting these bounds together yields

$$\frac{1}{p}\mathrm{tr}\,(A\Sigma) = \frac{1}{p}\mathrm{tr}\,(A_1) + \frac{1}{p}\mathrm{tr}\,(A_4) + \mathcal{O}(m^{-1}\|A\|) = \frac{1}{p}\mathrm{tr}\,(A) + \mathcal{O}(m^{-1}\|A\|),$$

which completes the proof. □

Our main concentration bound is the following.

**Lemma 9.** *For a deterministic $p \times p$ complex symmetric matrix $A$ with $\|A\| = \mathcal{O}(1)$,*

$$\mathbb{E}\left|\frac{1}{p}z^T Az - \frac{1}{p}\mathbb{E}[z^T Az]\right| = o(1).$$

*Proof.* By the Cauchy–Schwarz inequality, it suffices to show that

$$\mathbb{E}\left|\frac{1}{p}z^T Az - \frac{1}{p}\mathbb{E}[z^T Az]\right|^2 = o(1).$$

Identifying the left-hand side as the variance of $\frac{1}{p}z^T Az$ and applying Lemma 8, it suffices to show that

$$\mathbb{E}\left|\frac{1}{p}z^T Az\right|^2 = \left|\frac{1}{p}\mathrm{tr}\,(A)\right|^2 + o(1).$$

We expand the left-hand side as

$$\mathbb{E}\left|\frac{1}{p}z^T Az\right|^2 = \frac{1}{p^2}\sum_{i,j,k,l=1}^{p} A_{ij}\overline{A_{kl}}\mathbb{E}[z_i z_j z_k z_l],$$

where we have denoted $z = (z_i)_{i=1}^p$. Decompose the sum as

$$\frac{1}{p^2}\sum_{i,j,k,l=1}^{p} A_{ij}\overline{A_{kl}}\mathbb{E}[z_i z_j z_k z_l] = S_1 + S_2 + S_3 + S_4,$$

where

$$S_r := \frac{1}{p^2}\sum A_{ij}\overline{A_{kl}}\mathbb{E}[z_i z_j z_k z_l]$$

and the sum is over all $1 \le i, j, k, l \le p$ such that $|\{i, j, k, l\}| = r$.

We will show that $S_2$ contains the main contribution and the remaining terms $S_r$, $r \neq 2$ can be appropriately bounded. We begin by bounding $S_3$ and $S_4$ together. Let $\Lambda$ be a $p^2 \times p^2$ matrix, indexed by $\{(i, j) : 1 \le i, j \le p\}$ and defined by

$$\Lambda_{(i,j),(k,l)} = \begin{cases} \mathbb{E}[z_i z_j z_k z_l] & \text{if } |\{i,j,k,l\}| \ge 3, \\ 0, & \text{otherwise.} \end{cases}$$

Then

$$S_3 + S_4 = \frac{1}{p^2}\sum_{i,j,k,l=1}^{p} A_{ij}\overline{A_{kl}}\Lambda_{(i,j),(k,l)}$$

can be identified as a quadratic form in the matrix $\Lambda$. Thus, we can bound (see for instance (Bobkov *et al.*, 2019))

$$|S_3 + S_4| \le \frac{1}{p^2}\|A\|_2^2\|\Lambda\|_2.$$

Trivially, by (42), we have the bound

$$\frac{1}{p}\|A\|_2^2 \le \|A\|^2. \tag{46}$$

Additionally, by assumption 33 and the fact that $m \ge cp$,

$$\frac{1}{p^2}\|\Lambda\|_2^2 = \frac{1}{p^2}\sum_{|\{i,j,k,l\}|\ge 3} |\mathbb{E}[z_i z_j z_k z_l]|^2 = o(1).$$

Thus, we conclude that $|S_3 + S_4| = o(1)$.

For $S_1$, we easily bound

$$|S_1| \le \frac{1}{p^2}\sum_{i=1}^{p} \mathbb{E}[z_i^4]|A_{ii}|^2 \le \frac{\kappa^4}{p}\|A\|^2 = o(1)$$

using assumption (1) on page 26.

It remains to control the terms from $S_2$. When $i = j$ and $k = l \ne i$, the sum in $S_2$ gives (using assumption 2 on page 26)

$$\frac{1}{p^2}\sum A_{ij}\overline{A_{kl}}\mathbb{E}[z_i z_j z_k z_l] = \left(\frac{1}{p}\mathrm{tr}\,(A)\right)^2 + o(1),$$

which is the main contribution. Thus, we need to show that the other terms in $S_2$ give negligible contribution. Indeed, we have

$$\left| S_2 - \left(\frac{1}{p}\mathrm{tr}\, A\right)^2 \right| \le C\frac{1}{p^2}\|A\|_2^2 + o(1)$$

by assumptions 2 and 3 on page 26. Here $C > 0$ is a constant. Using (46), we conclude that

$$\left| S_2 - \left(\frac{1}{p}\mathrm{tr}\, A\right)^2 \right| = o(1),$$

and the proof is complete. □

##### S2.3.6 Proof of Theorem 1

We are now in a position to complete the proof of Theorem 1. We let $z_1, \ldots, z_n$ denote the rows of $Z$ (which are iid copies of $z$). Set $\tau_n := \frac{n}{p}$ and recall that $\tau_n \to \tau \in (0, \infty)$.

By the Schur complement formula (see for instance (Bai *et al.*, 2010, Theorem A.4), it follows that

$$\begin{aligned} s_n(\alpha) &= \frac{1}{n}\mathrm{tr}\, R(\alpha) \\ &= \frac{1}{n}\sum_{k=1}^{n} \frac{1}{\frac{1}{p}z_k z_k^T - \alpha - \frac{1}{p^2}z_k Z^{(k)T} R^{(k)}(\alpha) Z^{(k)} z_k^T} = \frac{1}{n}\sum_{k=1}^{n}\frac{1}{a_k}, \end{aligned}$$

where

$$a_k := \frac{1}{p} z_k z_k^T - \alpha - \frac{1}{p^2} z_k Z^{(k)T} R^{(k)}(\alpha) Z^{(k)} z_k^T.$$

Moreover, it follows (see for example page 472 of (Bai *et al.*, 2010)) that

$$|a_k| \geq \Im(\alpha). \tag{47}$$

So far we have considered arbitrary $\alpha \in \mathbb{C}^+$, but we now restrict ourselves to

$$\alpha \in Q := \{w \in \mathbb{C}^+ : \Re(w) = 0, 1/2 \leq \Im(w) \leq 1\}.$$

We have, for any $\alpha \in Q$,

$$\begin{aligned} \left| \mathbb{E}s_n(\alpha) - \frac{1}{n}\sum_{k=1}^{n} \frac{1}{\mathbb{E}[a_k]} \right| &\leq \frac{1}{n}\sum_{k=1}^{n} \mathbb{E}\left| \frac{1}{a_k} - \frac{1}{\mathbb{E}[a_k]} \right| \\ &\leq \frac{1}{n}\sum_{k=1}^{n} \mathbb{E}\left[ \frac{|a_k - \mathbb{E}[a_k]|}{(\Im(\alpha))^2} \right] \\ &\leq 4\mathbb{E}\left[ |a_1 - \mathbb{E}[a_1]| \right], \end{aligned} \tag{48}$$

where we used (47) and the fact that $\Im(\alpha) \geq 1/2$ for any $\alpha \in Q$. In the last inequality we also used that the random variables $a_k$ are identically distributed since the rows of $Z$ are iid.

Letting

$$A := I - \frac{1}{p} {Z^{(1)}}^T R^{(1)}(\alpha) Z^{(1)},$$

we note that

$$a_1 - \mathbb{E}[a_1] = \frac{1}{p} z_1 A z_1^T - \frac{1}{p}\mathbb{E}[z_1 A z_1^T].$$

By (41), $\|A\| \leq 3$ for all $\alpha \in Q$. In addition, $A$ is independent of $z_1$. Let $\mathbb{E}_1[\cdot]$ denote the expectation with respect to only the first row $z_1$. Our goal is to now show that

$$\mathbb{E}\left[ |a_1 - \mathbb{E}[a_1]| \right] = o(1) \tag{49}$$

uniformly for $\alpha \in Q$. Indeed, using the triangle inequality and Lemma 9 (by conditioning on the matrix $A$), we see that

$$\mathbb{E}\left[|a_1 - \mathbb{E}[a_1]|\right] \leq \mathbb{E}\left[|a_1 - \mathbb{E}_1[a_1]|\right] + \mathbb{E}\left[|\mathbb{E}_1[a_1] - \mathbb{E}[a_1]|\right] = o(1) + \mathbb{E}\left[|\mathbb{E}_1[a_1] - \mathbb{E}[a_1]|\right]$$

uniformly for $\alpha \in Q$. To bound the second term we note that

$$\mathbb{E}\left[|\mathbb{E}_1[a_1] - \mathbb{E}[a_1]|\right] = \mathbb{E}\left[\left|\frac{1}{p}\text{tr}\, A - \mathbb{E}\left[\frac{1}{p}\text{tr}\, A\right]\right|\right] + o(1)$$

due to Lemma 8 and Fubini's theorem. By the cyclic property of the trace

$$|\text{tr}\, A - \mathbb{E}[\text{tr}\, A]| = |\alpha| \left|\text{tr}\, R^{(1)}(\alpha) - \mathbb{E}[\text{tr}\, R^{(1)}(\alpha)]\right|$$

since

$$\frac{1}{p} R^{(1)} Z^{(1)} {Z^{(1)}}^T = R^{(1)} \left(\frac{1}{p} Z^{(1)} {Z^{(1)}}^T - \alpha I + \alpha I\right) = I + \alpha R^{(1)}. \tag{50}$$

So by Lemmas 2 and 7, we see that

$$\mathbb{E}\left[|\mathbb{E}_1[a_1] - \mathbb{E}[a_1]|\right] \leq \tau_n \mathbb{E}\left[|s_n(\alpha) - \mathbb{E}[s_n(\alpha)]|\right] + o(1) = o(1)$$

uniformly for $\alpha \in Q$. Putting together the bounds from above, we obtain (49).

Combining (49) with (48), we conclude that

$$\left|\mathbb{E}[s_n(\alpha)] - \frac{1}{\mathbb{E}[a_1]}\right| = \left|\mathbb{E}[s_n(\alpha)] - \frac{1}{n}\sum_{k=1}^{n} \frac{1}{\mathbb{E}[a_k]}\right| = o(1) \tag{51}$$

uniformly for $\alpha \in Q$. Here we have again exploited the fact that the random variables $a_k$ are identically distributed.

In view of Lemma 8 and Fubini's theorem,

$$\frac{1}{p}\mathbb{E}[z_1 A z_1^T] = \frac{1}{p}\mathbb{E}[\text{tr}\, A] + o(1)$$

uniformly for $\alpha \in Q$. Using Lemma 2, the cyclic property of the trace, and (50), we obtain

$$\begin{aligned}
\frac{1}{p}\mathbb{E}[z_1 A z_1^T] &= 1 - \frac{1}{p^2}\mathbb{E}\left[\text{tr}\, R^{(1)}(\alpha) Z^{(1)} {Z^{(1)}}^T\right] + o(1) \\
&= 1 - \tau_n - \frac{\alpha}{p}\mathbb{E}[\text{tr}\, R^{(1)}(\alpha)] + o(1) \\
&= 1 - \tau_n - \frac{\alpha}{p}\mathbb{E}[\text{tr}\, R(\alpha)] + o(1) \\
&= 1 - \tau_n - \alpha\tau_n \mathbb{E}[s_n(\alpha)] + o(1)
\end{aligned}$$

uniformly for $\alpha \in Q$. Thus, since $\tau_n \to \tau$,

$$\mathbb{E}[a_1] = 1 - \alpha - \tau - \alpha\tau\mathbb{E}[s_n(\alpha)] + o(1)$$

uniformly for $\alpha \in Q$. Returning to (51), we can express the Stieltjes transform of the empirical spectral measure as

$$\mathbb{E}[s_n(\alpha)] = \frac{1}{\mathbb{E}[a_1]} + o(1) = \frac{1}{1-\alpha-\tau-\alpha\tau\mathbb{E}[s_n(\alpha)]} + o(1)$$

uniformly for $\alpha \in Q$, where we used (47) and Lemma 3 to remove the $o(1)$ error term from the denominator. We are now in a position to apply Lemma 4. Indeed, Lemma 4 gives

$$\sup_{\alpha\in Q} |\mathbb{E}[s_n(\alpha)] - s(\alpha)| = o(1).$$

Combined with Lemma 7 and the Borel–Cantelli lemma, this implies that

$$\sup_{\alpha\in Q} |s_n(\alpha) - s(\alpha)| \to 0$$

almost surely as $n \to \infty$. Since $s_n$ and $s$ are analytic in $\mathbb{C}^+$ and satisfy $|s_n(\alpha)| \leq \frac{1}{\Im(\alpha)}$ and $|s(\alpha)| \leq \frac{1}{\Im(\alpha)}$ (by Lemma 3), Vitali's convergence theorem (see page 168 in (Tao, 2011) or (Bai *et al.*, 2010, Lemma 2.14)) implies that almost surely

$$s_n(\alpha) \to s(\alpha)$$

as $n \to \infty$ for each $\alpha \in \mathbb{C}^+$. In other words, we have shown that the Stieltjes transform of the ESD of $\frac{1}{p}ZZ^T$ converges almost surely to the Stieltjes transform of $F_\tau^{\mathrm{MP}}$. This implies the almost sure convergence of the ESD by (Bai *et al.*, 2010, Theorem B.9). The proof of Theorem 1 is complete.

#### S2.4 The limiting spectral distribution of the genetic relatedness matrix

We now demonstrate that the ESD of the GRM $m^{-1}ZZ^T$ converges almost surely to $F_\tau^{\mathrm{MP}}$. All that remains is to show that the elements of $Z$ conform to the assumptions needed for Theorem 1. We restate those assumptions here: There exists a constant $\kappa > 0$ so that $\sup_{1\leq i\leq m} |z_i| \leq \kappa$ with probability 1[5]. $\mathbb{E}[z_i^2 z_j^2] = 1 + o(1)$ uniformly for all distinct $i, j$.

1. $\mathbb{E}[z_i^3 z_j] = o(1)$ uniformly for all distinct $i, j$.
2. One has

$$\mathbb{E}[z_i z_j z_k z_l] = \begin{cases} o\left(\frac{1}{m}\right), & \text{if } |\{i,j,k,l\}| = 4, \\ o\left(\frac{1}{\sqrt{m}}\right), & \text{if } |\{i,j,k,l\}| = 3, \end{cases} \tag{52}$$

   uniformly in $i, j, k, l$.

[5]With a bit of extra work, this can be generalized to the case where $\sup_{1\leq i\leq m} |z_i| = o(\log(m))$.

First, S2.4 is ensured by choosing a minor allele frequency cutoff $\delta > 0$ independent of sample size, as is common practice. Because each $\mathcal{Z} = 2^{-1/2}(\mathcal{G}_1 + \mathcal{G}_2)$ is the sum of two standardized Bernoullis, we then have

$$\sup_{1 \leq i \leq m} |\mathcal{Z}_i| \leq \sqrt{2} \frac{1-\delta}{\sqrt{\delta(1-\delta)}}.$$

Thus, setting $\delta = 2(2+\kappa^2)^{-1}$ ensures $\sup_{1 \leq i \leq m} |\mathcal{Z}_i| \leq \kappa$.

###### S2.4.1 From diploid genotypes to haploid genotypes

In what follows we establish that Theorem 1 can be applied directly to the GRM under assortative mating. For a given collection of diploid sites indexed $\alpha_1, \ldots, \alpha_N$, denote

$$\mathcal{S}_{\alpha_1|\cdots|\alpha_N} := \sup_{\{\iota_1, \ldots, \iota_N\}} \left| \mathbb{E}\left[ \prod_{i=1}^{N} \mathcal{G}_{\iota_i}^{\alpha_i} \right] \right|.$$

Then $|\mathbb{E}[\mathcal{Z}_i^2 \mathcal{Z}_{j \neq i}^2]|$ is a homogeneous fourth degree polynomial bounded by

$$|\mathbb{E}[\mathcal{Z}_i^2 \mathcal{Z}_{j \neq i}^2]| \leq \mathcal{S}_{2|2} + 2\mathcal{S}_{1|1|2} + \mathcal{S}_{1|1|1|1},$$

Likewise, we have

$$\begin{aligned} |\mathbb{E}[\mathcal{Z}_i^3 \mathcal{Z}_{j \neq i}]| &\leq \mathcal{S}_{1|3} + 3\mathcal{S}_{1|1|2}, \\ |\mathbb{E}[\mathcal{Z}_i^2 \mathcal{Z}_{j \neq i} \mathcal{Z}_{k \neq i,j}]| &\leq 2\mathcal{S}_{1|1|2} + 2\mathcal{S}_{1|1|1|1}, \\ |\mathbb{E}[\mathcal{Z}_i \mathcal{Z}_{j \neq i} \mathcal{Z}_{k \neq i,j} \mathcal{Z}_{l \neq i,j.k}]| &\leq 4\mathcal{S}_{1|1|1|1}. \end{aligned}$$

Thus, we need simply need to establish equivalent bounds on the moments of haploid genotypes:

1. $\mathbb{E}[\mathcal{G}_i \mathcal{G}_{j \neq i} \mathcal{G}_{k \neq i,j} \mathcal{G}_{l \neq i,j,k}] = o\left(\frac{1}{m}\right)$,
2. $\mathbb{E}[\mathcal{G}_i^2 \mathcal{G}_{j \neq i} \mathcal{G}_{k \neq i,j}] = o\left(\frac{1}{\sqrt{m}}\right)$,
3. $\mathbb{E}[\mathcal{G}_i^3 \mathcal{G}_{j \neq i}] = o(1)$,
4. $\mathbb{E}[\mathcal{G}_i^2 \mathcal{G}_{j \neq i}^2] = 1 + o(1)$,

all which we demonstrated in Section S2.1.

### S3 Impact of assortative mating on marker-based heritability estimators

#### S3.1 Haseman-Elston regression

Denote the lower triangular components of the phenotypic and genotypic sample covariance matrices as

$$\mathcal{Y} = \text{vec}\,(\{\sigma_y^{-2} yy^T\}_{i,j:i<j}), \qquad \mathcal{S} = \text{vec}\,(\{p^{-1}\mathcal{Z}\mathcal{Z}^T\}_{i,j:i<j}).$$

The HE regression heritability is estimator is obtained by regressing $\mathcal{Y}$ on to $\mathcal{S}$:

$$\hat{h}^2_{\text{HE}} = \frac{\widehat{Cov}(\mathcal{Y}, \mathcal{S})}{\widehat{Var}(\mathcal{S})}.$$

Under exchangeable, loci, the same is true for each of the haploid standardized genotypes at casual variants, which we denote $\{\mathcal{G}_k\}_{k=1}^{2m}$, where $Z_k = 2^{-1/2}(\mathcal{G}_{2k} + \mathcal{G}_{2k+1})$, $k = 1, \ldots, m$. We assume that all genotypes at non-causal variants are independent.

For each allelic effect $u_k$ define $\phi_k = \sqrt{r}\nu_{k,\infty}$ such that each correlation between haploid variants is $\mu_{kl,\infty} = \phi_k\phi_l$. The variance/covariance matrix of the standardized diploid genetic effects is thus

$$\begin{aligned} \mathbb{E}\left(m^{-1}\mathcal{Z}^T\mathcal{Z}\right) &= \text{diag}\{1 - \phi_k^2\}_{k=1}^m + 2\phi\phi^T \\ &:= \Upsilon_\infty. \end{aligned}$$

Now consider the element $\mathcal{S}_{ij}$ of $\mathcal{S}$ corresponding to the average similarity at $m$ diploid loci among the $i^{th}$ and $j^{th}$ unrelated individuals:

$$\mathcal{S}_{ij} = \frac{1}{m}\sum_{k=1}^m \mathcal{Z}_{i,k}\mathcal{Z}_{j,k}.$$

Because these individuals are unrelated, $\mathcal{Z}_{i,k}$ and $\mathcal{Z}_{j,l}$ are independent for all $k, l$ and thus $\mathbb{E}[\mathcal{S}_{ij}] = 0$. The variance is then simply computed as

$$\begin{aligned} Var\,(\mathcal{S}_{ij}) &= m^{-2}\sum_{k=1}^m\sum_{l=1}^m \mathbb{E}\,[\mathcal{Z}_{i,k}\mathcal{Z}_{j,k}\mathcal{Z}_{i,l}\mathcal{Z}_{j,l}] \\ &= m^{-2}\text{tr}\,[\Upsilon_\infty^2]. \end{aligned}$$

Turning our attention to $\mathcal{Y}_{ij}$ and marginalizing over independent error terms, we see that

$$\hat{\sigma}^2_{y,\infty}\mathcal{Y}_{ij} = \sum_{k,l=1}^{m} \mathcal{Z}_{i,k}\mathcal{Z}_{j,l}u_k u_l,$$

has zero expectation. The covariance between $\mathcal{Y}_{ij}$ and $\mathcal{S}_{ij}$ is then

$$Cov(\mathcal{Y}_{ij}, \mathcal{S}_{ij}) = \hat{\sigma}^{-2}_{y,\infty} m^{-1} u^T \Upsilon^2_\infty u.$$

Thus, the expected value of the HE estimator is computed

$$\begin{aligned}
\mathbb{E}[\hat{h}^2_{\text{HE}}] &= \frac{Cov(\mathcal{Y}, \mathcal{S})}{Var(\mathcal{S})} \\
&= \frac{m^{-1} u^T \Upsilon^2_\infty u}{\sigma^2_{y,\infty} m^{-2} \text{tr}\left[\Upsilon^2_\infty\right]} \left(\frac{\sigma^2_{g,\infty}}{\sigma^2_{g,\infty}}\right) \\
&= \frac{m \cdot u^T \Upsilon^2_\infty u}{\sigma^2_{g,\infty} \cdot \text{tr}\left[\Upsilon^2_\infty\right]} h^2_\infty.
\end{aligned}$$

Note that when $\Upsilon_\infty = I$, as is expected under random mating, we simply have $\mathbb{E}[\hat{h}^2_{\text{HE}}] = h^2_\infty = h^2_0$; i.e. HE regression is unbiased under random mating. We can further simplify the above if we assume exchangeable loci (i.e., for all $k, l$, $u_k u_l = m^{-1}\sigma^2_{g,0}$). In this case, define $\phi \in (-1, 1)^m$ by $\phi_k \equiv r^{1/2}\nu_\infty := \phi$ for all $k$. We can then write $\Upsilon_\infty = D + 2\phi\phi^T$, where $D = \text{diag}\{1 - \phi^2_k\}^m_{k=1}$. We compute $u^T \Upsilon^2_\infty u$ by first noting that

$$\begin{aligned}
u^T \Upsilon^2_\infty u =& u^T (D^2 + 2\phi\phi^T D + 2D\phi\phi^T + 4\phi\phi^T\phi\phi^T) u \\
=& u^T D^2 u + 4u^T\phi\phi^T D u + 4\phi^T\phi u^T\phi\phi^T u.
\end{aligned}$$

Computing the above terms individually yields

$$\begin{aligned}
u^T D^2 u &= \sigma^2_{g,0}(1 - \phi^2)^2, \\
u^T \phi\phi^T D u &= m\sigma^2_{g,0}\phi^2(1 - \phi^2), \\
\phi^T\phi u^T \phi\phi^T u &= m^2\sigma^2_{g,0}\phi^4, \\
\implies Cov(\mathcal{Y}_{ij}, \mathcal{S}_{ij}) &= \hat{\sigma}^{-2}_{y,\infty} m^{-1}\sigma^2_{g,0}\left(1 - \phi^2 + 2m\phi^2\right)^2,
\end{aligned}$$

Likewise, we compute the denominator

$$Var\left(\mathcal{S}_{ij}\right) = m^{-1}\left(1 + 2\phi^2 + 4m\phi^4\right).$$

Thus, under exchangeable loci,

$$\mathbb{E}[\hat{h}^2_{\text{HE}}] = \frac{\sigma^2_{g,0}\left(1-\phi^2+2m\phi^2\right)^2}{\sigma^2_{g,\infty}\left(1+2\phi^2+4m\phi^4\right)} h^2_\infty$$

where $\phi^2 = m\sigma^2_{y,\infty}(8r\sigma^2_{g,0})^{-1}\left(\sqrt{4r\sigma^2_{g,0}m^{-1}\sigma^{-2}_{y,\infty}+(1-r_{g,\infty})^2}-(1-r_{g,\infty})\right)^2$. Substituting in panmictic parameters and taking the limit as $m \to \infty$ yields

$$\mathbb{E}[\hat{h}^2_{\text{HE}}] \approx \frac{\sigma^2_{g,0}}{\sigma^2_{g,\infty}} h^2_\infty = \frac{h^2_\infty}{1-r_{g,\infty}}.$$

This gives us a way to estimate the panmictic and equilibrium heritabilities as a function of the HE estimator when $r$ is known:

$$\hat{h}^2_0 := \mathbb{E}[h^2_0|\hat{h}^2_{\text{HE}}] = \frac{\hat{h}^2_{\text{HE}}}{1+2r\hat{h}^2_{\text{HE}}+r(r-1)\hat{h}^4_{\text{HE}}},$$
$$\hat{h}^2_\infty := \mathbb{E}[h^2_\infty|\hat{h}^2_{\text{HE}}] = \frac{\hat{h}^2_{\text{HE}}}{1+r\hat{h}^2_{\text{HE}}}.$$

We can then employ the delta method to approximate the standard errors:

$$\text{se}\left[\hat{h}^2_0\right] \approx \text{se}\left[\hat{h}^2_{\text{HE}}\right] \frac{1+r\hat{h}^4_{\text{HE}}-r^2\hat{h}^4_{\text{HE}}}{\left(1-r(\hat{h}^2_{\text{HE}}-2)\hat{h}^2_{\text{HE}}+r^2\hat{h}^4_{\text{HE}}\right)^2},$$
$$\text{se}\left[\hat{h}^2_\infty\right] \approx \frac{\text{se}\left[\hat{h}^2_{\text{HE}}\right]}{1+r\hat{h}^2_{\text{HE}}}.$$

#### S3.2 Residual maximum likelihood

##### S3.2.1 Notation and problem statement

Consider the model

$$y = X\beta + \tilde{Z}u + e,$$
$$u \overset{i.i.d.}{\sim} \mathcal{N}(0,\sigma^2_g), \qquad e \overset{i.i.d.}{\sim} \mathcal{N}(0,\sigma^2_e),$$

or, equivalently,

$$(y|Z) \sim \mathcal{MVN}(X\beta, \sigma^2_e V_{\mathring{\gamma}}),$$
$$V_{\mathring{\gamma}} = \mathring{\gamma}\tilde{Z}\tilde{Z}^T + I_n, \qquad \mathring{\gamma} = \sigma^2_g/\sigma^2_e.$$

Above, $Z \in \mathbb{R}^{n \times p}$ and $u \in \mathbb{R}^p$ are independent of $e \in \mathbb{R}^n$ and $X \in \mathbb{R}^{n \times c}$, $\beta \in \mathbb{R}^c$ are deterministic. The variance components $\sigma_g^2$, $\sigma_e^2$, and thus their ratio $\mathring{\gamma}$, are strictly positive. $\tilde{Z}\tilde{Z}^T = p^{-1}ZZ^T$, the matrix we've been concerned with, is sub-Gaussian with independent rows and, by our results from Section S2, is such that its empirical spectral density $F^{\tilde{Z}\tilde{Z}^T} \overset{\text{a.s.}}{\to} F_\tau^{\text{MP}}$, where $\tau = n/p$. For simplicity, we assume $m = p$ (all variants are causal) and $Cov(Z_{.k}, Z_{.l}) \equiv \mu = \mathcal{O}(m^{-1})$ for all $k \neq l$ (exchangeable loci), though our results will hold as long as $m/p \to c \in (0, 1]$ and $\sup_{k \neq l} Cov(Z_{.k}, Z_{.l}) = \mathcal{O}(m^{-1})$. In what follows, $\mathring{\gamma}$ will denote true parameter value, $\hat{\gamma}$ the REML estimator, and $\gamma > 0$ an arbitrary value.

Let $A^T : \mathbb{R}^n \to (\operatorname{col} X)^\perp$ such that $A^T X = 0$, $A^T A = I_{n-c}$ and define the following quantities for any value of $\gamma > 0$:

- $\tilde{y} = y - X\beta$, the mean-deviated outcome vector
- $\Sigma_\gamma = A^T V_\gamma A = I_{n-c} + \gamma A^T \tilde{Z}\tilde{Z}^T A$, the transformed marginal covariance
- $\zeta = A^T \tilde{Z}$, the transformed standardized genotypes
- $P_\gamma = A\Sigma_\gamma^{-1} A^T$, the Schur complement of $X^T V_\gamma^{-1} X$ in $\begin{pmatrix} V_\gamma^{-1} & X^T V_\gamma^{-1} \\ V_\gamma^{-1} X & X^T V_\gamma^{-1} X \end{pmatrix}$
- $\Delta(\gamma) = \sigma_e^{-2} y^T \left( \frac{P_\gamma \tilde{Z}\tilde{Z}^T P_\gamma}{\operatorname{tr}[P_\gamma \tilde{Z}\tilde{Z}^T]} - \frac{P_\gamma^2}{\operatorname{tr}[P_\gamma]} \right) y = \sigma_e^{-2} \tilde{y}^T \left( \frac{P_\gamma \tilde{Z}\tilde{Z}^T P_\gamma}{\operatorname{tr}[P_\gamma \tilde{Z}\tilde{Z}^T]} - \frac{P_\gamma^2}{\operatorname{tr}[P_\gamma]} \right) \tilde{y}$, the "stationarity function".

The REML estimator $\hat{\gamma}$ satisfies

$$\sigma_e^{-2} \frac{y^T P_{\hat{\gamma}} \tilde{Z}\tilde{Z}^T P_{\hat{\gamma}} y}{\operatorname{tr}[P_{\hat{\gamma}} \tilde{Z}\tilde{Z}^T]} = \sigma_e^{-2} \frac{y^T P_{\hat{\gamma}}^2 y}{\operatorname{tr}[P_{\hat{\gamma}}]}, \tag{53}$$

or, equivalently, it is the root of the stationarity function $\Delta(\hat{\gamma}) = 0$. Choosing $\hat{\gamma}$ such that $\Delta(\hat{\gamma}) = 0$ and setting $\hat{\sigma}_e^2 = \frac{y^T P_{\hat{\gamma}}^2 y}{\operatorname{tr}[P_{\hat{\gamma}}]}$, $\hat{\sigma}_g^2 = \hat{\sigma}_e^2 \hat{\gamma}$ yields the variance component estimators that maximize the likelihood of

$$(A^T y | Z) \sim \mathcal{MVN}(0, \sigma_e^2 \Sigma_{\mathring{\gamma}}).$$

Our goal is to demonstrate that $\hat{\gamma} \overset{P}{\to} \mathring{\gamma}$ as $n, p \to \infty$, $n/p \to \tau$. For simplicity, we address the exchangeable loci case where each $u_k \equiv \pm m^{-1/2} \sigma_{g,0}$ for $k = 1, \ldots, m$, though a similar analysis is possible under a random SNP effect model given the moments of $u|Z$.

##### S3.2.2 The limiting spectral distribution of $\zeta\zeta^T = A^T \tilde{Z}\tilde{Z}^T A$

From your previous results, we know that the ESD of the $n \times p$ matrix $\tilde{Z} = p^{1/2} Z$ is such that $F^{\tilde{Z}\tilde{Z}^T} \overset{\text{a.s.}}{\to} F_\tau^{\text{MP}}$. Here we show that the ESD of the $(n - c) \times p$ matrix $\zeta = A^T \tilde{Z}$, is such that $F^{\zeta\zeta^T} \overset{\text{a.s.}}{\to} F_\tau^{\text{MP}}$ as well. We first establish some intermediate results.

**Lemma 10.** *Suppose $Q$ is a Hermitian matrix. If each eigenvalue of $Q$ is either 0 or 1, then $Q$ is an orthogonal projection.*

*Proof.* By supposition $Q$ is Hermitian, so it suffices to show that $Q^2 = Q$. By the spectral theorem, we can write

$$Q = U\Lambda U^*,$$

where $U$ is a unitary matrix and $\Lambda$ is a diagonal matrix whose diagonal entries are the eigenvalues of $Q$. This implies that $\Lambda^2 = \Lambda$ since the eigenvalues of $Q$ can only be 0 or 1 by assumption. It follows that

$$Q^2 = U\Lambda U^* U\Lambda U^* = U\Lambda^2 U^* = Q,$$

and the proof is complete. $\square$

**Lemma 11.** *Suppose $Q$ is an $n \times n$ orthogonal projection matrix. If zero is an eigenvalue of $Q$ with multiplicity $m$, then* $\operatorname{rank}(I_n - Q) = m$.

*Proof.* By the spectral theorem, we can decompose

$$Q = U\Lambda U^*,$$

where $U$ is an $n \times n$ unitary matrix and $\Lambda$ is an $n \times n$ diagonal matrix whose diagonal entries are the eigenvalues of $P$ (and so must be either 0 or 1). Hence,

$$I_n - Q = U(I_n - \Lambda)U^*.$$

Since $U$ and $U^*$ have full rank, it follows that

$$\operatorname{rank}(I_n - Q) = \operatorname{rank}(I_n - \Lambda).$$

Since $I_n - \Lambda$ is a diagonal Hermitian matrix, it follows that the rank of $I_n - \Lambda$ is just the number of nonzero diagonal entries of $I_n - \Lambda$. In other words, the rank of $I_n - \Lambda$ is simply the number of diagonal entries of $\Lambda$ that are zero. $\square$

**Lemma 12.** *Let $A$ be an $n \times m$ matrix and $B$ an $m \times n$ matrix. Then the non-trivial eigenvalues of $AB$ are the same as the non-trivial eigenvalues of $BA$ (counting algebraic multiplicity).*

*Proof.* The lemma will follow as an application of Sylvester's determinant identity (also called the Weinstein-Aronszajn identity): if $C$ and $D$ are matrices of size $n \times m$ and $m \times n$ respectively, then

$$\det(I_n + CD) = \det(I_m + DC).$$

This identity can be found on pg. 271 of (Pozrikidis, 2014).

We now use Sylvester's determinant identity to complete the proof of the lemma. Assume without loss of generality that $n \geq m$. Then, for $z \neq 0$,

$$\det(zI_n - AB) = z^n \det(I_n - z^{-1}AB) = z^n \det(I_m - z^{-1}BA) = z^{n-m} \det(zI_m - BA).$$

We conclude that $AB$ and $BA$ have the same characteristic polynomials up to a factor of $z^{n-m}$. The factor $z^{n-m}$ corresponds to the trivial eigenvalues at zero. □

We now return to the problem of determining the limiting spectral distribution of $F^{\zeta\zeta^T}$.

**Lemma 13.** *If $A$ is an $n \times (n-c)$ matrix such that $A^TA = I_{n-c}$, then*

$$\|F^{p^{-1}A^TZZ^TA} - F^{p^{-1}ZZ^T}\| \leq 3\frac{c}{n}.$$

*Proof.* The eigenvalues of $A^TA$ are all one. By Lemma 12, we conclude that $AA^T$ has $n-c$ eigenvalues that are one and $c$ eigenvalues that are zero. It follows from Lemma 10 that $P := AA^T$ is an orthogonal projection matrix (since it is clearly Hermitian).

We now consider the eigenvalues of $\frac{1}{p}A^TZZ^TA$. By Lemma 12, the eigenvalues of $\frac{1}{p}A^TZZ^TA$ are the same as the eigenvalues of $\frac{1}{p}AA^TZZ^T = \frac{1}{p}QZZ^T$, except the latter matrix has $c$ additional zero eigenvalues. Hence, it follows that

$$\|F^{p^{-1}A^TZZ^TA} - F^{p^{-1}PZZ^T}\| \leq 2\frac{c}{n}.$$

Since $Q = Q^2 = Q^TQ$, by another application of Lemma 12, the eigenvalues of $\frac{1}{p}QZZ^T$ are the same as the eigenvalues of $\frac{1}{p}QZZ^TQ^T$, counting multiplicity (this latter matrix is nicer to work with because it is Hermitian). In other words,

$$\|F^{p^{-1}QZZ^T} - F^{p^{-1}QZZ^TQ^T}\| = 0.$$

To complete the proof, we now apply Theorem A.44 from (Bai *et al.*, 2010), which yields

$$\|F^{p^{-1}QZZ^TQ^T} - F^{p^{-1}ZZ^T}\| \leq \frac{1}{n}\text{rank}\,(QZ - Z).$$

By properties of the rank,

$$\text{rank}\,(QZ - Z) = \text{rank}\,((Q - I_n)Z) \leq \text{rank}\,(Q - I_n).$$

Finally, by Lemma 11, we see that $\text{rank}\,(Q - I_n) = c$. Therefore, putting together the previous bounds (and applying the triangle inequality), we conclude that

$$\|F^{p^{-1}A^TZZ^TA} - F^{p^{-1}ZZ^T}\| \leq 3\frac{c}{n}.$$

□

Finally, combining Theorem 1 with the above lemma immediately yields $F^{\zeta\zeta^T} \overset{\text{a.s.}}{\to} F_\tau^{\text{MP}}$.

###### S3.2.3 Spectral functions of $\zeta\zeta^T$

Here we introduce some notation that will simplify future computations. For fixed $\gamma, \tau > 0$, and for non-negative integers $k, l$ such that $k \leq l$, denote the continuous bounded functions

$$\begin{aligned}
\psi_{(k,l)} :& [0,\infty) \to [0,\infty), \\
\psi_{(k,l)} :& x \mapsto \frac{x^k}{(\gamma x + 1)^l}.
\end{aligned}$$

Likewise, define their integrals with respect to the MP-law $F_\tau^{MP}$, $\tau > 0$ by

$$\begin{aligned}
\Psi_{(j,l)} &= \int \psi_{(k,l)} dF_\tau^{MP} \\
&= \frac{1}{2\pi\tau} \int_{a_\tau}^{b_\tau} \psi_{(k,l)}(x) x^{-1} \sqrt{(x-a_\tau)(b_\tau - x)}\, dx,
\end{aligned}$$

where $a_\tau = (1-\sqrt{\tau})^2$, $b_\tau = (1+\sqrt{\tau})^2$. Note that $0 < \Psi_{(k,l)} < \infty$.

###### S3.2.4 Conditional expectation of the stationarity function

Recall that $y = \tilde{Z}u + e$, where $u$ is deterministic with $u^Tu = \sigma_{g,0}^2$, $\max_k |u_k| = \alpha m^{-1}$ for some $\alpha$ independent of $m$ and $e \overset{i.i.d.}{\sim} \mathcal{N}(0, \sigma_e^2)$, and, additionally, that $\zeta = A^T\tilde{Z}$, $P_\gamma = A\Sigma_\gamma^{-1}A^T$, $A^TA = I$, and $\mathring{\gamma} = \sigma_{g,0}^2/\sigma_e^2$. We consider the conditional expectation of $\Delta(\gamma)$ given $Z$:

$$\begin{aligned}
\mathbb{E}[\Delta(\gamma)|Z] &= \sigma_e^{-2} u^T \left( \frac{\Sigma_\gamma^{-1}\zeta\zeta^T\Sigma_\gamma^{-1}\zeta\zeta^T}{\text{tr}\,[\Sigma_\gamma^{-1}\zeta\zeta^T]} - \frac{\Sigma_\gamma^{-2}\zeta\zeta^T}{\text{tr}\,[\Sigma_\gamma^{-1}]} \right) u + \left( \frac{\text{tr}\,[\Sigma_\gamma^{-1}\zeta\zeta^T\Sigma_\gamma^{-1}]}{\text{tr}\,[\Sigma_\gamma^{-1}\zeta\zeta^T]} - \frac{\text{tr}\,[\Sigma_\gamma^{-2}]}{\text{tr}\,[\Sigma_\gamma^{-1}]} \right) \\
&= \sigma_e^{-2} \text{tr} \left[ \left( \frac{\Sigma_\gamma^{-1}\zeta\zeta^T\Sigma_\gamma^{-1}\zeta\zeta^T}{\text{tr}\,[\Sigma_\gamma^{-1}\zeta\zeta^T]} - \frac{\Sigma_\gamma^{-2}\zeta\zeta^T}{\text{tr}\,[\Sigma_\gamma^{-1}]} \right) uu^T \right] + \left( \frac{\text{tr}\,[\Sigma_\gamma^{-2}\zeta\zeta^T]}{\text{tr}\,[\Sigma_\gamma^{-1}\zeta\zeta^T]} - \frac{\text{tr}\,[\Sigma_\gamma^{-2}]}{\text{tr}\,[\Sigma_\gamma^{-1}]} \right). \qquad (54)
\end{aligned}$$

Write the spectral decomposition $\zeta\zeta^T = Q\Lambda Q^T$, such that $\Sigma_\gamma = Q(I + \gamma\Lambda)Q^T$, and recall that $u^Tu = \sigma_{g,0}^2$.

Under our exchangeable loci assumption, we have

$$\mathbb{E}[\Delta(\gamma)|Z] = \mathring{\gamma} \left( \frac{\mathrm{tr}\,[(I+\gamma\Lambda)^{-1}\Lambda(I+\gamma\Lambda)^{-1}\Lambda]}{\mathrm{tr}\,[\Sigma_\gamma^{-1}\zeta\zeta^T]} - \frac{\mathrm{tr}\,[(I+\gamma\Lambda)^{-2}\Lambda]}{\mathrm{tr}\,[\Sigma_\gamma^{-1}]} \right) + \left( \frac{\mathrm{tr}\,[\Sigma_\gamma^{-2}\zeta\zeta^T]}{\mathrm{tr}\,[\Sigma_\gamma^{-1}\zeta\zeta^T]} - \frac{\mathrm{tr}\,[\Sigma_\gamma^{-2}]}{\mathrm{tr}\,[\Sigma_\gamma^{-1}]} \right). \tag{55}$$

Thus, we can write $\Delta(\gamma)$ exclusively in terms of the true parameter value $\mathring{\gamma}$ and spectral functions with known limits:

$$\mathbb{E}[\Delta(\gamma)|Z] \xrightarrow{P} \mathring{\gamma} \left( \frac{\Psi_{(2,2)}}{\Psi_{(1,1)}} - \frac{\Psi_{(1,2)}}{\Psi_{(0,1)}} \right) + \left( \frac{\Psi_{(1,2)}}{\Psi_{(1,1)}} - \frac{\Psi_{(0,2)}}{\Psi_{(0,1)}} \right) := h(\gamma).$$

Let $w = \mathring{\gamma}/\gamma$. Using a computer algebra system to compute the above integrals and defining $\chi_\gamma = \sqrt{\gamma^2(\tau-1)^2 + 2\gamma(\tau+1) + 1}$, the above quantity can be written as

$$\begin{aligned}
& w\gamma \left( \frac{\Psi_{(2,2)}}{\Psi_{(1,1)}} - \frac{\Psi_{(1,2)}}{\Psi_{(0,1)}} \right) + \left( \frac{\Psi_{(1,2)}}{\Psi_{(1,1)}} - \frac{\Psi_{(0,2)}}{\Psi_{(0,1)}} \right) \\
=& (4\chi_\gamma)^{-1} \Bigg( \frac{4\gamma\left(-(\sqrt{\tau}+1)\,|1-\sqrt{\tau}\,|\chi_\gamma+\gamma(\tau-1)^2+\tau+1\right)}{\gamma(\sqrt{\tau}+1)\,|1-\sqrt{\tau}\,|-\chi_\gamma+1} + 4 - \frac{4w(-\chi_\gamma+\gamma\tau+\gamma+1)^2}{(-\chi_\gamma+\gamma\tau+\gamma+1)(-\gamma(\sqrt{\tau}+1)\,|1-\sqrt{\tau}\,|+\chi_\gamma-1)} \\
& \quad + \frac{w}{-\chi_\gamma+\gamma\tau+\gamma+1}\left(2\gamma(\sqrt{\tau}-1)^2\left(\chi_\gamma-2\gamma(\sqrt{\tau}+1)^2-3\right)+2\gamma(\sqrt{\tau}+1)^2(\chi_\gamma-3)+8(\chi_\gamma-1)\right)\Bigg) \\
:=& \frac{C_1 + wC_2}{4\chi_\gamma}, \quad \text{where } C_2/C_1 = -1,
\end{aligned}$$

where $h$ is strictly monotone in $\gamma$ and $h(\gamma) = 0 \iff \mathring{\gamma}/\gamma = 1$.

Under the more general case, dropping the assumption of exchangeable loci, we can use the following lemma to demonstrate that $\mathbb{E}[\Delta(\gamma)|Z] \xrightarrow{P} \tilde{h}(\gamma)$ with $|\tilde{h}(\gamma)| \leq |h(\gamma)|$:

**Lemma 14.** *Let $A = A^T \in \mathbb{R}^{m\times m}$, $B = B^T \succeq 0$. Then*

$$0 \leq \mathrm{tr}\,[AB] \leq \mathrm{tr}\,[B]\lambda_{\max}[A].$$

*Proof.* This is an immediate corollary of Theorem 1 in (Fang *et al.*, 1994). ☐

Denote the rank one matrix $A = uu^T$ and observe that

$$\sigma_e^{-2}Au = \sigma_e^{-2}u^Tuu = \sigma_e^{-2}\sigma_{g,0}^2 u = \mathring{\gamma}u.$$

Thus $\sigma_{g,0}^2$ is the only non-zero eigenvalue of $A$ and $\sigma_e^{-2}\mathrm{tr}\,[AB] \leq \mathring{\gamma}\mathrm{tr}\,[B]$. All of the matrices in (13) (besides the outer product $uu^T$) are symmetric positive definite, and we have

$$\mathbb{E}[\Delta(\gamma)|Z] \leq \mathring{\gamma} \left( \frac{\mathrm{tr}\,\Sigma_\gamma^{-1}\zeta\zeta^T\Sigma_\gamma^{-1}\zeta\zeta^T}{\mathrm{tr}\,[\Sigma_\gamma^{-1}\zeta\zeta^T]} - \frac{\mathrm{tr}\,\Sigma_\gamma^{-2}\zeta\zeta^T}{\mathrm{tr}\,[\Sigma_\gamma^{-1}]} \right) + \left( \frac{\mathrm{tr}\,[\Sigma_\gamma^{-2}\zeta\zeta^T]}{\mathrm{tr}\,[\Sigma_\gamma^{-1}\zeta\zeta^T]} - \frac{\mathrm{tr}\,[\Sigma_\gamma^{-2}]}{\mathrm{tr}\,[\Sigma_\gamma^{-1}]} \right).$$

In what follows, we will assume exchangeable loci, though the above inequality can be used to demonstrate consistency under any set of effects $u$ for which $\frac{d}{d\gamma}\mathbb{E}[\Delta(\gamma)|Z]$ can be bounded away from zero in some neighborhood of $\mathring{\gamma}$.

##### S3.2.5 Asymptotic variance of the stationarity function

Split the stationarity functions into two components:

$$\Delta(\gamma) = \underbrace{\frac{\sigma_e^{-2} y^T P_\gamma \tilde{Z}\tilde{Z}^T P_\gamma y}{\text{tr}\,[P_\gamma \tilde{Z}\tilde{Z}^T]}}_{:=\text{LHS}(\gamma)} - \underbrace{\frac{\sigma_e^{-2} y^T P_\gamma^2 y}{\text{tr}\,[P_\gamma]}}_{:=\text{RHS}(\gamma)}.$$

Consider the conditional variance of the stationarity function $\Delta(\gamma)$ given $Z$:

$$\begin{aligned} Var(\Delta(\gamma)|Z) =& \mathbb{E}[\Delta(\gamma)^2|Z] - \mathbb{E}[\Delta(\gamma)|Z]^2 \\ =& \big(\mathbb{E}[\text{LHS}(\gamma)^2|Z] + \mathbb{E}[\text{RHS}(\gamma)^2|Z] - 2\mathbb{E}[\text{LHS}(\gamma)\text{RHS}(\gamma)|Z]\big) - \mathbb{E}[\Delta(\gamma)|Z]^2. \end{aligned}$$

To compute the above, we make use of the following lemma regarding the products of quadratic forms:

**Lemma 15.** *Let $x \sim \mathcal{MVN}(0, I\sigma^2)$ and let $A, B$ denote symmetric real matrices conformable with $x$. Then*

$$\mathbb{E}[x^T Ax \cdot x^T Bx] = \text{tr}\,[A]\text{tr}\,[B] + 2\text{tr}\,[AB].$$

*Proof.* This is an immediate corollary of Lemma 2 in (Bao *et al.*, 2010). □

Now, computing the above terms comprising $Var(\Delta(\gamma)|Z)$ individually, we have

$$\sigma_e^{-2} u^T \left( \frac{\Sigma_\gamma^{-1}\zeta\zeta^T\Sigma_\gamma^{-1}\zeta\zeta^T}{\text{tr}\,[\Sigma_\gamma^{-1}\zeta\zeta^T]} - \frac{\Sigma_\gamma^{-2}\zeta\zeta^T}{\text{tr}\,[\Sigma_\gamma^{-1}]} \right) u + \left( \frac{\text{tr}\,[\Sigma_\gamma^{-1}\zeta\zeta^T\Sigma_\gamma^{-1}]}{\text{tr}\,[\Sigma_\gamma^{-1}\zeta\zeta^T]} - \frac{\text{tr}\,[\Sigma_\gamma^{-2}]}{\text{tr}\,[\Sigma_\gamma^{-1}]} \right)$$

First noting that

$$\begin{aligned} \mathbb{E}[\text{LHS}(\gamma)|Z]^2 &= \sigma_e^{-4} \left( \frac{u^T\Sigma_\gamma^{-1}\zeta\zeta^T\Sigma_\gamma^{-1}\zeta\zeta^T u + \text{tr}\,[\Sigma_\gamma^{-1}\zeta\zeta^T\Sigma_\gamma^{-1}]}{\text{tr}\,[\Sigma_\gamma^{-1}\zeta\zeta^T]} \right)^2 \\ &= \frac{\text{tr}\,[\zeta^T\Sigma_\gamma^{-1}\zeta\zeta^T\Sigma_\gamma^{-1}\zeta u u^T]^2 + \text{tr}\,[\Sigma_\gamma^{-1}\zeta\zeta^T\Sigma_\gamma^{-1}]^2}{\sigma_e^4 \text{tr}\,[\Sigma_\gamma^{-1}\zeta\zeta^T]} \\ &\quad + \frac{2\text{tr}\,[\zeta^T\Sigma_\gamma^{-1}\zeta\zeta^T\Sigma_\gamma^{-1}\zeta u u^T]\text{tr}\,[\Sigma_\gamma^{-1}\zeta\zeta^T\Sigma_\gamma^{-1}]}{\sigma_e^4 \text{tr}\,[\Sigma_\gamma^{-1}\zeta\zeta^T]}, \end{aligned}$$

we see that

$$
\begin{aligned}
\mathbb{E}[\mathrm{LHS}(\gamma)^2|Z] =& \sigma_e^{-4}\mathbb{E}\left[\left(\frac{u^T\zeta^T\Sigma_\gamma^{-1}\zeta\zeta^T\Sigma_\gamma^{-1}\zeta u + e^T P_{\hat{\gamma}}\tilde{Z}\tilde{Z}^T P_{\hat{\gamma}} e}{\mathrm{tr}\,[P_\gamma \tilde{Z}\tilde{Z}^T]}\right)^2 |Z\right] \\
=& \frac{(u^T\zeta^T\Sigma_\gamma^{-1}\zeta\zeta^T\Sigma_\gamma^{-1}\zeta u)^2 + \mathbb{E}[(e^T P_{\hat{\gamma}}\tilde{Z}\tilde{Z}^T P_{\hat{\gamma}} e)^2|Z]}{\sigma_e^4 \mathrm{tr}\,[P_\gamma \tilde{Z}\tilde{Z}^T]} \\
&+ \frac{+2(u^T\zeta^T\Sigma_\gamma^{-1}\zeta\zeta^T\Sigma_\gamma^{-1}\zeta u)\mathbb{E}[e^T P_{\hat{\gamma}}\tilde{Z}\tilde{Z}^T P_{\hat{\gamma}} e|Z]}{\sigma_e^4 \mathrm{tr}\,[P_\gamma \tilde{Z}\tilde{Z}^T]} \\
=& \frac{\mathrm{tr}\,[\zeta^T\Sigma_\gamma^{-1}\zeta\zeta^T\Sigma_\gamma^{-1}\zeta uu^T]^2 + \mathrm{tr}\,[\Sigma_\gamma^{-1}\zeta\zeta^T\Sigma_\gamma^{-1}]^2 + 2\mathrm{tr}\,[(\Sigma_\gamma^{-1}\zeta\zeta^T\Sigma_\gamma^{-1})^2]}{\sigma_e^4 \mathrm{tr}\,[\Sigma_\gamma^{-1}\zeta\zeta^T]^2} \\
&+ \frac{2\mathrm{tr}\,[\zeta^T\Sigma_\gamma^{-1}\zeta\zeta^T\Sigma_\gamma^{-1}\zeta uu^T]\mathrm{tr}\,[\Sigma_\gamma^{-1}\zeta\zeta^T\Sigma_\gamma^{-1}\zeta\zeta^T]}{\sigma_e^4 \mathrm{tr}\,[\Sigma_\gamma^{-1}\zeta\zeta^T]^2} \\
=& \mathbb{E}[\mathrm{LHS}(\gamma)|Z]^2 + \frac{2\mathrm{tr}\,[\Sigma_\gamma^{-2}\zeta\zeta^T\Sigma_\gamma^{-2}\zeta\zeta^T]}{\sigma_e^4 \mathrm{tr}\,[\Sigma_\gamma^{-1}\zeta\zeta^T]^2}.
\end{aligned}
$$

Similarly, we have that

$$
\mathbb{E}[\mathrm{RHS}(\gamma)^2|Z] = \mathbb{E}[\mathrm{RHS}(\gamma)|Z]^2 + \frac{2\mathrm{tr}\,[\Sigma_\gamma^{-4}]}{\sigma_e^4 \mathrm{tr}\,[\Sigma_\gamma^{-1}]^2}.
$$

Further, noting that,

$$
\mathbb{E}[\mathrm{LHS}(\gamma)|Z]\cdot\mathbb{E}[\mathrm{RHS}(\gamma)|Z] = \frac{\left(\mathrm{tr}\,[\zeta^T\Sigma_\gamma^{-1}\zeta\zeta^T\Sigma_\gamma^{-1}\zeta uu^T] + \mathrm{tr}\,[\Sigma_\gamma^{-2}\zeta\zeta^T]\right)\left(\mathrm{tr}\,[\zeta^T\Sigma_\gamma^{-2}\zeta uu^T] + \mathrm{tr}\,[\Sigma_\gamma^{-2}]\right)}{\sigma_e^4 \mathrm{tr}\,[\Sigma_\gamma^{-1}\zeta\zeta^T]\mathrm{tr}\,[\Sigma_\gamma^{-1}]},
$$

we can write

$$
\mathbb{E}[\mathrm{LHS}(\gamma)\mathrm{RHS}(\gamma)|Z] = \mathbb{E}[\mathrm{LHS}(\gamma)|Z]\cdot\mathbb{E}[\mathrm{RHS}(\gamma)|Z] + \frac{2\mathrm{tr}\,[\Sigma_\gamma^{-4}\zeta\zeta^T]}{\sigma_e^4 \mathrm{tr}\,[\Sigma_\gamma^{-1}\zeta\zeta^T]\mathrm{tr}\,[\Sigma_\gamma^{-1}]}.
$$

All together, we have

$$
\begin{aligned}
\mathbb{E}[\mathrm{LHS}(\gamma)^2|Z] =& \mathbb{E}[\mathrm{LHS}(\gamma)|Z]^2 + \frac{2\mathrm{tr}\,[(\Sigma_\gamma^{-2}\zeta\zeta^T\Sigma_\gamma^{-2}\zeta\zeta^T]}{\sigma_e^4 \mathrm{tr}\,[\Sigma_\gamma^{-1}\zeta\zeta^T]^2}, \\
\mathbb{E}[\mathrm{RHS}(\gamma)^2|Z] =& \mathbb{E}[\mathrm{RHS}(\gamma)|Z]^2 + \frac{2\mathrm{tr}\,[\Sigma_\gamma^{-4}]}{\sigma_e^4 \mathrm{tr}\,[\Sigma_\gamma^{-1}]^2}, \\
\mathbb{E}[\mathrm{LHS}(\gamma)\mathrm{RHS}(\gamma)|Z] =& \mathbb{E}[\mathrm{LHS}(\gamma)|Z]\cdot\mathbb{E}[\mathrm{RHS}(\gamma)|Z] + \frac{2\mathrm{tr}\,[\Sigma_\gamma^{-4}\zeta\zeta^T]}{\sigma_e^4 \mathrm{tr}\,[\Sigma_\gamma^{-1}\zeta\zeta^T]\mathrm{tr}\,[\Sigma_\gamma^{-1}]}, \\
\mathbb{E}[\Delta(\gamma)|Z]^2 =& \sigma_e^4\left(\mathbb{E}[\mathrm{RHS}(\gamma)|Z]^2 + \mathbb{E}[\mathrm{LHS}(\gamma)|Z]^2\right. \\
& \left. -2\mathbb{E}[\mathrm{LHS}(\gamma)|Z]\cdot\mathbb{E}[\mathrm{RHS}(\gamma)|Z]\right).
\end{aligned}
$$

Canceling terms, we thus have

$$
\frac{1}{2\sigma_e^4}Var(\Delta(\gamma)|Z) = \mathbb{E}[\mathrm{LHS}(\gamma)^2|Z] + \mathbb{E}[\mathrm{RHS}(\gamma)^2|Z] - 2\mathbb{E}[\mathrm{LHS}(\gamma)\mathrm{RHS}(\gamma)|Z] - \mathbb{E}[\Delta(\gamma)|Z]^2.
$$

$$= \frac{\text{tr}\,[\Sigma_\gamma^{-2}\zeta\zeta^T\Sigma_\gamma^{-2}\zeta\zeta^T]}{\text{tr}\,[\Sigma_\gamma^{-1}\zeta\zeta^T]^2} + \frac{\text{tr}\,[\Sigma_\gamma^{-4}]}{\text{tr}\,[\Sigma_\gamma^{-1}]^2} - 2\frac{\text{tr}\,[\Sigma_\gamma^{-4}\zeta\zeta^T]}{\text{tr}\,[\Sigma_\gamma^{-1}\zeta\zeta^T]\text{tr}\,[\Sigma_\gamma^{-1}]}.$$

We will show that these terms vanish with high probability. First note that

$$\frac{\text{tr}\,[\Sigma_\gamma^{-2}\zeta\zeta^T\Sigma_\gamma^{-2}\zeta\zeta^T]}{\text{tr}\,[\Sigma_\gamma^{-1}\zeta\zeta^T]^2} = \frac{(n-c)^{-2}\text{tr}\,[\Sigma_\gamma^{-2}\zeta\zeta^T\Sigma_\gamma^{-2}\zeta\zeta^T]}{(n-c)^{-2}\text{tr}\,[\Sigma_\gamma^{-1}\zeta\zeta^T]^2}.$$

The trace term in the numerator is a continuous, bounded spectral function of the eigenvalues of $\zeta\zeta^T$:

$$\begin{aligned} \text{tr}\,[\Sigma_\gamma^{-2}\zeta\zeta^T\Sigma_\gamma^{-2}\zeta\zeta^T] &= \sum \lambda_i \psi_{(2,4)}(\lambda_i) \\ \implies (n-c)^{-2}\text{tr}\,[\Sigma_\gamma^{-2}\zeta\zeta^T\Sigma_\gamma^{-2}\zeta\zeta^T] &= (n-c)^{-1}\left((n-c)^{-1}\text{tr}\,[\Sigma_\gamma^{-2}\zeta\zeta^T\Sigma_\gamma^{-2}\zeta\zeta^T]\right) \\ &\xrightarrow{P} (n-c)^{-1}\Psi_{(2,4)}. \end{aligned}$$

On the other hand, we've already seen that $(n-c)^{-1}\text{tr}\,[\Sigma_\gamma^{-1}\zeta\zeta^T] \xrightarrow{P} \Psi_{(1,1)}$. Again applying Slutzky's theorem, we have

$$\frac{\text{tr}\,[\Sigma_\gamma^{-2}\zeta\zeta^T\Sigma_\gamma^{-2}\zeta\zeta^T]}{\text{tr}\,[\Sigma_\gamma^{-1}\zeta\zeta^T]^2} \xrightarrow{P} (n-c)^{-1}\frac{\Psi_{(2,4)}}{\Psi_{(1,1)}^2} = o_p(1).$$

By a similar argument, we have that

$$\begin{aligned} \frac{\text{tr}\,[\Sigma_\gamma^{-4}]}{\text{tr}\,[\Sigma_\gamma^{-1}]^2} &\xrightarrow{P} (n-c)^{-1}\frac{\Psi_{(0,4)}}{\Psi_{(0,1)}^2} = o_p(1), \\ \frac{\text{tr}\,[\Sigma_\gamma^{-4}\zeta\zeta^T]}{\text{tr}\,[\Sigma_\gamma^{-1}\zeta\zeta^T]\text{tr}\,[\Sigma_\gamma^{-1}]} &\xrightarrow{P} (n-c)^{-1}\frac{\Psi_{(1,4)}}{\Psi_{(1,1)}\Psi_{(0,1)}} = o_p(1), \end{aligned}$$

thus yielding $Var(\Delta(\gamma)|Z) \xrightarrow{P} 0$. Now, applying the conditional form of Chebyshev's inequality, we have $\forall\epsilon > 0$,

$$P\left(|\Delta(\gamma) - \mathbb{E}[\Delta(\gamma)|Z]| \geq \varepsilon|Z\right) \leq \varepsilon^{-2}Var(\Delta(\gamma)|Z) \xrightarrow{P} 0,$$

i.e., that $\Delta(\gamma) - \mathbb{E}[\Delta(\gamma)|Z] \xrightarrow{P} 0$. Together with the result of the previous section ($|\mathbb{E}[\Delta(\gamma)|Z]| \xrightarrow{P} h(\gamma)$ with $h(\gamma) = 0$ iff $\gamma = \mathring{\gamma}$), this implies that $\hat{\gamma} \xrightarrow{P} \mathring{\gamma}$.

### A Supplemental figures

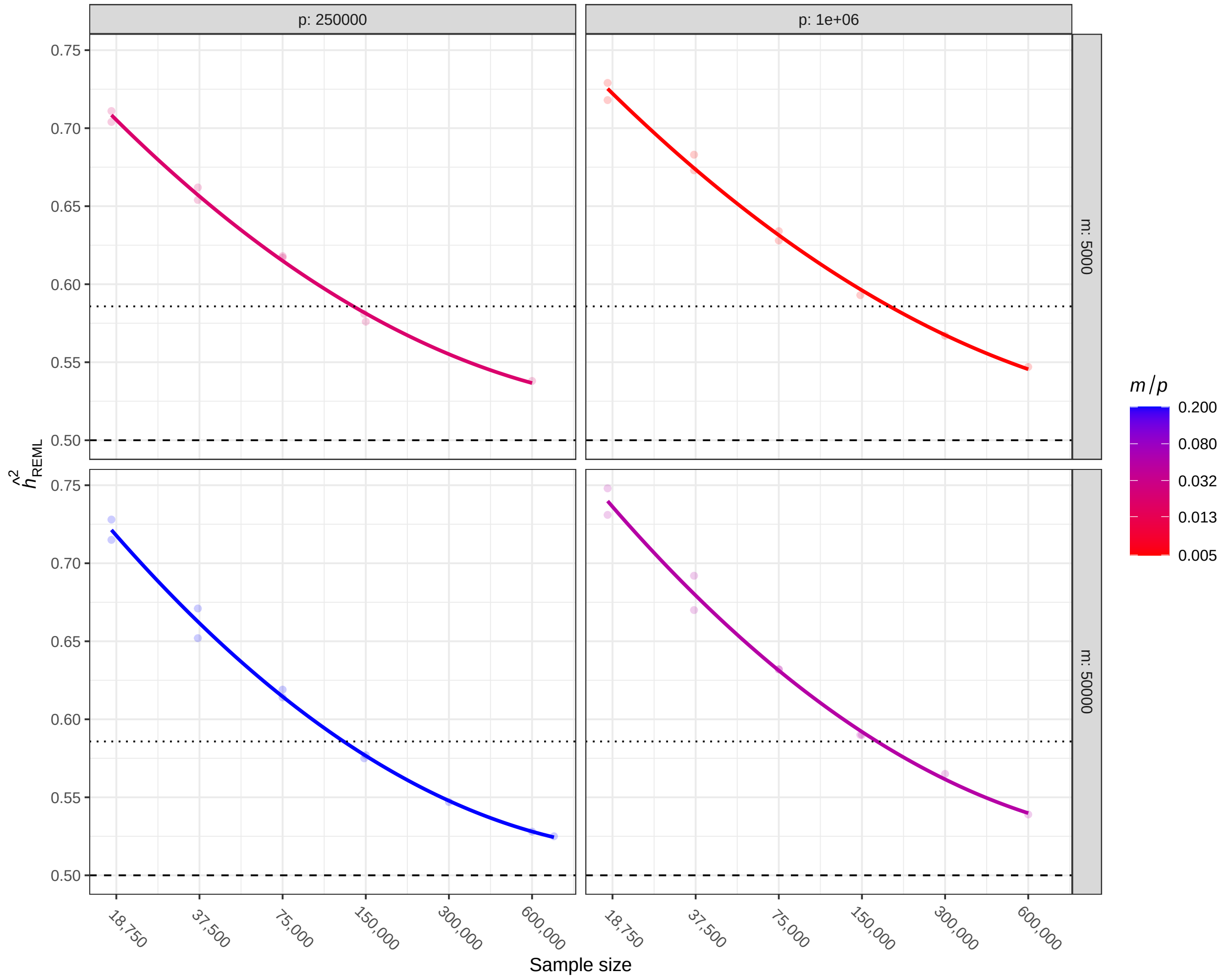

Figure S1: $\hat{h}^2_{\text{REML}}$ as a function of sample size ($n$), the total number of SNPs included in the model ($p$), and the number of causal variants ($m$) in synthetic data. We observe convergence of $\hat{h}^2_{\text{REML}}$ toward $h_0^2$ for all values of $m/p$.
